# Strain, procedures, and tools for reproducible genetic transformation and genome editing of the emerging plant model *Spirodela polyrhiza* (L.) Schleid

**DOI:** 10.1101/2025.08.28.672806

**Authors:** Verónica Barragán-Borrero, Amanda de Santana Lopes, Enrico Diniz Rodrigues Batista, Martin Höfer, Rana Elias, Abhisek Chakraborty, Arturo Ponce-Mañe, Clotilde Descombes, Laura Diezma-Navas, Lydia Petraki, Meret Huber, Shuqing Xu, Arturo Marí-Ordóñez

## Abstract

Duckweeds (*Lemnaceae*) have excellent potential for fundamental and applied research due to ease of cultivation, small size, and continuous fast clonal growth. However, their usage as model organisms and platforms for biotechnological applications is often limited by the lack of universal genetic manipulation methods necessary for transgene expression, gene editing, and other methods to modify gene expression. To identify suitable strains for genetic manipulation of the giant duckweed, *Spirodela polyrhiza,* we screened several genotypes for callus induction and regeneration and established genetic transformation. We have identified *SP162* to be amenable to *Agrobacterium*-mediated transformation via tissue culture. The procedure is robust and reproducible across laboratories, allowing stable expression of different reporter genes and selectable markers, enabling CRISPR/Cas9-mediated genome editing. In addition, due to a weak small RNA-based silencing response, *S. polyrhiza* sustains prolonged periods of transgene activity in transient expression assays. To promote duckweed research and encourage the adoption of *S. polyrhiza*, we have made *SP162* (ID#: *5676*) and its genome publicly available and provide here detailed procedures for its cultivation and transformation. Furthermore, we created a web server to explore its genome, retrieve gene sequences, and implemented orthologous gene search and a gRNA design function for diverse CRISPR/Cas-based applications (https://agxu.uni-mainz.de/SP162/).

## Introduction

Despite the vast diversity of land plants, most mechanistic understanding of plant gene function stems from few model systems with well-established genetic engineering protocols, such as *Arabidopsis thaliana*, *Oryza sativa* (rice), and *Marchantia polymorpha*. Research with these models has advanced plant biology tremendously yet revealed only a narrow slice of the evolutionary, ecological, and physiological diversity across the plant kingdom. Expanding the repertoire of genetically tractable species is therefore essential, not only to uncover lineage-specific adaptations, but also to develop sustainable applications capitalizing on the unique traits of underexplored plants.

Among these are duckweeds, floating aquatic plants in the family *Lemnaceae* comprising 36 species across five genera (Tippery et al. 2021). As the smallest flowering plants, they develop a simple body with 1-10 mm leaf-like fronds and hairless adventitious roots, with some genera having completely lost roots and vasculature (Acosta et al. 2021; Ziegler et al. 2023). Their exceptionally rapid growth, ease of cultivation, and efficient uptake of labeled compounds made duckweeds attractive model organisms for plant physiology and ecology in the mid-20th century (Hillman 1961; Acosta et al. 2021). However, interest declined in the 1990s with the rise of more genetically tractable models, like *A. thaliana*.

Recently, duckweed research has resurged, driven by advances in omics, renewed interest in plant ecological diversity, and potential applications in environmental and agricultural biotechnology (Acosta et al. 2021; Yang et al. 2021). Their small size and clonal propagation make them especially suitable for controlled multigenerational experiments, crucial for studying transgenerational plasticity, epigenetic memory, and stress responses (Huber et al. 2021; Antro et al. 2023; Chávez et al. 2025). Duckweeds are also emerging as key contributors to circular bioeconomy, through applications such as wastewater nutrient recycling (Cheng and Stomp 2009), phytoremediation (Ekperusi et al. 2019), carbon sequestration, and bio-based protein and starch production for food, feed, and fuels (Cui and Cheng 2015; Appenroth et al. 2017; 2021).

Among duckweeds, *Spirodela polyrhiza* stands out as a promising model. It possesses the smallest known genome in its family (140–158 Mb) (Wang et al. 2014; Harkess et al. 2021), and, despite its name “giant duckweed”, still offers rapid clonal growth and large fronds for experimental manipulation. Its simple development, high phenotypic plasticity, and stress tolerance, make it valuable for fundamental molecular, developmental, evolutionary and ecological studies as well as applied research (Huber et al. 2021; Baggs et al. 2022; Ware et al. 2023; Sun et al. 2023; Turcotte et al. 2024; Wang et al. 2024; Malacrinò et al. 2024; Smith et al. 2024; Chávez et al. 2025; Böttner et al. 2025).

Despite these desirable characteristics, absence of a reliable, reproducible protocol for stable genetic transformation limits the exploitation of *S. polyrhiza* as a model. In most monocotyledonous plants, stable transformation is often hindered by challenges in tissue culture, regeneration and recalcitrance to *Agrobacterium*-mediated transformation as they are not natural hosts of *A. tumefaciens*. Thus, successful establishment of transformation procedures require screening and selection of suitable and compatible plant genotypes and bacterial strains together with method optimization (Sood et al. 2011). While transient gene expression via carbon nanotubes (Islam et al. 2024) and *Agrobacterium*-mediated DNA delivery (Peterson et al. 2021; Dombey et al. 2025) have been reported in *S. polyrhiza*, only one publication described stable transformation (Yang et al. 2018). However, the study in a not publicly available genotype lacked adequate genotypic and phenotypic characterization, not allowing to reproduce the findings.

To complement the toolbox for duckweed research, we developed a detailed, reproducible *Agrobacterium*- and tissue culture-based transformation protocol for a publicly available *S. polyrhiza* genotype. We generated stable transgenic lines expressing various reporter genes and selectable markers and demonstrated CRISPR/Cas9-mediated genome editing. This know-how will support the growing duckweed research community and help unlock the potential of *S. polyrhiza* for fundamental discoveries and sustainable biotechnological applications.

## Results

### *S. polyrhiza* genotypes vary for callus induction rates

Success rates in transformation protocols including tissue culture steps vary between strains and genotypes of the same species depending on the ability to form callus (Abe and Futsuhara 1986; Nguyen et al. 2020; Lardon et al. 2020). Therefore, we screened a collection of *S. polyrhiza* genotypes (Wang et al. 2024) for strains displaying high callus induction rates. We first established a suitable callus induction procedure by testing three published protocols for *S. polyrhiza* (Wang 2016; Huang et al. 2018; Yang et al. 2018), and one for the related species *Lemna aequinoctialis* (Liu et al. 2019). The protocols, differing in the type of explants, composition of phytohormones, and their concentration (**Figure 1a**), were tested on axenic cultures (**Method S1, S2**) of four *S. polyrhiza* genotypes from diverse geographical origin. The methods based on wound-induced callus formation (Wang 2016) and root-derived calli from dorsoventrally laid fronds (Huang et al. 2018) had the highest callus induction rates (**Figure 1b**) and produced calli from frond cuttings or root meristems as expected from either protocol (**Figure 1c**).

**Figure 1:**
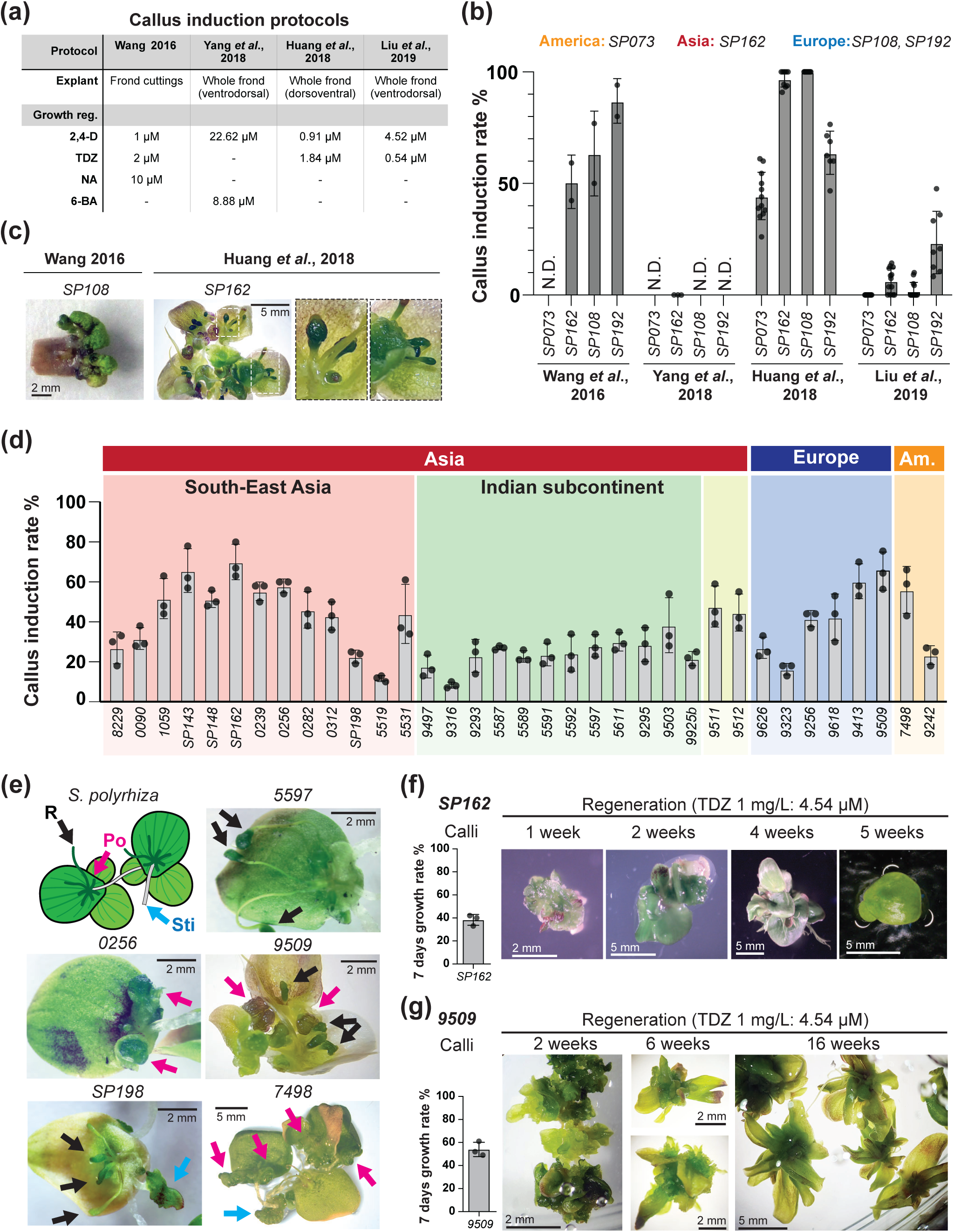
Identification of compatible procedures and genotypes for *S. polyrhiza* callus induction and frond regeneration. **(a)** Types of explants and concentrations of phytohormones (growth regulators) used in previously published protocols for callus induction tested here. **(b)** Mean callus induction rate (as % of explants initiating callus formation) in four different *S. polyrhiza* genotypes, from different origins indicated above, using the protocols in (a). Dots display individual values from independent replicates. Error bars represent standard deviation of the mean. N.D.: not determined. **(c)** Pictures of callus induced from cutting explants or roots according to the indicated published protocols. **(d)** Mean callus induction rate in % of 35 *S. polyrhiza* genotypes from different geographic origin using the Huang *et al*. callus induction protocol. Dots display individual values from three independent replicates. Error bars represent standard deviation of the mean. **(e)** Schematic representation of *S. polyrhiza* frond structure from a ventral position and pictures of different genotypes with ongoing callus formation from different tissues using Huang *et al*. callus induction protocol. R: roots (black arrow), Po: budding pockets (magenta arrow), Sti: stipe, stalk-like connecting structure between mother and daughter fronds (blue arrow). **(f)** *S. polyrhiza SP162* calli growth rate in % (as increase of biomass) (dots display individual values from three independent replicates. Error bars represent standard deviation of the mean) and representative images during regeneration following Huang *et al*. protocol with increased Thidiazuron (TDZ) to 1 mg/L. **(g)** Same as (f) but for *S. polyrhiza 9509*.

The less laborious procedure with root-derived calli was chosen to expand the screen to 35 *S. polyrhiza* genotypes. Callus induction rates varied across genotypes, with some South-East Asian and European genotypes displaying mean callus induction rates above 60% (**Figure 1d**). We also observed variability between genotypes for the tissue-of-origin of the calli: in addition to the calli initiated from root meristems (R), some genotypes formed calli also from vegetative budding pockets (Po), where shoot meristems are located, or from stipes (Sti), the stalk-like structure connecting mother-to-daughter fronds (**Figure 1e**). Hence, *S. polyrhiza* displays variation across genotypes in callus induction responses, both in initiation rate and tissue-of-origin.

### The *S. polyrhiza SP162* genotype can be regenerated into fronds

High rates of callus induction does not necessarily result in successful plant regeneration, two independent and often uncorrelated processes (Abe and Futsuhara 1986; Ikeuchi et al. 2013; Motte et al. 2014; Ikeuchi et al. 2019). Therefore, we examined regeneration in two genotypes with high callus induction, the Asian *SP162* and European *9509* (**Figure 1d**). *9509* offers extensive genomic resources, including an assembled reference genome, transcriptomes and epigenomes (Hoang et al. 2018; Harkess et al. 2021; 2024; Dombey et al. 2025). *SP162*, despite lacking such resources, exhibited good formation of turions, sinking dormant vegetative propagules (Landolt 1986; Appenroth 2002; Ziegler et al. 2023; Kim et al. 2024), with near 100% germination after months of cold storage (**Figure S1, Method S3**), providing a convenient storage solution. In contrast, *9509* lacked turion production in our hands.

Despite Huang and colleagues had shown that *S. polyrhiza* regeneration is best induced at low concentrations of the synthetic cytokinin thidiazuron (TDZ) (0.2 mg/L, 0.9 µM), or in absence of any cytokinin (Huang et al. 2018), neither genotype initiated regeneration under these conditions. Others had reported increasing regeneration efficiencies at higher TDZ concentrations (1 mg/L, 4.54 µM) (Wang 2016; Yang et al. 2018). Indeed, both genotypes initiated regeneration in the presence of 1mg/L TDZ (**Figure 1f,g**). However, complete regeneration of individual fronds was only achieved in *SP162* (**Figure 1f**), whereas *9509* calli produced only elongated frond-like structures but not fully regenerated fronds (**Figure 1f**). Therefore, *SP162* was selected as the genotype for further studies.

### Callus induction and maintenance under low light improves *SP162* callus growth and proliferation

Although *SP162* calli did regenerate, they were prone to browning despite the presence of anti-browning agents, polyvinylpyrrolidone (PVP) and citric acid, in the callus induction medium (CIM) (**Figure 2a, Method S1**) and grew slowly (**Figure 1f**), two factors negatively affecting *Agrobacterium*-mediated transformation and regeneration (Potrykus 1990; Sangwan et al. 1992; Greenwood and Glaus 2022; Liu et al. 2024). Browning of *S. polyrhiza* calli can be prevented in darkness, although this results in decreased proliferation (Huang et al. 2018). Therefore, we tested whether low light conditions could represent a compromise to reduce browning without limiting growth and proliferation. We performed *SP162* callus induction on CIM at 21°C and covered the culture plates with two sheets of white paper (80 gr/m^2^), bringing light intensity during the 16 h light period to 20 µmol/m^2^/s (**Figure 2b, Method S4**). Swelling of young roots was already apparent after a week (**Figure S2**). By the fourth week, 2-3 mm wide dark-green calli had formed at root tips and budding pockets (**Figure 2c, S2**). Newly formed calli under low light showed little or no browning (**Figure 2c, S2**). Hence, reducing light intensity had a beneficial impact on callus induction by producing healthier calli.

**Figure 2:**
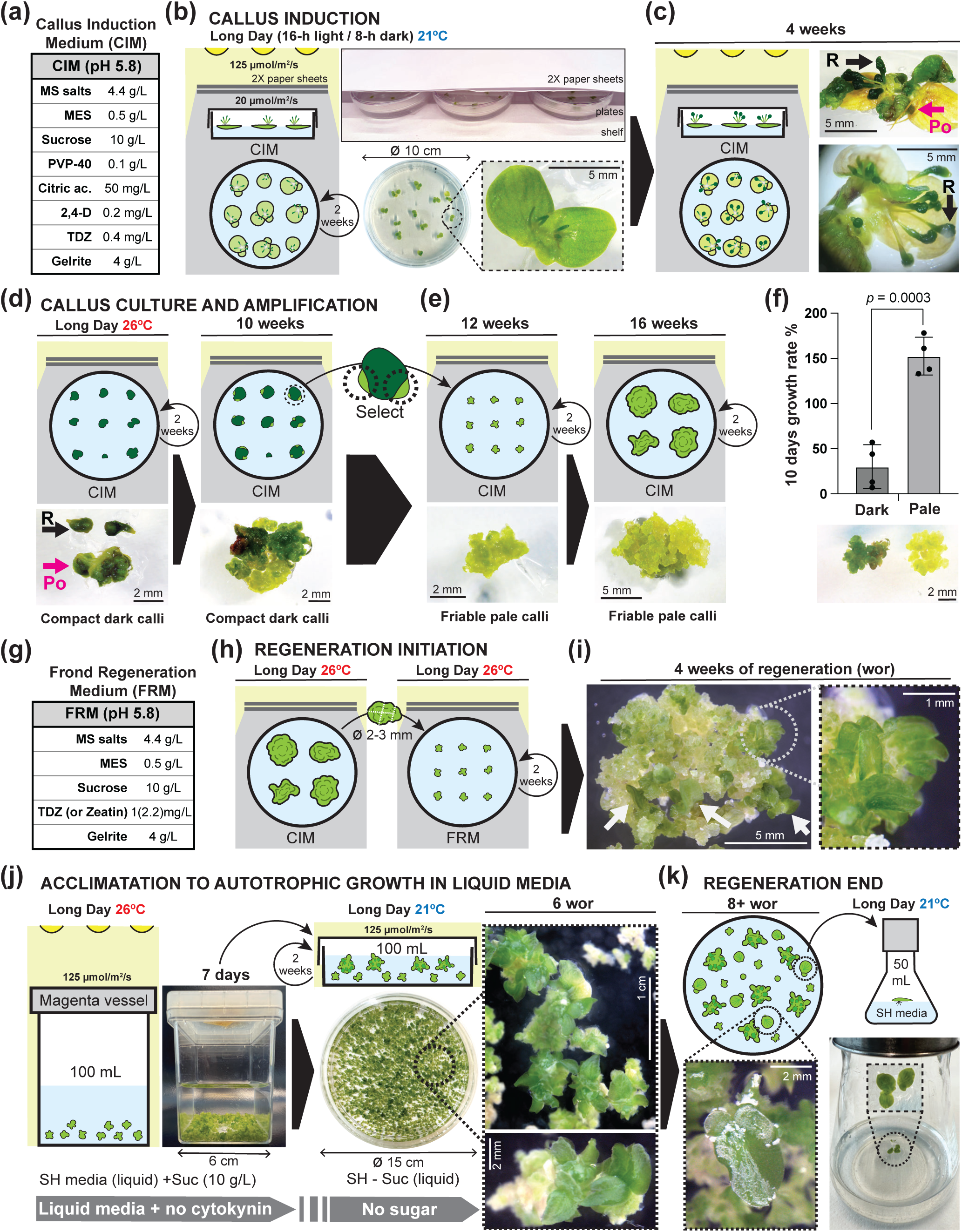
*S. polyrhiza SP162* callus induction and frond regeneration under low light. **(a)** Composition of the *SP162* callus induction medium (CIM). **(b)** Schematics of *SP162* callus induction conditions under low light and images of experimental setup and fronds in dorsoventral position on CIM. **(c)** *SP162* calli induced from either roots (R) or budding pockets (Po) after four weeks in CIM under low light. **(d)** Images of initial calli transferred to CIM and 10 weeks after callus induction starting to develop pale sectors cultured on CIM under low light. **(e)** Images of selected and amplified pale friable calli sectors grown under low light. **(f)** Comparative growth rate in % (as increase of biomass) of dark-green calli cultured under high light and pale-green calli cultured under low light. Dots display individual values from three independent replicates. Error bars represent standard deviation of the mean. *p* indicates two-tailed P-value of unpaired t-test between the two groups. **(g)** Composition of the *SP162* frond regeneration medium (FRM) medium. **(h)** Schematic of the transfer procedure from CIM to FRM to initiate regeneration under low light. **(i)** Image of early regenerating fronds (marked with white arrow) after 4 weeks of regeneration (wor) on solid FRM. **(j)** Schematics and images of the two-step acclimation for frond regeneration in liquid media together with pictures of regenerating fronds after removal of cytokinin and sugar after 6 wor. **(k)** Images of a regenerated frond in SH-medium in culture plate and proliferation after individualization.

After four weeks, calli were detached and subcultured under low light conditions, but temperature was increased to 26°C to promote faster growth (**Figure 2d**). About six weeks later, pale-green sectors started to appear. These were less compact, more friable (**Figure 2d**), and seemed to proliferate faster. They were isolated, subcultured and amplified (**Figure 2e, Method S4**). Even after several months of culture under low light, pale-green calli did not show browning and grew five to six times faster than dark-green calli cultured without shading (**Figure 2f**).

### *SP162* calli can regenerate through a two-step acclimatation process under low light

Given the positive impact of low light on callus health and growth, we tested whether *SP162* pale calli could regenerate under low light on frond regeneration medium (FRM) containing either 1 mg/L TDZ (**Figure 1f**), or zeatin, another cytokinin shown to induce high levels of regeneration in *S. polyrhiza* at 2.2 mg/L (10 µM) (Wang 2016) (**Figure 2g, Method S1**). Callus fragments were cultured on solid FRM with either cytokinin under callus induction and maintenance conditions (**Figure 2h, Method S5**). After four weeks, regeneration initiated, with small dark-green frond-like structures appearing, singly or in clusters, in both TDZ or zeatin containing FRM (**Figure 2i**, **Figure S3**).

To progressively acclimate regenerating fronds to autotrophic growth in standard liquid culture conditions, we applied a stepwise process. First, calli with regenerating fronds were accustomed to liquid media and increased light intensity (125 µmol/m^2^/s) by culturing them in Magenta vessels in 5 cm-deep liquid Schenk and Hildebrant (SH)-medium with sucrose but without growth regulators for seven days. Afterwards, they were moved to 1 cm shallow SH-medium without sugar in culture dishes to improve gas exchange and light exposure at 21°C (**Figure 2j, Method S1, S5**).

Around six weeks after induction, regenerating fronds visible to the naked eye provided increased buoyancy and calli emerged to the surface (**Figure 2j, S4a**). Two weeks later, individual fronds displaying a vitrified aspect (**Figure S4a**) started to appear on the surface of the culture and were regularly transferred to individual flasks (**Figure 2k, Method S5**). Frond clusters continued to appear up to twelve weeks after induction (**Figure S4b**). Thus, we established culture conditions enabling healthy and fast-growing calli of *SP162* to regenerate into fronds.

### *SP162* calli are sensitive to several selective antibiotics

Crucial for the successful identification and amplification of transgenic cells is the use of selectable markers to eliminate or arrest the proliferation of non-transgenic cells. To identify suitable selective agents for *SP162* calli, we tested their sensitivity to different concentrations of four broadly used antibiotics in plant selection: kanamycin, hygromycin, G418/geneticin, and spectinomycin. *SP162* calli were sensitive to all four antibiotics, but to different degrees. While calli showed lower sensitivity to kanamycin, they displayed high sensitivity to the other three antibiotics, with spectinomycin and hygromycin being the most efficient at a broad range of concentrations (**Figure 3, S5**). Given that hygromycin had the fastest impact on calli survival at low concentrations (**Figure 3, S5**), hygromycin resistance was chosen as selectable marker for testing transformation in *SP162*.

**Figure 3:**
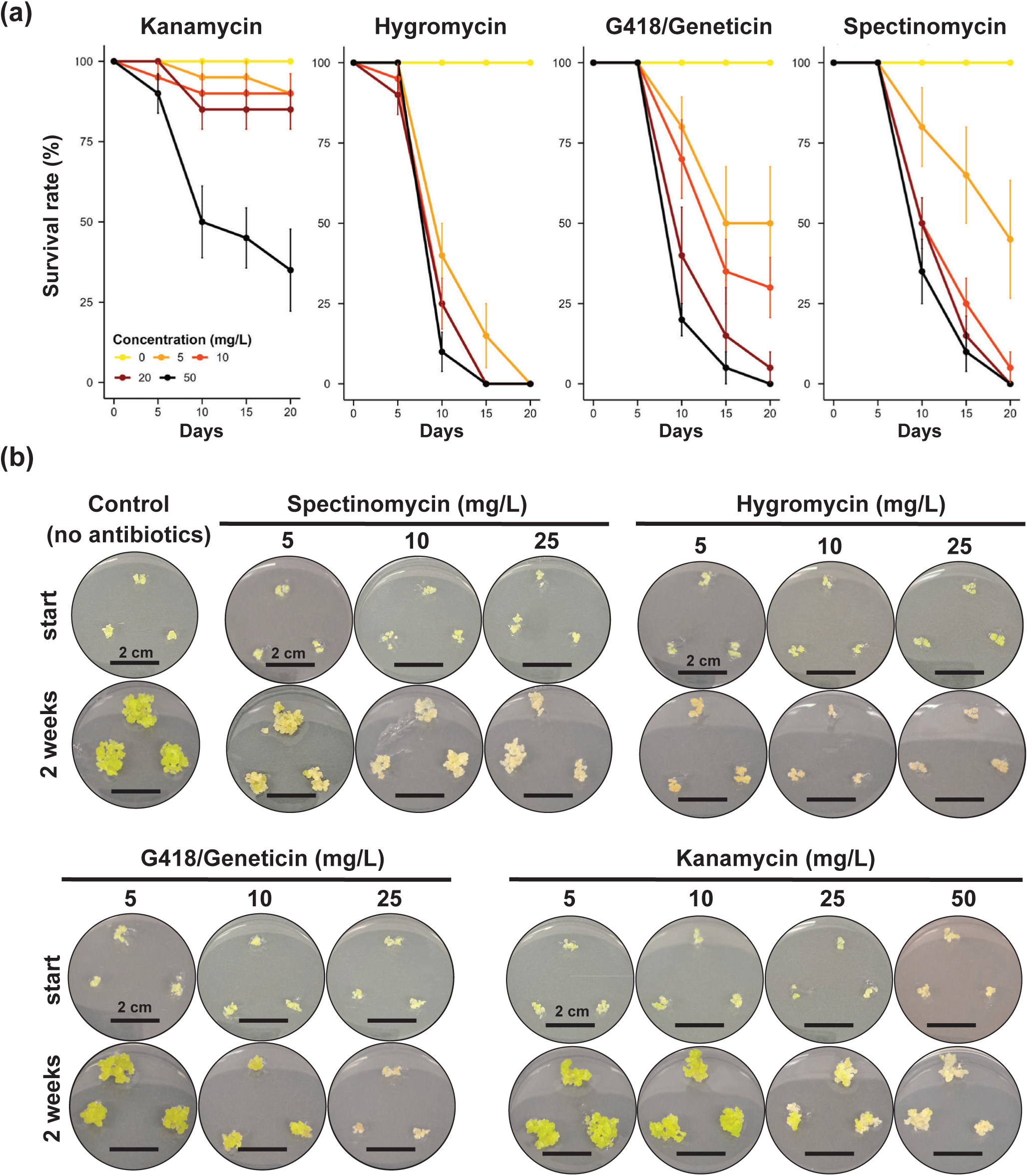
Sensitivity of *S. polyrhiza SP162* calli to antibiotics. **(a)** Survival rates (%) of calli exposed to different concentrations of the indicated antibiotics along a 20-day period. Dots indicate average survival rates and error bars represent standard deviation of the mean of 5 replicates. Survival was assessed as absence of chlorosis, browning or impact on growth. **(b)** Representative images of calli exposed to different concentrations of indicated antibiotics in CIM media after transfer to media with antibiotics, or without as control, and two weeks later. See Figure S5 for a broader range of concentrations.

### *SP162* calli are suitable for stable *Agrobacterium*-mediated transformation

To assess the feasibility of *Agrobacterium-*mediated transformation in *SP162*, we aimed at transforming calli with green fluorescent protein (*GFP)* driven by the maize *UBIQUITIN* promoter (*pZmUBQ::eGFP*), suitable for strong ubiquitous expression in monocots (Christensen and Quail 1996), and hygromycin resistance (*HptII*) under the *NOPALINE SYNTHASE* promoter *(pNOS)* as a selectable marker. The hypervirulent *Agrobacterium tumefaciens EHA105* strain was chosen given it’s increased general infectivity (Hood et al. 1993), which has improved transformation in other monocots (Yu et al. 2013), and it was previously shown to successfully transform other duckweeds (Chhabra et al. 2011; Liu et al. 2019), including *S. polyrhiza* in transient transgene expression (Dombey et al. 2025).

Small callus fragments were inoculated with *Agrobacterium* carrying the *pZmUBQ::eGFP* T-DNA vector and co-cultured for two days before rinsing off the *Agrobacterium* (**Figure 4a, Method S1, S6**). Calli were transferred to CIM medium containing antibiotics, ticarcillin and cefotaxime, to select against residual *Agrobacterium* for recovery (**Figure 4b, Method S1, S6**). To confirm transformation and assess the impact of *Agrobacterium* on callus health, calli appearance and *GFP* expression were regularly monitored. Already two days post inoculation (dpi), bright GFP fluorescence spots were observed under UV light (**Figure 4b**). Calli grew for two weeks without displaying browning or any side effects from exposure to the bacteria, while GFP fluorescent sectors increased in number and size (**Figure 4b**).

**Figure 4:**
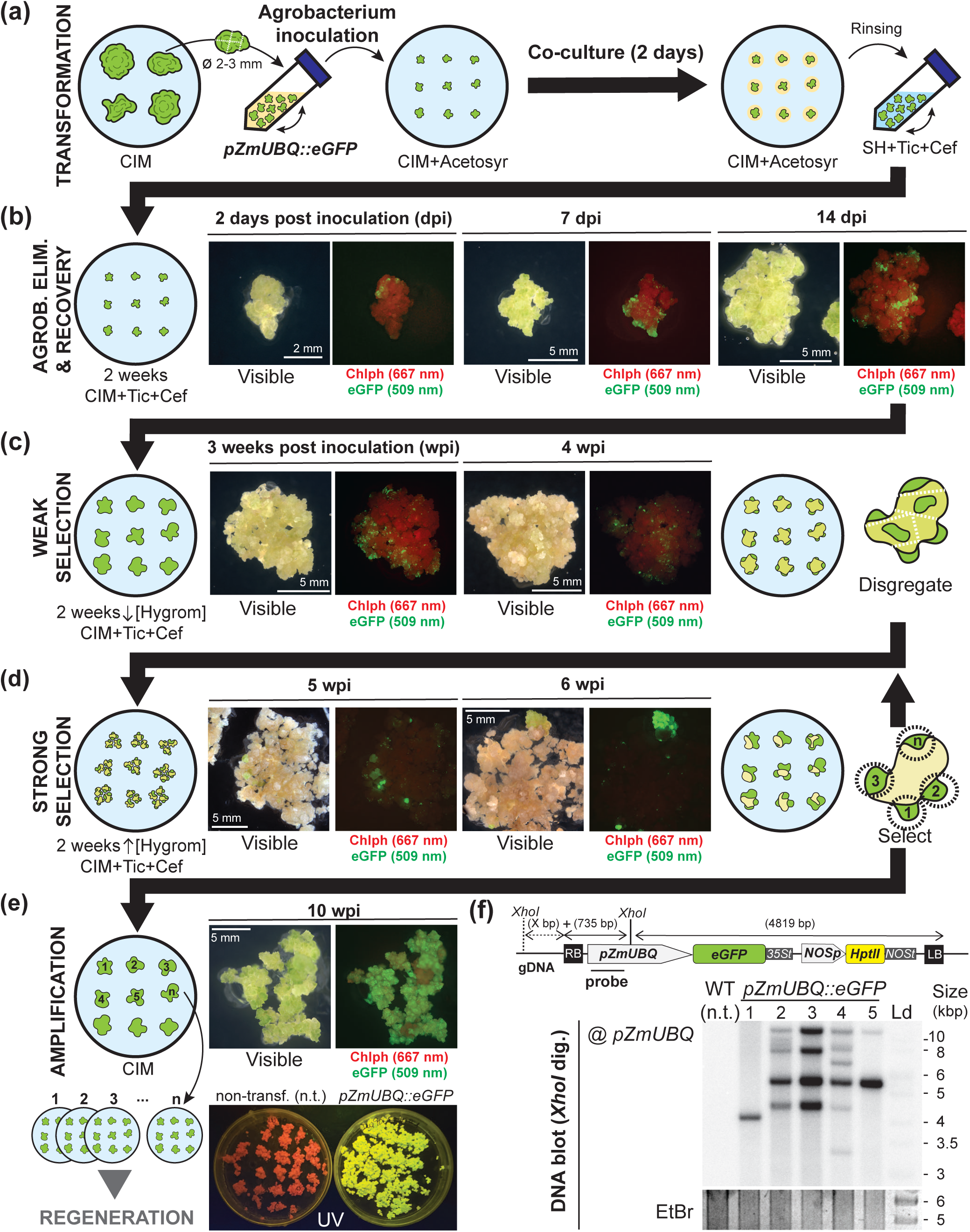
*Agrobacterium*-mediated stable transformation and selection of *S. polyrhiza SP162* calli. **(a)** Schematic representation of the calli transformation procedure. **(b)** Representative visible and fluorescent images of *SP162* calli in recovery 2, 7 and 14 days after inoculation (dpi) with *Agrobacterium* (EHA105) carrying a T-DNA vector for constitutive e*GFP* expression. Red: chlorophyll (Chlp) autofluorescence (excitation 655 nm, detection 667 nm), green: eGFP (excitation 488 nm, detection 509 nm). **(c,d)** Same as in (b) but during weak (5 mg/L hygromycin, c) and high (10 mg/L hygromycin, d) selection of transformed cells. **(e)** Schematics of isolation and amplification of hygromycin-resistant calli and visible and fluorescent images of selected transgenic callus and plates of non-transformed and transgenic calli. In all images: green: GFP fluorescence, red: chlorophyll autofluorescence. **(f)** Southern blot analysis of T-DNA integration events in five independent calli. T-DNA integration was assessed by digesting gDNA with *XhoI* and detected with a probe against the *pZmUBQ* as indicated in the T-DNA scheme. Ethidium bromide (EtBr) staining of the gel prior to transfer is shown as loading control. n.t.: non-transformed regenerated wild-type (WT) control; Ld: DNA size ladder.

Hygromycin selection was applied progressively to avoid browning of dying cells, which can have toxic effects on neighboring transformed cells (Sangwan et al. 1992; Zhang et al. 2020; Liu et al. 2024). Weak selection with a low hygromycin concentration (5 mg/L) resulted in chlorosis and loss of chlorophyll autofluorescence (**Figure 4c, Method S1, S6**). After two weeks, selection pressure was increased to 10 mg/L hygromycin (**Method S1, S6**), causing further chlorosis and browning, while hygromycin-resistant, GFP-positive sectors started to enlarge (**Figure 4d**). These healthy sectors were transferred to either CIM medium without antibiotics for amplification or kept on 10 mg/L hygromycin if resistant sectors were small, heterogenous, or difficult to isolate from non-resistant tissue (**Figure 4d,e, Method S6**). Individual hygromycin-resistant sectors were subcultured to obtain larger quantities of fast-growing and uniformly GFP-positive calli, candidates for successful integration of the transgenes (**Figure 4e**).

T-DNA integration into the genome was confirmed by Southern blot analysis of five calli cultures, revealing single to multiple T-DNA insertion events (**Figure 4f**). Thus, *Agrobacterium*-mediated transformation enables stable genetic transformation of *SP162* calli.

### Transformed *SP162* calli regenerate into stable transgenic lines

Since T-DNAs can become unstable during tissue culture and plant regeneration (Peerbolte et al. 1987; Phillips et al. 1994; Risseeuw et al. 1997), we investigated T-DNA stability in line #1 (**Figure 4f**) with a single insertion of the *pZmUBQ::eGFP* transgene. As non-transformed *SP162* calli, transgenic calli started to regenerate after four weeks in FRM, with the formation of darker frond-like structures with higher chlorophyll autofluorescence (**Figure 5a**). Transgenic frond regeneration also proceeded normally after acclimation (**Figure 5b**). Fully regenerated GFP-positive fronds started to appear within 7-8 weeks and were able to clonally propagate (**Figure 5c**). Ten regenerated lines were amplified and subjected to Southern blot analysis. All lines carried the single T-DNA insertion (**Figure 5d**) from the initial callus line. Therefore, at least for this single T-DNA insertion, the integrated DNA remained stable during tissue culture and frond regeneration.

**Figure 5:**
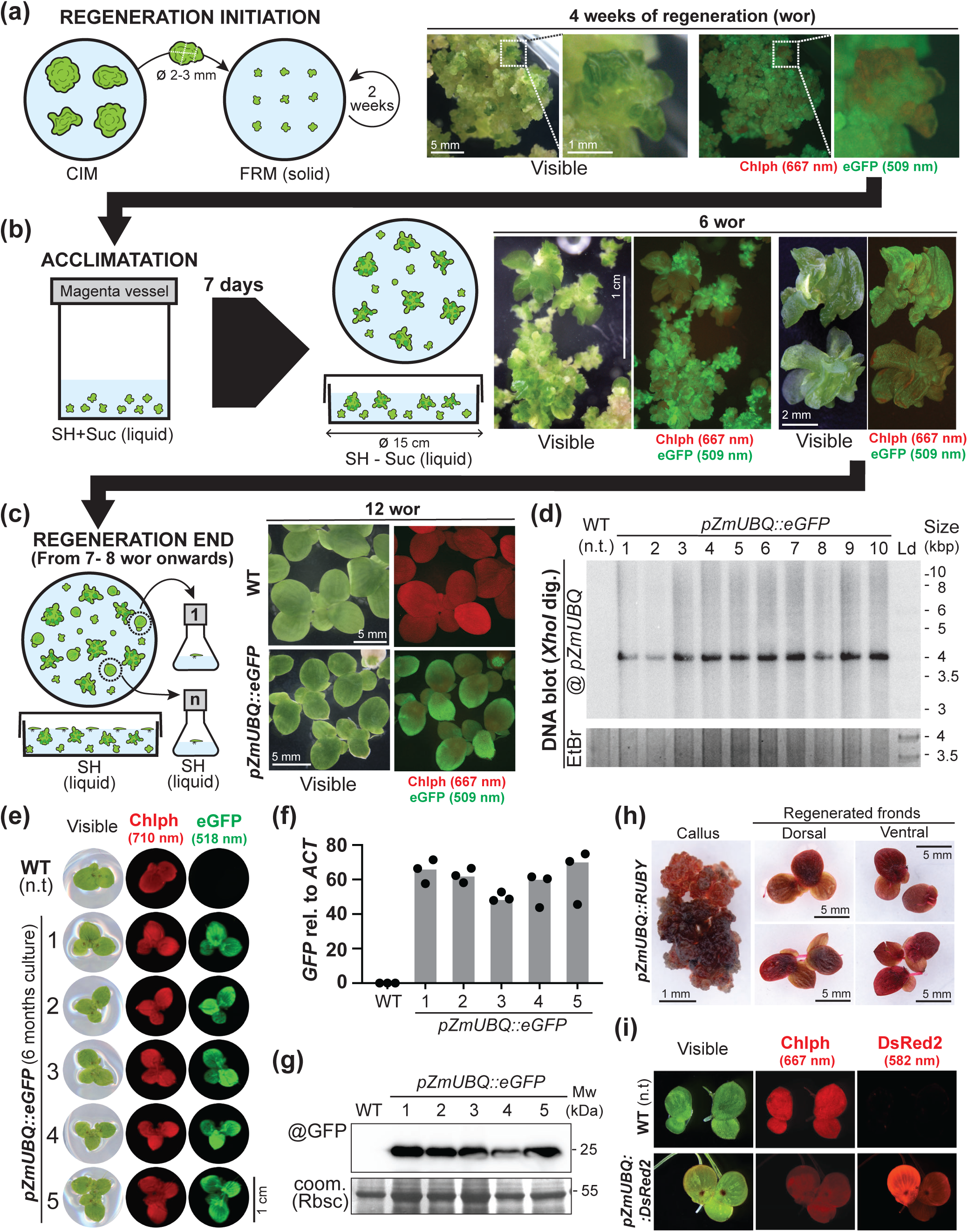
Frond regeneration and molecular characterization of *S. polyrhiza SP162* transgenic plants. **(a-c)** Schematics and representative visible and fluorescent images of *SP162* single T-DNA insertion transgenic calli expressing e*GFP* during the regeneration initiation (a), at the end of acclimation (b) or at the end of regeneration (c). In all images, red: chlorophyll (Chlp) autofluorescence (excitation 655 nm, detection 667 nm), green: eGFP (excitation 488 nm, detection 509 nm). **(d)** Southern blot analysis of T-DNA integration in 10 cultures of independently regenerated transgenic fronds. T-DNA integration was assessed as indicated in Figure 4f. Ethidium bromide (EtBr) staining of the gel prior to transfer is shown as loading control. n.t.: non-transformed regenerated wild-type (WT) control; Ld: DNA size ladder. **(e)** Images of individual fronds from one non-transformed and five transgenic regenerated lines scanned under white light or in a biomolecular scanner. Red: chlorophyll (Chlp) autofluorescence (excitation 658 nm, detection 710 nm), green: eGFP (excitation 488 nm, detection 518 nm). **(f)** qPCR analysis of e*GFP* expression relative to endogenous *ACTIN* mRNA levels in the same regenerated lines as in (e). Median values (bars) from 3 technical replicates (dots) are represented for each biological sample. **(g)** Protein blot analysis of GFP accumulation in the same lines as in (e,f). Rubisco (Rbcs) bands in Coomassie (coom.) staining of the blot after GFP immuno-detection is shown as loading control. Mw: Molecular weight. **(h)** Images of transgenic callus and regenerated fronds expressing the *RUBY* visual reporter. **(i)** Visible and fluorescent images of transgenic fronds expressing the *DsRed* fluorescent reporter. Chlp (chlorophyll): excitation 590 nm, detection 612 nm; DsRed2: excitation 561 nm, detection 582 nm.

Furthermore, after continuous culture for six months, GFP fluorescence was homogeneous across single-insertion regenerated lines (**Figure 5e**), with high levels of *GFP* mRNA expression and comparable levels of GFP protein accumulation across lines (**Figure 5f,g**). Thus, transgene expression also remained stable after prolonged periods of clonal propagation.

### *SP162* transformation can be independently reproduced with different reporter genes and selectable markers

To test the reproducibility of *S. polyrhiza SP162* transformation, the process was repeated with different reporter genes and in a different laboratory. First, we tested the betalain-based visual reporter *RUBY, pZmUBQ::RUBY* (He et al. 2020). Both, transgenic calli and regenerated fronds, displayed the bright red color characteristic of betalain accumulation (**Figure 5h**). Next, the transformation and regeneration procedures were independently reproduced in a second laboratory to obtain transgenic *S. polyrhiza* cultures transformed with *pZmUBQ::eGFP* (**Figure S5**), as well as other fluorescent reporters such as *DsRed2* (*pZmUBQ::DsRed2*) (**Figure 5i, S5b**), or using spectinomycin resistance as selectable marker (**Figure S5b, Method S6**). These results showcase the applicability of different reporter genes or selectable markers and further consolidate the reproducibility and practicality of the methodology used here to perform genetic transformation in the *S. polyrhiza SP162* strain.

### A near-chromosomal level SP162 genome assembly was generated using long-reads

We have previously sequenced the *SP162* genome using short-read sequencing and found a low intra-specific diversity in *S. polyrhiza* at the single nucleotide polymorphisms level (Wang et al. 2024). Nonetheless, to account for structural variations between different *S. polyrhiza* strains and provide a high-quality reference genome to guide genetic editing, we sequenced the genome of *SP162* on a flowcell of PromethION. The sequencing produced a total of 20.2 million reads, of which >99.7% passed base-calling filters. The dataset comprised 25.2 billion bases, with a read N50 length of ∼4.5 kb. The assembly results in a near-chromosome level 139.5 Mbp genome assembly (**Figure S7**). For consistency on gene IDs across *S. polyrhiza* strains, genome annotations were transferred from those of the *9509* reference genome (Hoang et al. 2018). To facilitate access to the *SP162* genome, we created a web-server to browse its genome (Robinson et al. 2011), retrieve gene sequences and find orthologous genes (Steinegger and Söding 2017) (https://agxu.uni-mainz.de/SP162/).

### CRISPR-Cas9 genome editing can be achieved in *SP162* to generate mutations

Lack of regular flowering and sexual reproduction in *S. polyrhiza* (Fourounjian et al. 2021) hinders classic mutagenesis methods that require selfing or backcrossing to fix desired mutations and clean the background (Koornneef 2002). However, CRISPR/Cas technology allows to precisely introduce mutations through targeted genome editing (Jung and Till 2021; Cardi et al. 2023). Hence, we tested if CRISPR/Cas9 was suitable to generate mutants in *S. polyrhiza SP162*.

The CRISPR/Cas9 genome editing requires to design single-guide RNAs (sgRNAs) that exactly match the DNA target site. Using the newly assembled SP162 genome, we then built a single-vector sgRNA multiplexing CRISPR/Cas9 system based on Liu’s laboratory tools (Ma et al. 2015; Ma and Liu 2016). We used the intron-containing and maize codon-optimized *Cas9* (*zCas9i*), shown to improve editing efficiencies in plants (Grützner et al. 2021; Stuttmann et al. 2021), under the *ZmUBQ* promoter (**Figure 6a**). To facilitate selection and monitor *zCas9i* expression, we transcriptionally fused *GFP* to *zCas9i* through the *porcine teschovirus-1 2A* ribosome skipping peptide (2A), allowing independent translation of Cas9 and GFP proteins from the same ORF (*pZmUBQ::zCas9i:P2A:eGFP*; **Figure 6a**). As 2A amino acids remain appended at the C-terminus of the upstream protein, serving as tag for Cas9 detection.

**Figure 6:**
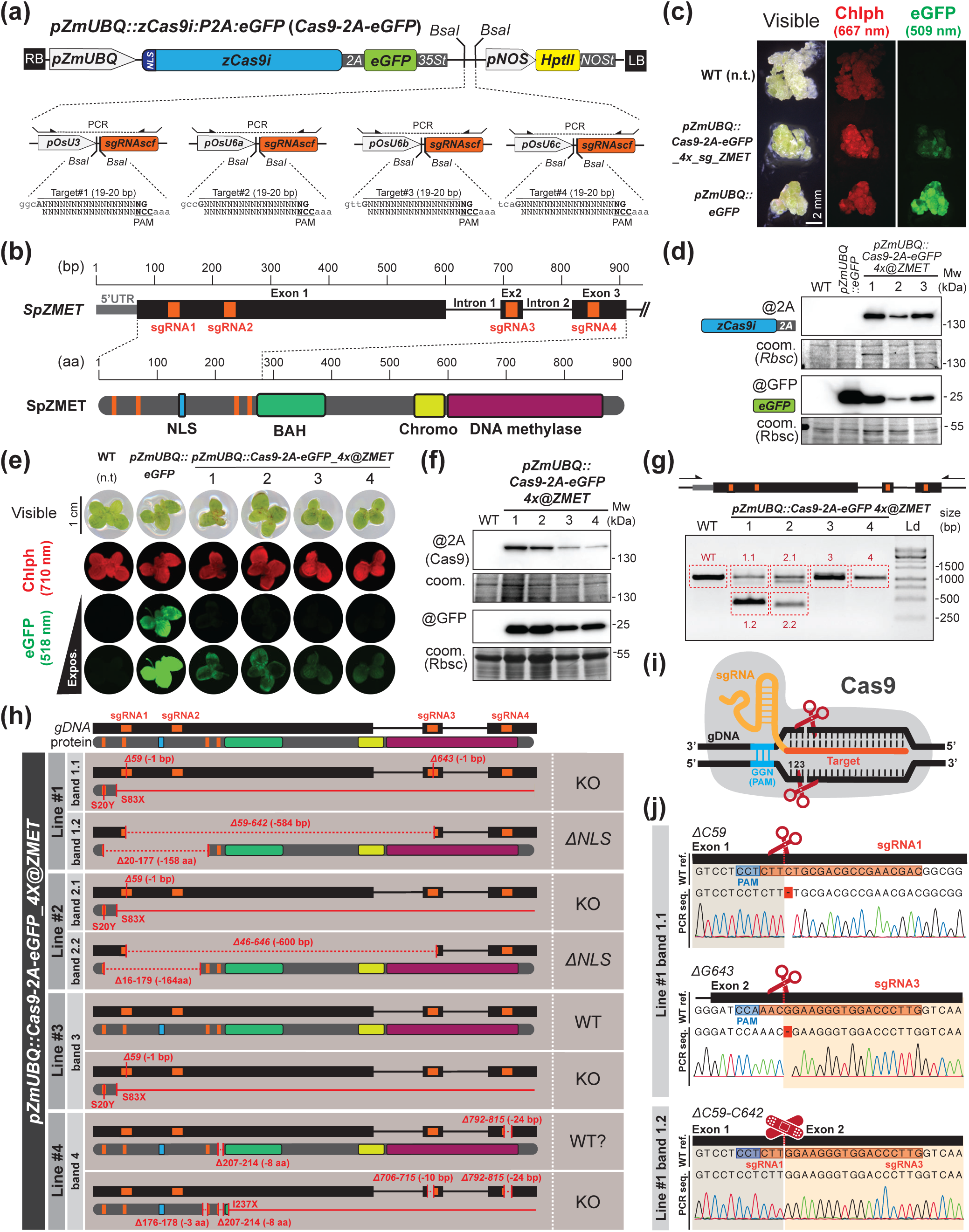
CRISPR/Cas9-mediated genome editing in *S. polyrhiza SP162*. **(a)** Vector for simultaneous expression of *zCas9i-2A-eGFP* and multiplexed sgRNAs. **(b)** Schematic representation of the *SpZMET* gDNA region targeted by sgRNAs (orange boxes), the SpZMET protein and its domains. NLS: nuclear localization signal; BAH: Bromo-adjacent homology domain; Chromo: Chromatin organization modifier domain; DNA methylase: C-5 cytosine-specific DNA methylase. **(c)** Visible and fluorescent images of WT and transgenic *SP162* calli transformed with the indicated plasmids. In UV images, red: chlorophyll (Chlp) autofluorescence (excitation 655 nm, detection 667 nm), green: eGFP (excitation 488 nm, detection 509 nm). **(d)** Protein blot analysis of zCas9i-2A and GFP accumulation in regenerated WT, e*GFP,* and three *zCas9i-2A-eGFP 4x@ZMET* calli. Rubisco (Rbcs) in Coomassie (coom.) staining of blots is shown as loading control. Mw: Molecular weight. **(e)** Images of individual fronds from one non-transformed (WT), one *eGFP* and four *zCas9i-2A-eGFP 4x@ZMET* transgenic regenerated lines scanned under white light or in a biomolecular scanner. Red: chlorophyll (Chlp) autofluorescence (excitation 658 nm, detection 710 nm), green: eGFP (excitation 488 nm, detection 518 nm). **(f)** Protein blot analysis of zCas9i-2A and GFP accumulation in non-transgenic and four *zCas9i-2A-eGFP 4x@ZMET* regenerated lines. Rubisco (Rbcs) in Coomassie (coom.) staining of blots is shown as loading control. **(g)** Agarose-gel electrophoresis of PCR amplification of *SpZMET* gDNA in non-transgenic and four *zCas9i-2A-eGFP 4x@ZMET* regenerated lines. Primers located outside of sgRNA targeted regions were used as indicated by arrows in the diagram above. Bands framed in red were isolated for subsequent cloning and Sanger sequencing analysis to investigate editing events. Ld: Ladder. **(h)** Graphical summary of the analysis of PCR products from (g) displayed in Figure S8. Resulting changes in DNA and predicted resulting protein sequences are shown. **(i)** Schematic representation of Cas9-mediated cleavage relative to the Protospacer Adjacent Motif (PAM) and target sgRNA sequence. **(j)** WT reference sequence, and Sanger sequencing chromatograms of examples of single nucleotide deletion and large nucleotide deletion between sgRNAs found in *SP162* regenerated line #1. PAM sequence is highlighted in blue and target sequence in orange.

As proof-of-concept, we targeted *ZMET* (SpZMET), a chromomethyltransferase required for DNA methylation at CHG sites (Bewick et al. 2017). Because *S. polyrhiza* has low CHG methylation (Michael et al. 2017; Dombey et al. 2025), we hypothesized that SpZMET loss-of-function would not impair callus survival or regeneration. To test small (1-100 bp) and larger (500-800 bp) deletions, four sgRNAs were designed in pairs flanking the SpZMET nuclear localization signal (NLS) (**Figure 6b**). The resulting plasmid (*pZmUBQ::zCas9i:P2A:eGFP_4x@ZMET*) was transformed into *SP162* calli as described above. Although GFP fluorescence was weaker in *Cas9* transgenic calli than in GFP-only lines (**Figure 6c**), western blotting confirmed Cas9-2A and GFP accumulation (**Figure 6d**). Cas9 expression did not impair frond regeneration and, albeit GFP fluorescence was low, Cas9-2A and GFP continued to be expressed in regenerated lines (**Figure 6e-f**).

Editing was first assessed in four regenerated lines by PCR amplification of the region targeted by the sgRNAs. Two of the lines produced additional shorter amplicons, compatible with large (∼600 bp) deletions (**Figure 6g**). Sequencing of PCR products confirmed that in three lines, all sequenced molecules carried mutations ranging from single nucleotide to larger (600 bp) deletions (**Figure 6h, S8**). Mutations were consistent with those expected from Cas9-mediated cleavage upstream of the third nucleotide after the PAM sequence (Jinek et al. 2012) (**Figure 6i**), resulting in single nucleotide deletions (SND) or elimination of sequences between two cleavages (**Figure 6h,j, S8**). Other known mutations induced by Cas9, such as single nucleotide insertions or larger deletions (2-20 nt), were also observed (**Figure 6h, S8**). Low frequency mutations (e.g. sgRNA4 at band 1.1, 1.2 and 3; **Figure S8**) likely resulted from constitutive Cas9 expression leading to chimeric individuals or cultures.

SND and small deletions generally resulted in out-of-frame mutations, generating putative knockout (ko.) alleles in all lines (**Figure 6h**). In-frame deletions between sgRNA1 and sgRNA3 eliminated much of the N-terminal region, including the NLS, in lines #1 and 2 (**Figure 6h,i**). Thus, our data demonstrate that CRISPR/Cas9 efficiently induced both frameshift and domain-specific deletions in *SP162*, so that the transformation protocol can be applied to create targeted gene editing in *S. polyrhiza*.

To ease the use and further development of gene editing and other CRISPR/Cas-based methods in *SP162*, we implemented a gRNA design function for diverse CRISPR applications (McKenna and Shendure 2018) in the above-mentioned web-server.

### Regenerated *SP162* lines display phenotypic variation

Despite the reproducibility and stability of transformation, we noticed that lines of regenerated plants looked and grew differently from the original *WT SP162* and between themselves. The most obvious observable difference was frond size, as regenerated lines were smaller than *WT SP162* (**Figure S9a**). While this difference could be as little as 0.5 mm in length, *pZmUBQ::RUBY* lines were up to 2 mm, about 30%, shorter than *WT* (**Figure S9b**).

In addition, cultures of regenerated plants initiated from similar frond numbers grew to occupy less surface and displayed increased senesce in the same amount of time compared to *WT* (**Figure S10a-c**). While reduced culture surfaces could be explained by smaller frond size, their growth rates were also lower (**Figure S10d**). Although growth patterns of all lines still fit an exponential curve (**Figure S10e,f**), doubling times were delayed by nearly one day for *GFP* lines (3.1-3.5 days) and two for *RUBY* lines (4.2-4.5 days), almost twice the doubling time of *WT* (2.4 days) (**Figure S10f**). Similar phenotypes were observed in the *Cas9-*expressing lines (**Figure S11**).

The reduced growth rates of *RUBY* lines could be consequence of metabolism rerouting and tyrosine depletion for betalain production (He et al. 2020), as high *RUBY* expression negatively affects growth and development also in other plants (Lee et al. 2023; Tse et al. 2024; Swinnen et al. 2025). Thus, phenotypic variation between *RUBY* lines (**Figure S9, S10**) could be due to different transgene expression levels. Likewise, there was phenotypic variation between the *Cas9* lines (**Figure S11**). It remains to be determined whether such variation is the result of the modified trait in other cases of mutants or transgene-expressing lines. However, similar variation was observed between *GFP* lines (**Figure S9, S10**) that all originate from the same single T-DNA insertion event (**Figure 4f, 5d**). Hence, the underlying cause of the phenotypic differences might be induced rather by tissue culture and/or regeneration.

### Phenotypic variation in regenerated lines is not associated with changes in ploidy

Plants obtained through tissue culture usually display phenotypic variation from parental lines due to somaclonal variation generated during callus induction, propagation and/or regeneration (Bairu et al. 2011). Although its causes can be of epigenetic or genetic origin (Evans and Sharp 1986; Neelakandan and Wang 2012; Wibowo et al. 2022), changes in ploidy often correlate with changes in organ and plant size (Kuppu et al. 2015; Corneillie et al. 2018; Šmarda et al. 2023; Dermail et al. 2024). This is of special relevance for genome editing as gene copy number increases through polyploidy impact editing efficiency (Sánchez-Gómez et al. 2023). Therefore, we analyzed DNA content and estimated genome size of several regenerated lines, including those expressing *GFP* and *Cas9*, through flow cytometry (**Figure S12a,b**). The estimated 1C-value was about 160 Mbp (**Figure S12c**), close to the 139.5 Mbp obtained by genome sequencing, and compatible with *SP162* being a diploid strain (Hoang et al. 2022). No significant variations in genome size were observed between the original *WT SP162* and regenerated lines (**Figure S12c**). Hence, although a more detailed analysis of their genomes and epigenomes will be required to investigate the underlying cause behind the observed phenotypic variation, changes in ploidy during *SP162* tissue culture can be excluded.

### *SP162* is also suitable for transient expression

In addition to stable transformation, *Agrobacterium*-mediated transient expression enables rapid transgene testing without the arduousness and time requirements of tissue culture. As the technique has been previously applied and achieved high levels of protein expression in other *S. polyrhiza* strains (Peterson et al. 2021; Dombey et al. 2025), we tested GFP (*pZmUBQ::eGFP*) transient expression in *SP162* using the *Agrobacterium* vacuum infiltration (agro-infiltration) method (Dombey et al. 2025) (**Method S7**).

GFP foci appeared three days post infiltration (dpi) and expanded for two weeks (**Figure 7a, S13**), a pattern independently reproduced in a second laboratory (**Figure S14**). However, after 21 days, infiltrated plants appeared unhealthy, showing chlorosis and growth reduction (**Figure 7b**). Such symptoms resembled those of *S. polyrhiza* infected with pathogenic bacteria (Baggs et al. 2022). Indeed, bacteria extracted from surface-sterilized fronds grew on selective media for *Agrobacterium* (rifampicin and streptomycin) and for plasmid-containing cells (rifampicin, streptomycin and spectinomycin) (**Figure 7c**). Thus, *Agrobacterium* likely persisted within tissues despite the continuous presence of cefotaxime and ticarcillin antibiotics (**Figure 7a, Method S7**), compromising long-term culture viability.

**Figure 7:**
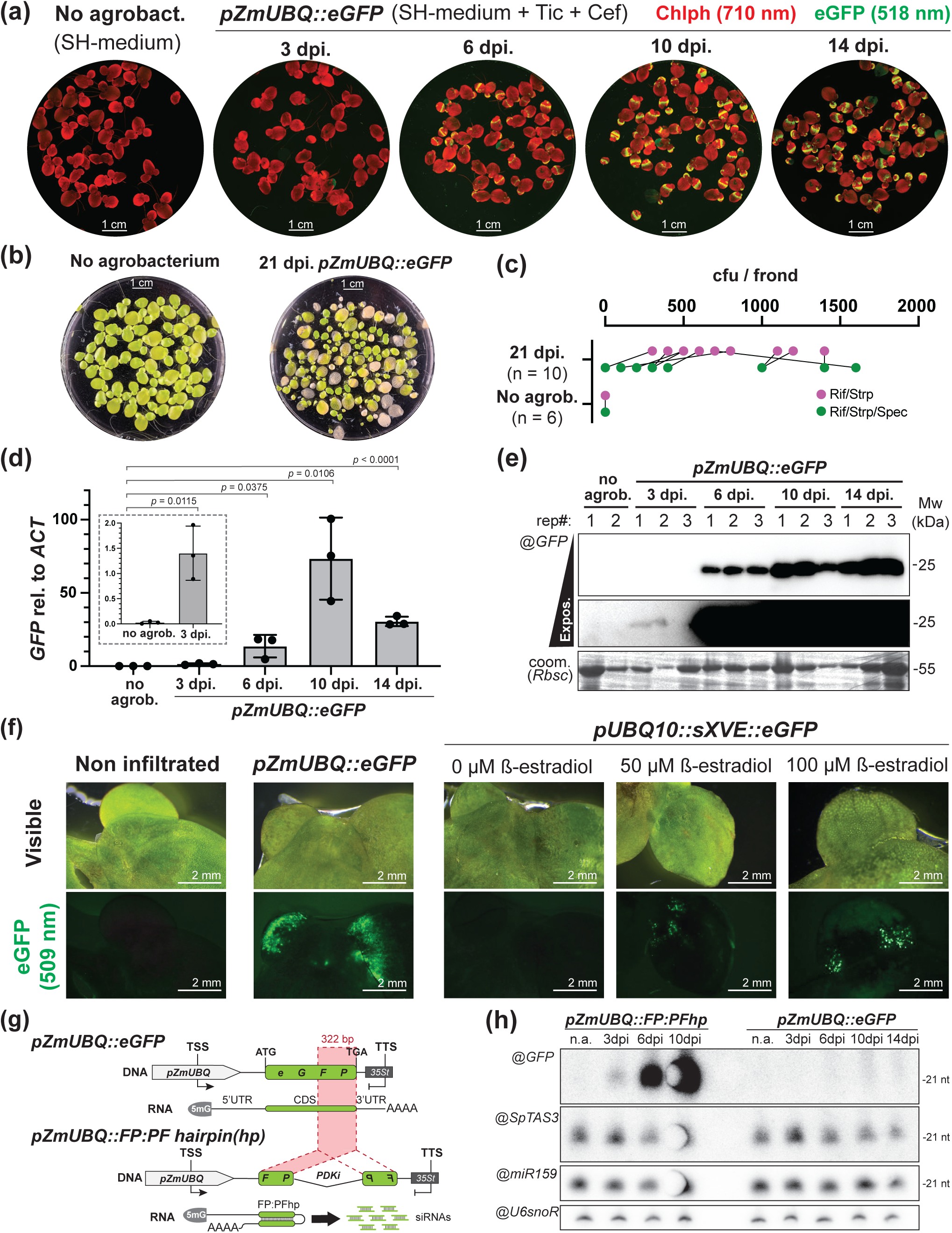
*Agrobacterium*-mediated transient expression in *S. polyrhiza SP162*. **(a)** Biomolecular scanning of eGFP fluorescence (green) and chlorophyll (Chlp) autofluorescence (red) of *SP162* cultures infiltrated with *pZmUBQ::eGFP* at different days post infiltration (dpi). Red: chlorophyll autofluorescence (excitation 658 nm, detection 710 nm), green: eGFP (excitation 488 nm, detection 518 nm). **(b)** Images of non-infiltrated and 21 dpi infiltrated cultures. **(c)** Colony forming units (cfu) in bacteria extracts from individual fronds from samples in (b) plated on *Agrobacterium* EHA105 selective media (magenta; rifampicin, streptomycin) or for bacteria carrying the *pZmUBQ::eGFP* plasmid (green; rifampicin, streptomycin, spectinomycin). Lines link extracts from single fronds plated in the two media **(d)** qPCR analysis of e*GFP* expression relative to endogenous *ACTIN* mRNA levels at the indicated samples and timepoints. Mean values (bars) from 3 biological replicates (dots) are represented for each sample. Error bars represent standard deviation of the mean. *p* indicates two-tailed P-value of unpaired t-test between the indicated groups. **(e)** Two exposure times of protein blot analysis of GFP accumulation during transient expression in three biological replicates (rep#) for each sample. Rubisco (Rbcs) in Coomassie (coom.) staining of blots is shown as loading control. Mw: Molecular weight. **(f)** Visible and fluorescent images of *SP162* fronds infiltrated with *eGFP* under the control of an estradiol-inducible promoter in presence or absence of estradiol. eGFP (excitation 488 nm, detection 509 nm) Non-infiltrated and constitutively driven *eGFP* infiltrated samples are shown as control. Darker zones in visible and chlorophyll auto-fluorescent images result from liquid infiltration in plant tissues during vacuum *Agrobacterium* infiltration **(g)** Schematic representation of *pZmUBQ::eGFP* and the *eGFP* derived hairpin (hp) *pZmUBQ::FP:PFhp* plasmids. **(h)** Small RNA blot analysis of samples used in (a-e) and infiltrated with *pZmUBQ::FP:PFhp*. *eGFP*-derived siRNAs were probed against full-length *eGFP*. Detection of *SpTAS3*- tasiRNAs served as control for siRNA biogenesis; *miRNA159* and *U6* snoRNA detection served as size and loading control, respectively.

To determine transgene expression dynamics, GFP RNA and protein levels were monitored. *GFP* mRNA was already detected at 3 dpi and peaked at 10 dpi and, albeit lower, remained high at 14 dpi (**Figure 7d**). GFP protein levels mirrored this trend, low at 3 dpi (only detectable in one of the biological replicates), rising at 6 dpi and persisting to 14 dpi (**Figure 7e**). We also tested the applicability of the methodology to more complex expression systems with an estradiol-inducible GFP construct. As expected, fluorescence only occurred after estradiol application to the media (**Figure 7f**). Similar induction was obtained in stable transgenic calli and regenerated fronds (**Figure S15**). Thus, *SP162* supports robust transient expression as an experimental system and is suitable for testing constructs before investing into the time and labor for stable transformation.

### Transient transgene expression does not trigger S-PTGS in *S. polyrhiza*

The transient expression dynamics seen here resembled previously reported *GFP* expression driven from the viral promoter *p35S* in *S. polyrhiza,* lasting up to 30 dpi (Peterson et al. 2021). Such a long period of transient transgene activity is unusual. Generally, in most tested experimental model plants like *Nicotiana benthamiana*, expression decays within days after infiltration due to the onset of silencing. Sense post-transcriptional gene silencing (S-PTGS) upon overexpression of transgenes impedes continuous high mRNA levels during transient assays through the production of 21-22-nt-long short interfering (si)RNAs (**Figure S16a**) (Vaucheret et al. 1997; Voinnet et al. 2000; Hamilton et al. 2002; Kościańska et al. 2005; Luo and Chen 2007; Paoli et al. 2009; Parent et al. 2015). To counteract this, transgenes of interest are usually co-infiltrated with viral suppressors of RNA silencing (VSRs), like P19, to reduce S-PTGS (Voinnet et al. 2000; Hamilton et al. 2002; Qu et al. 2003; Mérai et al. 2005; Chen et al. 2013; Jay et al. 2023).However, P19 does not enhance GFP expression in *S. polyrhiza* (Peterson et al. 2021). Furthermore, while endogenous siRNAs (tasiRNAs) whose biogenesis depends on S-PTGS factors (Peragine et al. 2004; Yoshikawa et al. 2005; Gasciolli et al. 2005) are present (**Figure S16b**), no siRNA after *RUBY* agro-infiltration could be detected (Dombey et al. 2025), supporting no or only weak transgene-triggered S-PTGS.

Previous studies lacked either simultaneous siRNA or transgene mRNA measurements, especially relevant for *RUBY* as betalain pigmentation can remain long after transgene expression has faded (Tabara et al. 2024), leaving uncertainty about any correlation between mRNA levels and siRNAs. Therefore, we investigated the presence of *GFP*-derived siRNA in the same samples used to quantify *GFP* transient expression. As control, we infiltrated a *GFP-*derived hairpin construct (*pZmUBQ::FP:PFhp*) to trigger siRNAs independently of S-PTGS (**Figure 7g**, **S16c**) and monitored *SpTAS3-*tasiRNAs (**Figure S16d-g**). *GFP* siRNAs in cultures infiltrated with *FP:PFhp* were already detected at 3 dpi and accumulated strongly at 6 and 10 dpi but were undetectable in plants expressing *GFP*, even at 10 dpi with maximum *GFP* mRNA levels (**Figure 7h**). However, *SpTAS3*-tasiRNAs were present in all samples (**Figure 7h**), further supporting the hypothesis that the transgene-induced branch of S-PTGS might be reduced or compromised in *S. polyrhiza*, enabling prolonged transgene expression.

## Discussion

Despite recent advances (Yamamoto et al. 2001; Cantó-Pastor et al. 2015; Yang et al. 2018; Liu et al. 2019), the deployment of duckweeds as model organisms has been hindered by the lack of publicly available strains accessible for genetic modification and suitable protocols. We identified *S. polyrhiza SP162* as a likely diploid, competent strain for stable transformation and genome editing and provide detailed, versatile, and reproducible protocols, including also transient expression. In addition, and to facilitate the adoption of *S. polyrhiza* in research laboratories, we have also made *SP162* available through the EuropeanƒNorth American and Asian duckweed germplasm collections under the ID#: *5676*, along with its genome and other aspects of *SP162* culture and a web-based genome browser allowing to retrieve gene sequences, find orthologs, and design CRISPR gRNAs. Hence, we hope that *SP162* serves as a common platform to stimulate duckweed research and the approach, tools, and procedures described here will help screen and identify other suitable strains for transformation.

The variability in callus induction and response to growth regulators during regeneration observed by us and others (Wang 2016; Huang et al. 2018; Yang et al. 2018), suggests indeed natural diversity and potential to identify additional strains transformable with appropriate media and phytohormone compositions by systematically testing different conditions. Furthermore, novel technologies such as expression of *GRF-GIF* chimeric morphogenic regulators, shown to significantly boost transformation, editing, and regeneration efficiencies in plants (Debernardi et al. 2020; Maher et al. 2020; Vandeputte et al. 2024; Swinnen et al. 2025) could further boost these processes in *SP162* and other genotypes.

Future research should address some of the current limitations of the system. Mutations introduced via genome editing might impair regeneration, limiting the recovery of mutant lines in *S. polyrhiza* without the chance to maintain certain alleles in the heterozygous configuration and generate homozygous mutants only upon sexual reproduction. Our successful use of inducible promoters to drive Cas9 expression after regeneration should circumvent this hurdle and reduce the continuous appearance of mutations due to constitutive expression.

Additionally, the origin of somaclonal variation warrants investigation to understand and minimize its confounding effects during phenotypic analysis of transgenic or mutant lines. While genetic causes cannot be excluded, the observed phenotypes - reduced size, slower growth, and early senescence - are characteristic of *Arabidopsis* regenerated from root- but not from leaf-derived callus due to the inheritance of root-specific epigenetic signatures (Wibowo et al. 2022). Therefore, as most of the calli in our study were derived from roots, it remains possible that the phenotypic variation is of epigenetic origin. Hence, not only the epigenetic profiles of regenerated plants, but also the difference between those regenerated from shoot- (budding pocket) or root-derived calli, deserve future investigation.

Future research should also address some of the current limitations of the system. Mutations introduced via genome editing might impair regeneration, limiting the recovery of mutant lines. In addition, limited sexual reproduction in *S. polyrhiza* impedes means to segregate away the Cas9 transgene, which under continuous expression might lead to the recurrent appearance of off-target site mutations. The use of inducible promoters, shown here to work in *SP162*, to drive Cas9 expression during controlled periods of time after regeneration should help circumvent these hurdles.

Finally, while stable transgenesis and genome editing are powerful tools, transient expression is a versatile technique with multiple applications such as molecular farming or as an experimental system to quickly assess protein localization, function, or interactions (Chen et al. 2013; Krenek et al. 2015). The absence of S-PTGS during transient expression in *S. polyrhiza*, allowing high transgene expression over prolonged periods of time, offers advantages over more traditional systems like *Nicotiana benthamiana*, particularly as a monocot transient expression platform. Given the inability of eudicots to properly splice complex introns, monocot genes cannot be directly expressed from genomic DNA and often require cDNA cloning to function in eudicots (Keith and Chua 1986; Goodall and Filipowicz 1991; Sinibaldi and Mettler 1992; Nitovska et al. 2018; Dombey et al. 2025), which is not always feasible. Moreover, despite recent advances (Li et al. 2025), widespread recalcitrance to *Agrobacterium* in monocots limits its application for transient expression and generally requires the use of viruses or direct DNA delivery to plant cells by complex methods, such as microinjection, biolistics, or electroporation and transfection of protoplasts (Lee et al. 2015; Ren et al. 2021), entailing specialized equipment and skills. Therefore, *S. polyrhiza* also offers a robust, fast, low-cost and technically simple monocot transient expression system.

## Materials and Methods

### Plant material and growth conditions

*Spirodela polyrhiza* (L.) Schleid accessions, described in (Wang et al. 2024), were obtained from the Institute of Organismic and Molecular Evolution, Johannes Gutenberg University (Mainz, Germany). *SP162* was deposited at the Landolt-IBBA duckweed collection (Institute of Agriculture Biology and Biotechnology, CNR, Milano, Italy; https://biomemory.cnr.it/collections/CNR-IBBA-MIDW) (Morello et al. 2024), and the Rutgers Duckweed Stock Cooperative (RDSC; Rutgers University, New Jersey, USA; http://www.ruduckweed.org) under ID#: *5676*, and Prof. Huan Weijuan’s lab in Shenzhen University of Advanced Technology, China. *Lemna minor 8623* was obtained from the Landolt-IBBA collection.

Plants were grown under sterile conditions on liquid N-medium (Appenroth 2002), SH-medium (*SP162*) [3.2 g/L Schenk and Hildebrandt salts, Duchefa #S0225, pH 5.8] or 0,5X SH-medium (*9509*) (full compositions in **Method S1**). Growth containers included MagentaTM GA-7 boxes (Sigma Aldrich, #V8505), 6-well plates (Greiner Bio-one, #657185), 94 or 145 mm diameter petri dishes (Greiner Bio-one, #633181, #639161), glass beakers covered with a sterile plastic lid or Erlenmeyer flasks, sealed with Parafilm or Leucopore, inside growth chambers under long-day (16-h light / 8-h dark) at 21°C or 26°C with a light intensity of 70-125 µmol/m^2^/s. Plants were sterilized in 1% NaClO (**Method S2**). Turions were induced in N-medium at 21°C, stored at 4°C, and germinated at 21/26°C in SH-medium (**Method S3, S8**).

### Callus induction screening and regeneration test

Axenic cultures were grown in liquid N-medium with 10 g/L sucrose at 26°C under long-day (70 µmol/m^2^/s) for seven days before callus induction. Explants and CIM media were prepared as in (Wang 2016; Huang et al. 2018; Yang et al. 2018; Liu et al. 2019). In most experiments, callus induction rates (number of explants initiating callus formation) were recorded in triplicates. Calli growth rates were expressed as percentage of fresh-weight (FW) biomass gain of 20-50 calli at 7-10-day interval. Frond regeneration was first tested according to (Huang et al. 2018) in the *SP162* and *9509* accessions and later as in (Wang 2016; Yang et al. 2018) by adding 1 mg/L TDZ or 2.2 mg/L zeatin to the original Frond Regeneration Media by Huang and colleagues and later modified for *SP162* (see below).

### *SP162* callus induction and maintenance

Sterile fronds were transferred dorsoventrally to solid CIM medium [4.4 g/L MS-salts (Duchefa #M0245), 0.5 g/L MES-salts (Duchefa #M1503), 4 g/L Gelrite (Duchefa #G1101), 10 g/L sucrose, 0.1 g/L PVP-40 (Sigma-Aldrich #PVP40-100), 50 mg/L citric acid (Merck #27109-100G-R), 0,221 mg/L 2,4-D (Duchefa #D0911), 0.4 mg/L TDZ (Sigma-Aldrich #P6186-25MG), pH 5.8]. Plates were kept at 21°C under low light (20 µmol/m^2^/s) in long-day conditions (see above) and refreshed every two weeks until ∼2-3 mm calli were formed at the root tip. Calli were transferred to CIM plates at 26°C under the same light conditions. Appearing pale-green calli were transferred to separate plates under the same conditions. Sensitivity to antibiotics was performed in 5 to 6 replicates, each composed of a small round plate (60 x 15 mm) containing 3 callus fragments (∼3 mm). Sensitivity was tracked by weekly imaging and comparing the developed phenotypes (growth arrest, whitening, and/or browning and death) to calli growing on CIM without antibiotics. Media and step-by-step procedure can be found in **Method S1, S4**.

### *SP162* frond regeneration

Calli fragments (2-3 mm) were placed on FRM [4.4 g/L MS-salts, 0.5 g/L MES-salts, 4 g/L Gelrite, 10 g/L sucrose, 1 mg/L TDZ or 2.2 mg/L zeatin (Sigma-Aldrich #Z0164), pH 5.8] at 26°C under low light (20 µmol/m^2^/s) and long-day conditions and refreshed biweekly. Calli with signs of frond regeneration were transferred to liquid SH-medium with 1% w/v sucrose and cultured in Magenta boxes for 7-10 days at 26°C under high light (125 µM/m^2^/s) in long-day conditions. Regenerating calli were transferred to 145 mm diameter petri dishes with liquid SH-medium under the same conditions. Emerging fronds were transferred to individual flasks containing SH-media and maintained at 21 or 26°C. Media and step-by-step procedures are in **Method S1, S5**.

### Callus transformation and selection

*Agrobacterium* (*EHA105*) overnight culture was pelleted (3000 *g*, 20 min), resuspended to OD_600_ 0.7 in 30 ml agroinfection medium [10 mM MgCl_2_, 50 g/L sucrose, 200 µM acetosyringone (Sigma-Aldrich #D134406)] and incubated 1h in darkness. Fifty 2-3 mm calli were incubated 10 min in the solution under gentle agitation. After 10 min, calli were transferred to solid co-culture medium (solid CIM with 100 µM acetosyringone) and incubated 2 days at 26°C under low light (20 µmol/m^2^/s). Excess *Agrobacterium* was rinsed off from calli in liquid SH-medium with 0.3 mg/L cefotaxime (Duchefa #G0124.0025) and 0.25 mg/L ticarcillin (Duchefa #T0190.0010) for 1 min and transferred to agroelimination medium (solid CIM with 0.25 mg/L ticarcillin and 0.3 mg/L cefotaxime). After 2 weeks, calli were transferred to low selection media, solid agroelimination-CIM containing low-dose antibiotic (e.g., 5 mg/L hygromycin, Sigma-Aldrich #400051-1MU). Two weeks later, resistant-looking calli were transferred to media with increased selection pressure. Media were refreshed every two weeks until resistant calli started to proliferate and could be individually transferred to CIM for amplification. Detailed media and step-by-step procedures are in **Method S1, S6**.

### Transient expression in SP162

*Agrobacterium*-mediated transient expression was performed as in (Dombey et al. 2025). 70-80% confluent axenic *SP162* cultures were pretreated in ice-cold 20 mg\L L-glutamine (PanReac Applichem #A1420) for 20 min before being infiltrated twice at -70 kPa twice for 10 min with an induced OD_600_ 0.6 *Agrobacterium* solution in modified agroinfection medium [10 mM MgCl_2_, 50 g/L sucrose, 150 µM acetosyringone, 0.02% (v/v) Silwet L-77 (Kurt-Obermeier #7060-10)]. Co-cultured was carried in 94 mm diameter petri dishes with SH-medium containing 100 µM acetosyringone for three days in standard growth conditions. After, plants were transferred to SH-medium containing 0.25 mg/L ticarcillin and 0.3 mg/L cefotaxime refreshed weekly. Detailed media and step-by-step procedures are in **Method S1, S7**. Extraction and quantification of bacteria from infiltrated fronds are described in **Method S8**.

### Genome sequencing and annotation

DNA sequencing was performed in a PromethION flow cell (v10.4.1 FLO-PRO114M), assembled into 30 linear contigs using PECAT v0.0.3 (Nie et al. 2024) , and scaffolded into 20 pseudochromosomes via synteny with the *SP9509* reference genome using Chromosemble in Satsuma_v2 (Grabherr et al. 2010; Ernst et al. 2025). Liftoff v1.6.3 (Shumate and Salzberg 2021), was used to transfer annotations from the *SP9509* genome (Ernst et al. 2025). Further details in **Method S8**.

### Plant imaging and image-based phenotyping

Images were acquired with a Canon EOS M50 MarkII camera, Andonstar AD409 digital microscope, or Epson V600 scanner. Fluorescent images were taken using a fluorescence stereo zoom microscope Zeiss Axio Zoom V16 (chlorophyll autofluorescence: excitation 655 nm, detection 667 nm; eGFP: excitation 488 nm, detection 509 nm; dsRed2: excitation 561 nm, detection 582 nm) or a Sapphire FL Biomolecular scanner (Azure biosystems) (chlorophyll autofluorescence: excitation 658 nm, detection 710 nm; eGFP: excitation 488 nm, detection 518 nm). Images were analyzed using Ilastik (v1.4.1; https://www.ilastik.org), a supervised machine-learning software (Sommer et al. 2011), suitable for duckweed image analysis (Romano et al. 2022), or Fiji (V2.16.0; https://imagej.net/software/fiji/)(Schindelin et al. 2012). More details about image analysis in **Methods S8**.

### Plasmids generated in this study

Details on molecular cloning procedures, including sgRNA design, can be found in **Method S8**. All pre-existing used plasmids are publicly available from Addgene (https://www.addgene.org) and listed in **Method S8**. Plasmids newly generated in this study have been deposited in the European Plasmid Repository (https://www.plasmids.eu): *pZmUBQ::2x_BsaI::35St_HygR* (<u>#877</u>); *pZmUBQ::2x_BsaI::35St_KanR* (<u>#878</u>); *pZmUBQ::eGFP* (<u>#869</u>); *pZmUBQ::zCas9i:P2A:GFP* (<u>#870</u>); *pOsU3m::sgRNAscf* (<u>#872</u>); *pOsU6a::sgRNAscf* (<u>#873</u>); *pOsU6aLacZ::sgRNAscf* (<u>#874</u>); *pOsU6b::sgRNAscf* (<u>#875</u>); *pOsU6c::sgRNAscf* (<u>#876</u>); *pZmUBQ::zCas9i:P2A:GFP_4x_sg_ZMET* (<u>#871</u>); *pZmUBQ::FPPFhp* (<u>#879</u>). Plasmids were introduced into *Agrobacterium* through the freeze-thaw method (Weigel and Glazebrook 2006).

### DNA, sRNA, and protein blotting and detection

DNA, RNA, and protein blotting and detection followed standard procedures described in **Method S8**. Sequences of oligonucleotides used as probes or as PCR primers for Klenow-based DNA probes are listed in **Method S9**. For protein detection, the following antibodies were used. <u>Primary antibodies</u>: monoclonal (rat) anti-GFP [3H9] (Chromotek, #3H9-100), dilution 1:5000; monoclonal (mouse) anti-2A peptide antibody, clone 3H4 (Sigma Aldrich, #MABS2005-25UG), dilution 1:1000. <u>Secondary antibodies</u>: polyclonal (goat) anti-mouse HRP-conjugated (ThermoFisher, #62-6520), dilution 1:10000; polyclonal (goat) anti-rat HRP-conjugated (cell signalling, #7077S), dilution 1:10000.

### qPCR analysis of gene expression

RNA was extracted from 100 mg frozen ground tissue using TRIzol (#93289, Sigma). One µg of RNA treated with DNase I (Thermo Scientific, #EN0521) was reverse-transcribed with Maxima First-Strand cDNA Synthesis Kit (Thermo Scientific, #K1641). qPCR was performed on a QuantStudio5 (Applied Biosystems) system using KAPA SYBR FAST qPCR Kit (Roche, SFLRAB). Ct values were determined by the 2nd derivative max method of three technical replicates for each biological replicate. Relative expression values were calculated as the ratio of target Ct values relative to reference mRNA (*SpACT*) Ct. Oligonucleotides are listed in **Method S9**.

### Genome size estimation by flow cytometry

DNA content was estimated by flow cytometry following established methodology for plants (Doležel et al. 2007) and further detailed in **Method S8**. *L. minor 8623* was used as standard with a known DNA content of 1 C = 0,4182 pg (Hoang et al. 2019).

### Statistical analysis

Statistical tests were performed using GraphPad Prism (V10.2.3; https://www.graphpad.com) or R-dependent ggpubr (v0.6.0) (https://rpkgs.datanovia.com/ggpubr/). Statistical tests performed are indicated accordingly on each figure legend.

## Acknowledgements

We thank present and past members of the Marí-Ordóñez, Hubert and Xu groups for helpful discussions, in particular Daniel Buendía for advice regarding flow cytometry measurements of DNA content and Rodolphe Dombey for help installing and running sgRNA design tools. We are also grateful to the IMP Molecular Biology Service for providing reagents and Sanger sequencing. We thank Dr. Siân Stockton and Dr. Christian Siadjeu for help in isolating HMW DNA of *SP162*. We would also like to thank Andreas Wachter for providing plasmids and sharing the S1 facilities, the nucleic acid core facility from the faculty of biology at JGU Mainz for supporting the genome sequencing. The authors gratefully acknowledge the computing time granted on the supercomputer MOGON 2 at Johannes Gutenberg University Mainz (hpc.uni-mainz.de).

## Author contributions

A.M-O., S.X., and M.Hubert conceived and designed the study. M.Höfer performed callus induction screens. M.Höfer, R.E., and V.B-B tested regeneration in *9509* and *SP162*. V.B-B and R.E. stablished *SP162* culture conditions. E.D.R.B., A.S.L., and V.B-B tested antibiotic sensitivity. V.B-B established *SP162* transformation and regeneration after transgenesis and A.S.L. and E.R.D.B. reproduced it. C.D. designed sgRNAs and transformed CRISPR/Cas9 vectors. V.B-B., A.M-O., and R.E. characterized regenerated, transgenic, and genome edited lines. A.C. and S.X. sequenced and assembled the *SP162* genome. A.P-M. performed transient expression experiments. E.D.R.B. and A.S.L. validated transient expression and tested inducible promoters. S.X. and A.C. implemented the web-server. L.D-N. and L.P. performed turion experiments. A.M-O., S.X., M.Hubert, V.B-B., A.S.L., E.D.R.B., and M.Höfer analyzed the data. A.M-O assembled the figures and wrote the manuscript together with A.S.L., E.D.R.B., V.B-B., S.X., and M.Hubert.

## Data availability

All raw source data used in this study can be found in the Zenodo open repository under the following doi: 10.5281/zenodo.16970721. The sequencing data generated have been submitted to NCBI under bio-project number PRJNA1308930. The genome assembly and annotation have been deposited on Figshare: https://figshare.com/articles/dataset/Genome_assembly_and_annotation_of_i_Spirodela_polyrhiza_i_strain_SP162_ID_5676_/29942036. The web-server for *SP162* genome and gRNA design tool can be found here: https://agxu.uni-mainz.de/SP162/.

