## Supplemental Figures for "Strain, procedures, and tools for reproducible genetic transformation and genome editing of the emerging plant model *Spirodela polyrhiza* (L.) Schleid"

#### Table of contents:

| Page | |
|---|---|
| 2 | **Supplemental Figure S1:** *SP162* turion induction and germination. |
| 3 | **Supplemental Figure S2:** *SP162* callus induction under low light conditions. |
| 4 | **Supplemental Figure S3:** Initiation of *SP162* frond regeneration from pale calli. |
| 5 | **Supplemental Figure S4:** Termination of *SP162* frond regeneration after acclimatation in SH-liquid medium. |
| 6 | **Supplemental Figure S5:** Sensitivity of *SP162* callus to four antibiotics. |
| 7 | **Supplemental Figure S6:** Reproducibility of *SP162* transformation with different reporters and selectable markers. |
| 8 | **Supplemental Figure S7:** *SP162* genome assembly. |
| 9 | **Supplemental Figure S8:** Analysis of CRISPR/Cas9 induced mutations in *SP162*. |
| 11 | **Supplemental Figure S9:** Frond size variation among regenerated *SP162* lines. |
| 12 | **Supplemental Figure S10:** Growth analysis of regenerated *SP162* lines. |
| 13 | **Supplemental Figure S11:** Phenotypic analysis of regenerated *SP162* lines expressing *zCas9i*. |
| 14 | **Supplemental Figure S12:** Flow cytometry measurement of DNA content in regenerated *SP162* lines. |
| 15 | **Supplemental Figure S13:** GFP fluorescent development during transient expression. |
| 16 | **Supplemental Figure S14:** GFP fluorescent development during transient expression in the Hubert's lab. |
| 17 | **Supplemental Figure S15:** Stable estradiol-inducible GFP expression in *SP162* calli and regenerated fronds. |
| 18 | **Supplemental Figure S16:** siRNAs biogenesis in plants and tasiRNAs identification in *S. polyrhiza*. |

### SUPPLEMENTAL FIGURE S1

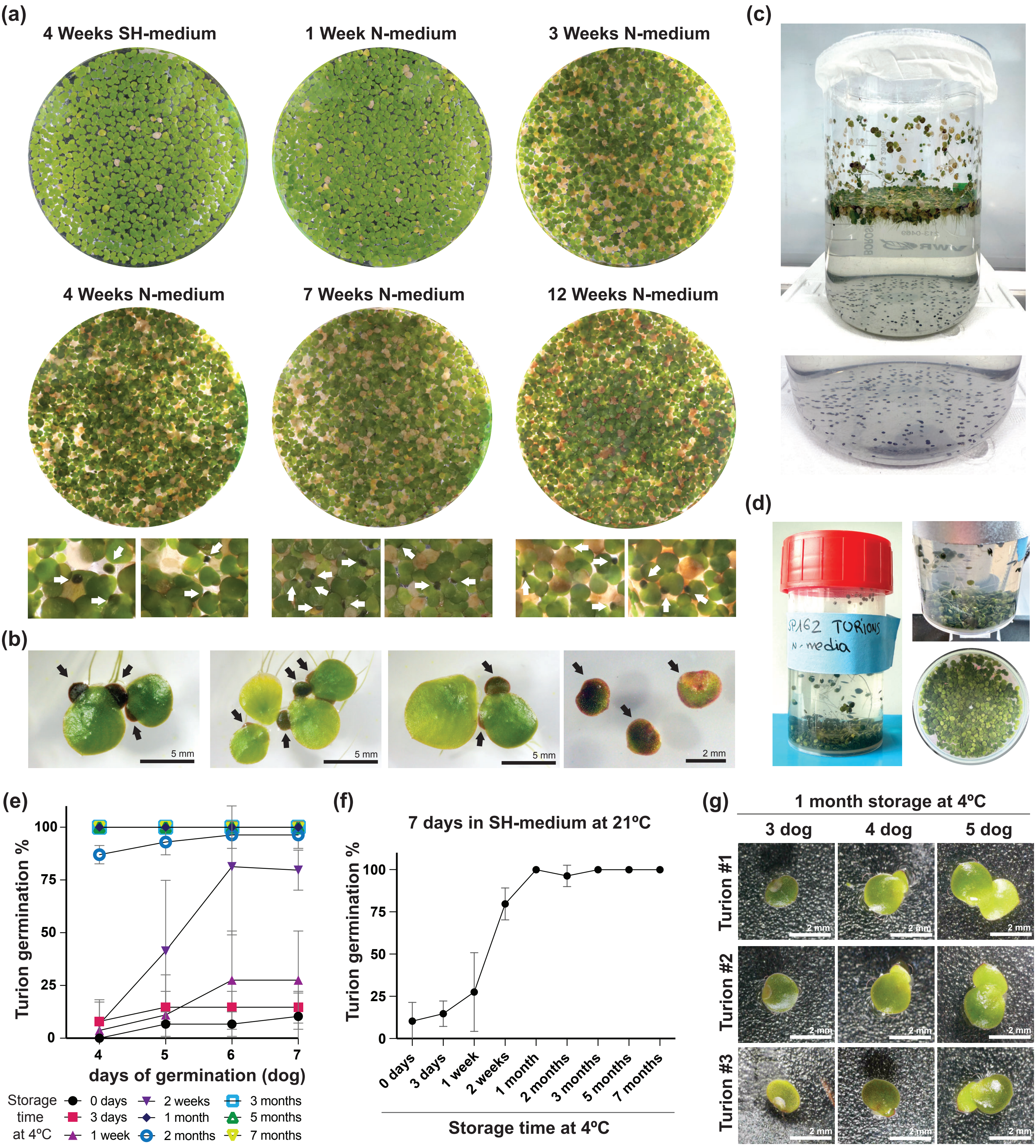


**Figure S1:** ***SP162*** **turion induction and germination.**
**(a)** Images of *SP162* cultures after 4 weeks in SH-medium and during turion induction in N-medium. **(b)** Representative images of SP162 turion development and released turions. **(c)** Image of *SP162* turion induction culture and sunk turions. **(d)** Pictures of turions during storage at 4ºC in Gossling red-capped containers. **(e)** Mean turion germination rates curves from 4 to 7 days of germination (dog) in SH-medium at 21ºC after different stratification (storage at 4ºC) times. Error bars represent standard deviation of the mean from 2-3 biological replicates. **(f)** Mean turion germination rates after 7 days in SH-medium at 21ºC from turions stratified (storage at 4ºC) for different periods of time. Error bars represent standard deviation of the mean from 2-3 biological replicates. **(g)** Representative images of early turion germination from three turions at 3-to-5 dog.

#### SUPPLEMENTAL FIGURE S2

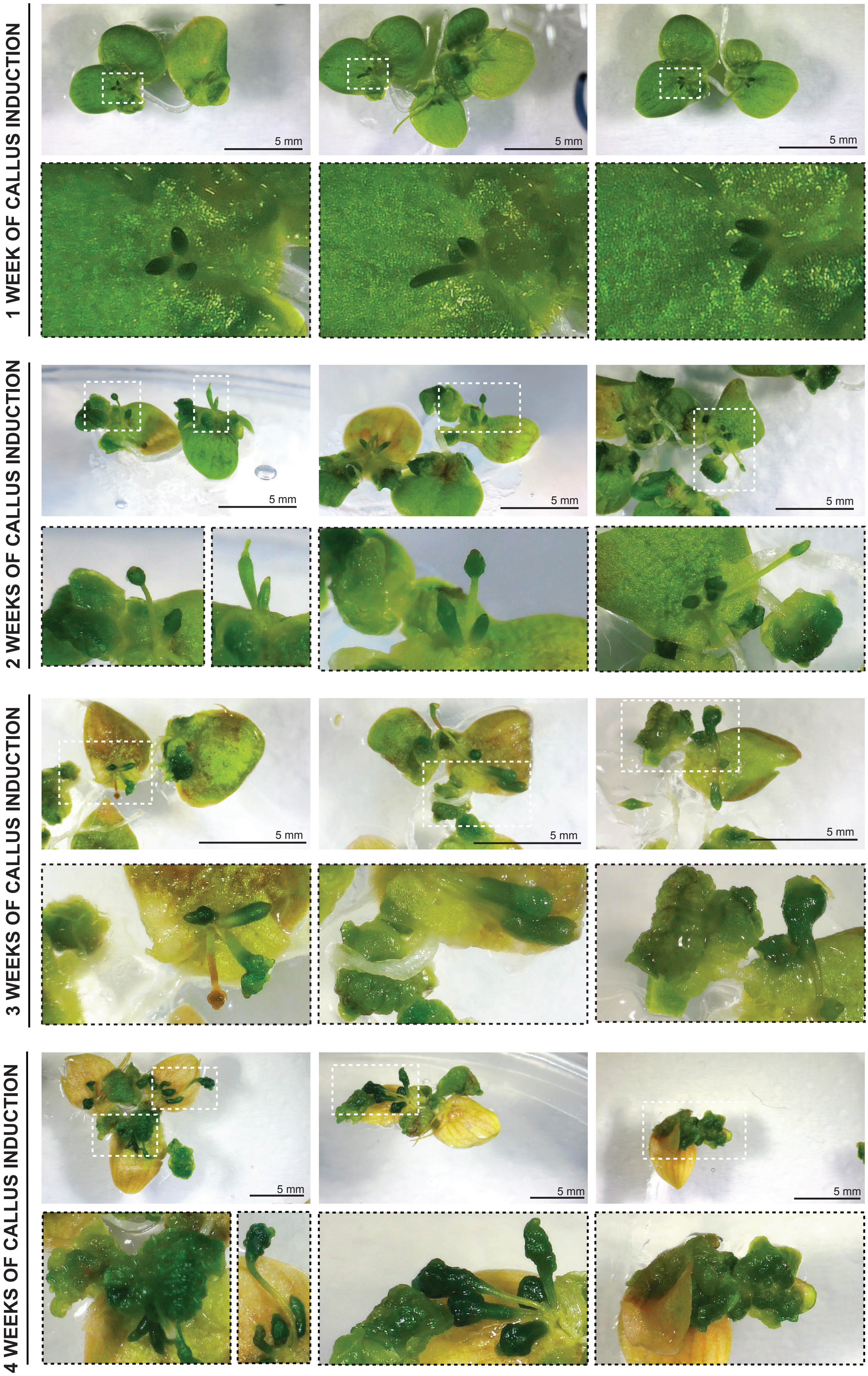


**Figure S2: *SP162* callus induction under low light conditions.**
Representative images of developing calli from roots and budding pockets after 1, 2, 3 and 4 weeks in callus induction medium in low light conditions under shade.

#### SUPPLEMENTAL FIGURE S3

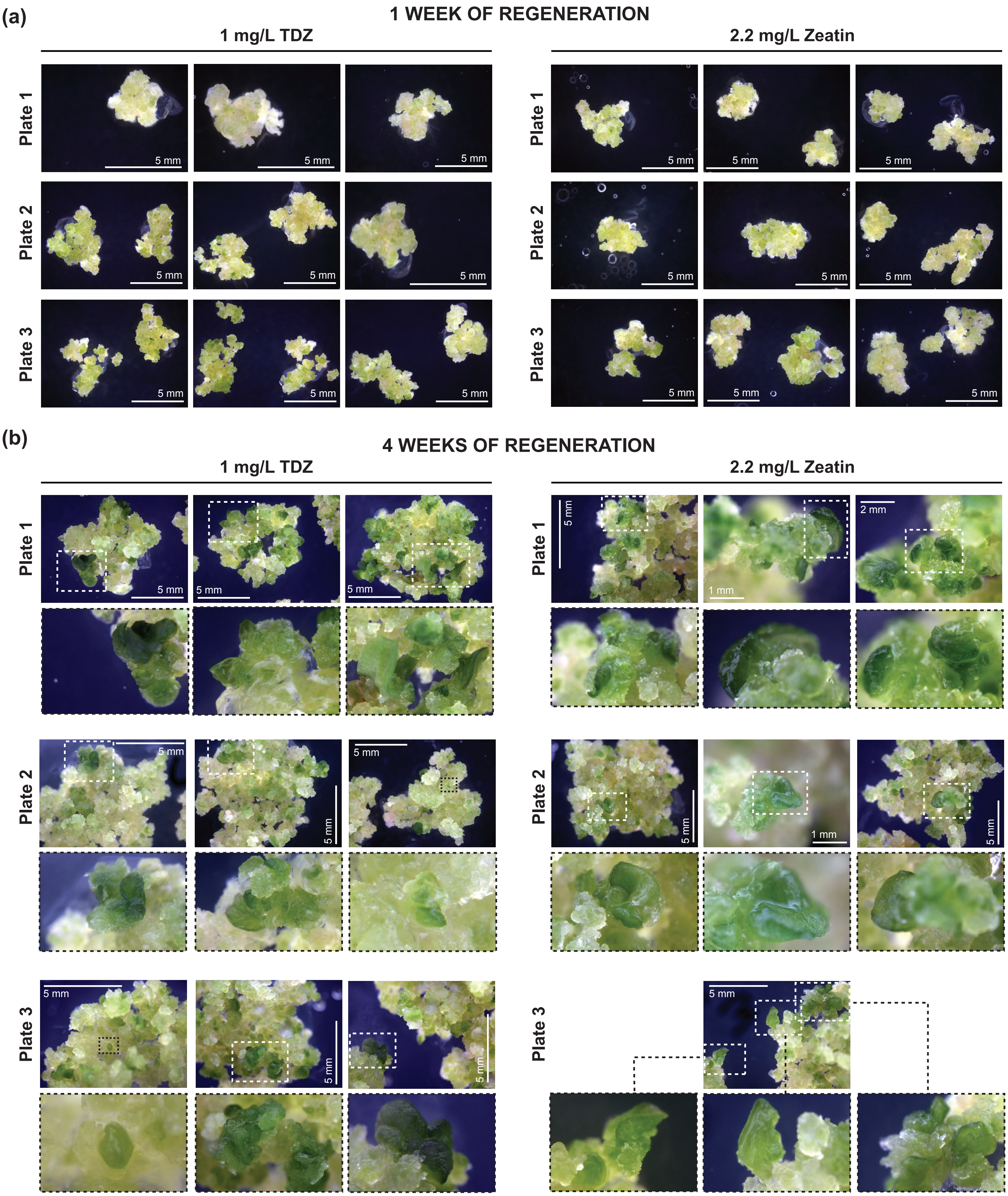


**Figure S3: Initiation of *SP162* frond regeneration from pale calli.**
Representative images of *SP162* calli in Frond Regeneration Medium (FRM) containing either 1mg/L TDZ or 2.2 mg/L Zeatin after 1 week (a) or 4 weeks (b) in FRM.

### SUPPLEMENTAL FIGURE S4

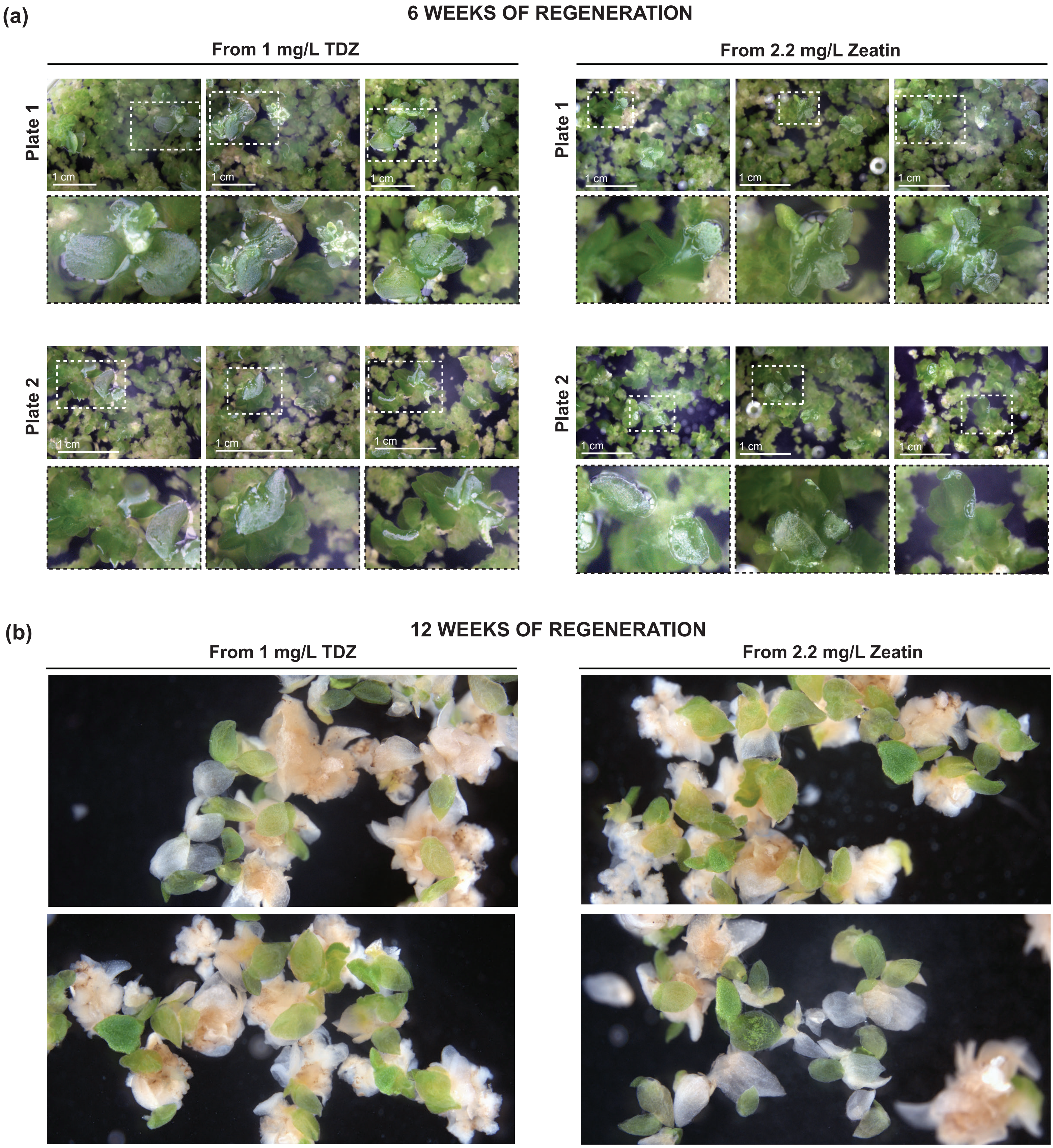


**Figure S4: Termination of *SP162* frond regeneration after acclimatation in SH-liquid medium.**
Representative images of *SP162* regenerating fronds in SH-medium 6 weeks **(a)** and 12 weeks **(b)** after initiation of regeneration in FRM with either TDZ or Zeatin.

### SUPPLEMENTAL FIGURE S5

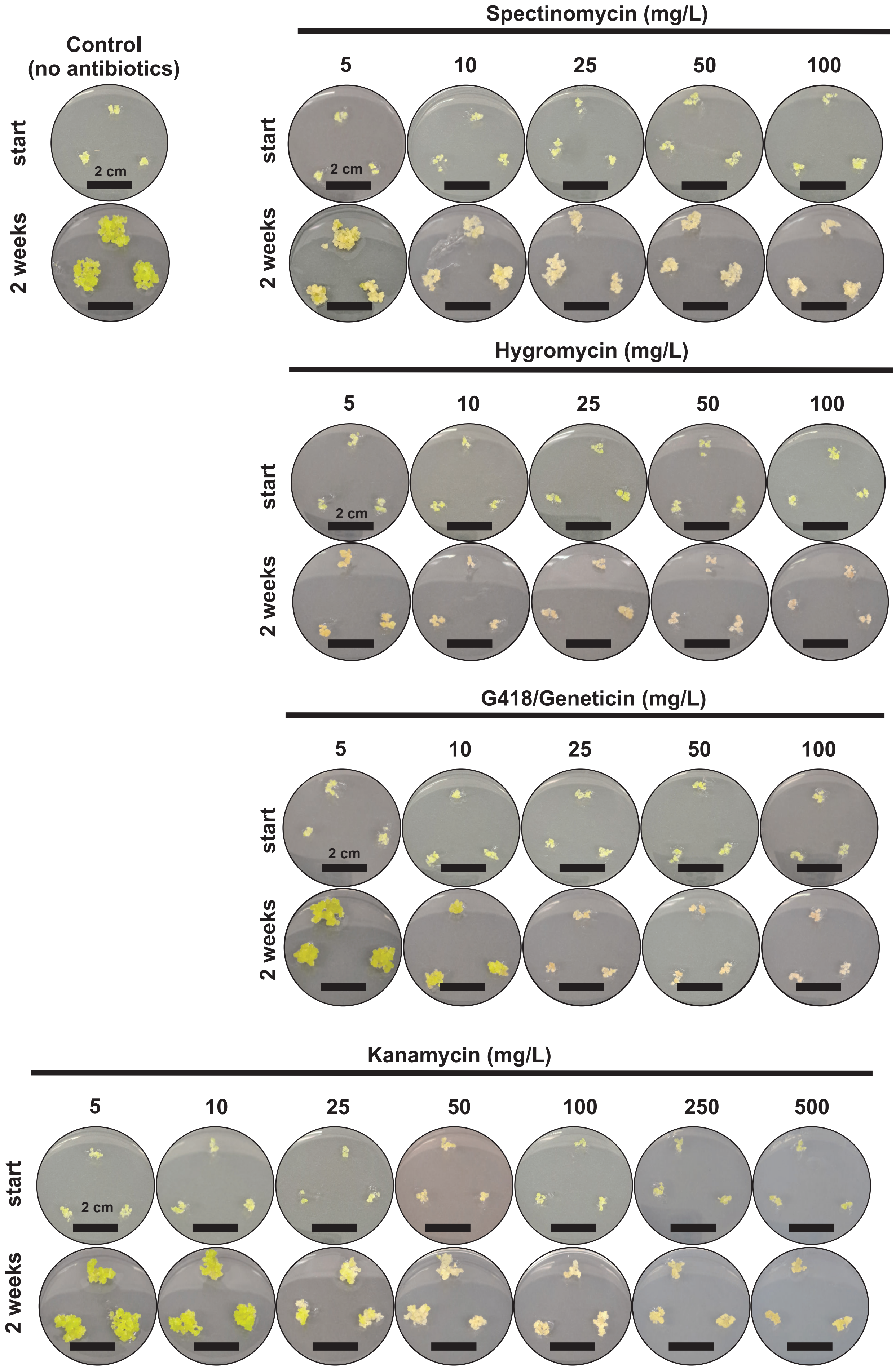


**Figure S5: Sensitivity of *SP162* callus to four antibiotics.**
*SP162* callus fragments (3-5 mm ø) cultured on CIM supplemented with inicated antibiotics or without any antibiotic (control). Each replicate consisted of 3 callus fragments grown onto 12 mL of CIM in small round culture plates (15 x 60 mm) under stadard callus induction and propagation conditions for *SP162.* In all cases n=6, all replicates are shown in the the Zenodo dataset associated to this manuscript (see Data avaliability).

#### SUPPLEMENTAL FIGURE S6

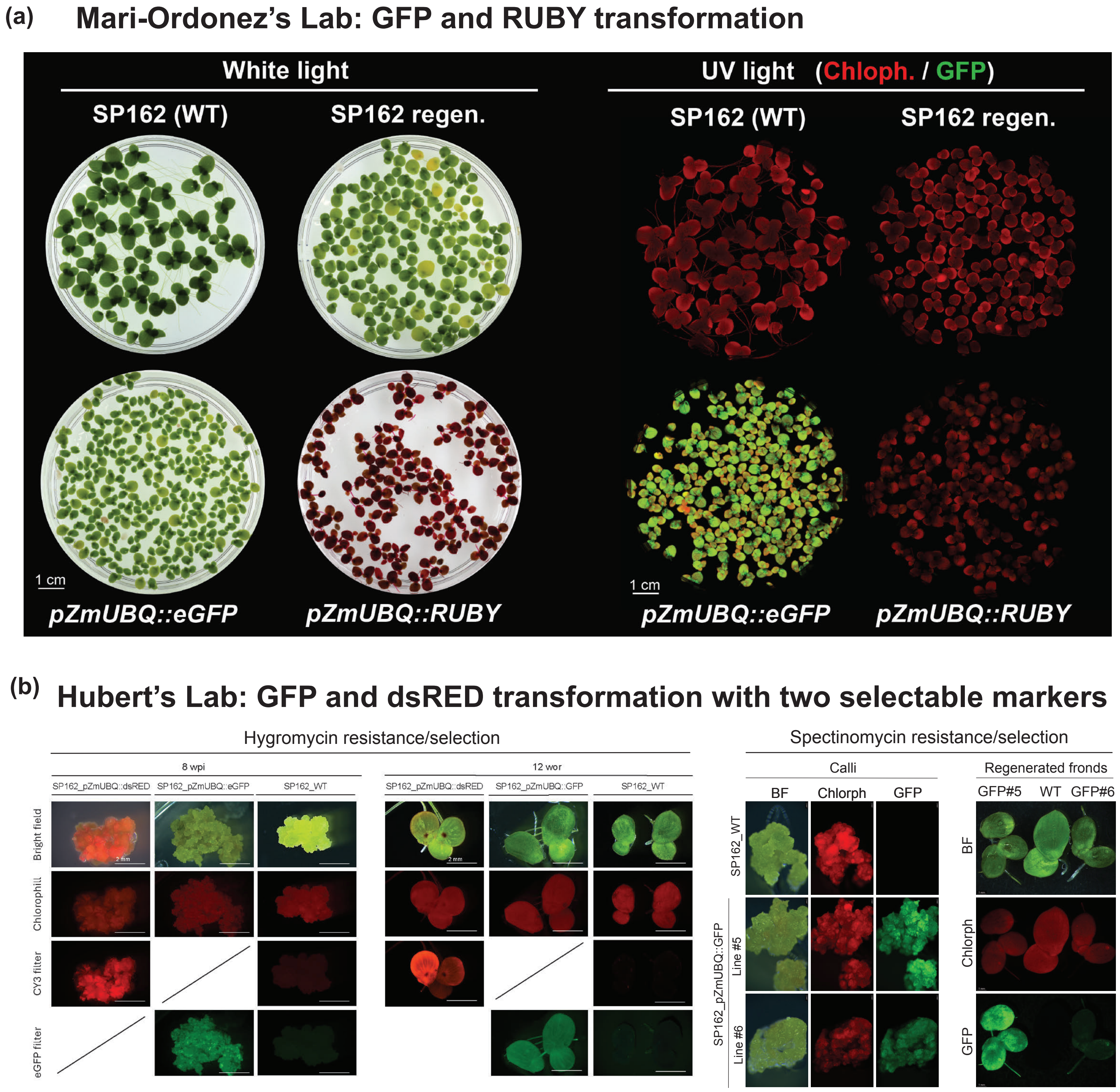


**Figure S6: Reproducibility of *SP162* transformation with different reporters and selectable markers.**
**(a)** Images under white or UV light of WT, regenerated non-transformed, and *eGFP* and *RUBY* transgenic *SP162* cultures obtained in the Marí-Ordóñez lab. **(b)** Reproduction of *SP162* transformation using *eGFP* and *DsRed* with both hygromycin or spectinomycin selectable markers in the Hubert lab.

### SUPPLEMENTAL FIGURE S7

(a)

| Genome assembly statistics | |
|---|---|
| **Assembly size** | 139.5 Mb |
| **Number of pseudochromosomes** | 20 |
| **N50** | 8.04 Mb |
| **BUSCO completeness** | 98.6% (viridiplantae_odb10)<br>96.9% (embryophyta_odb10) |
| **Number of coding genes** | 20,705 |

(b)

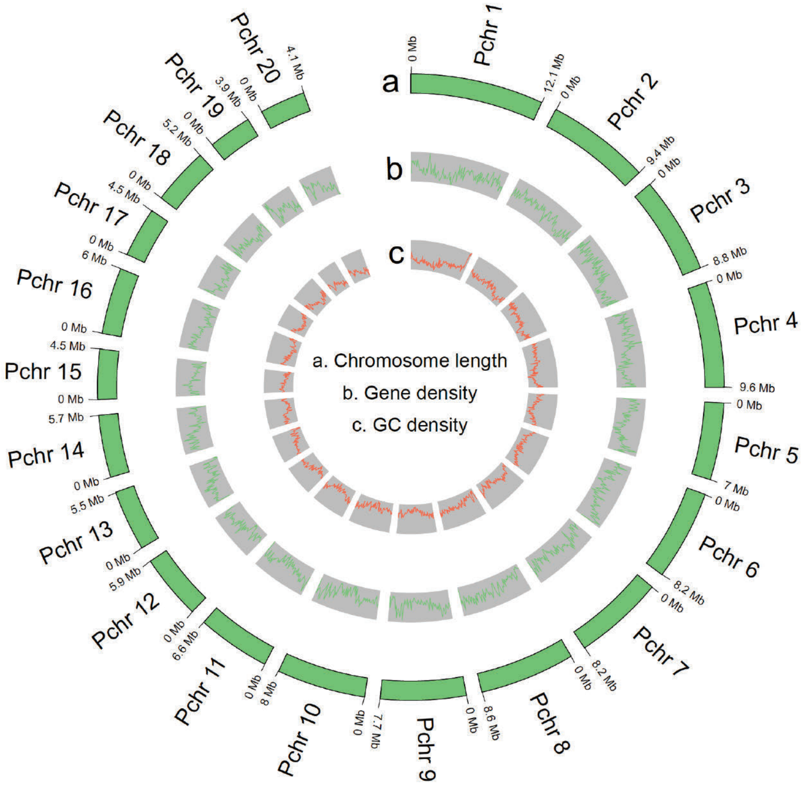


(c)

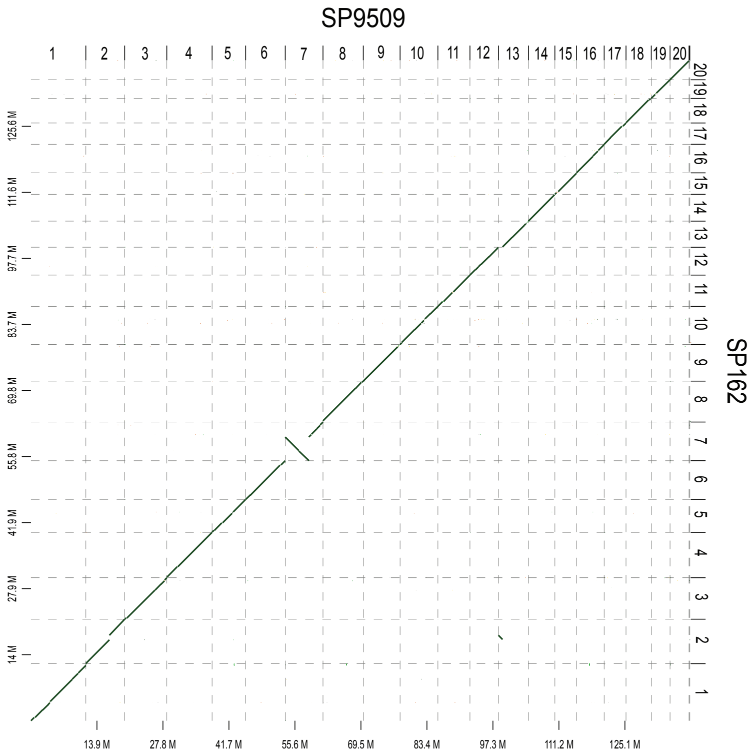


**Figure S7:** ***SP162*** **genome assembly.**
**(a)** *SP162* genome assembly and annotation sumary statistics. **(b)** Circos plot of *S. polyrhiza* strain *SP162* pseudochromosome-level genome assembly. **(c)** Dot-plot comparison of the Asian *SP162* pseudochromosomes (this study) and the European *9509* chromosomes. An inversion was founnd on chromosome 7.

### SUPPLEMENTAL FIGURE S8

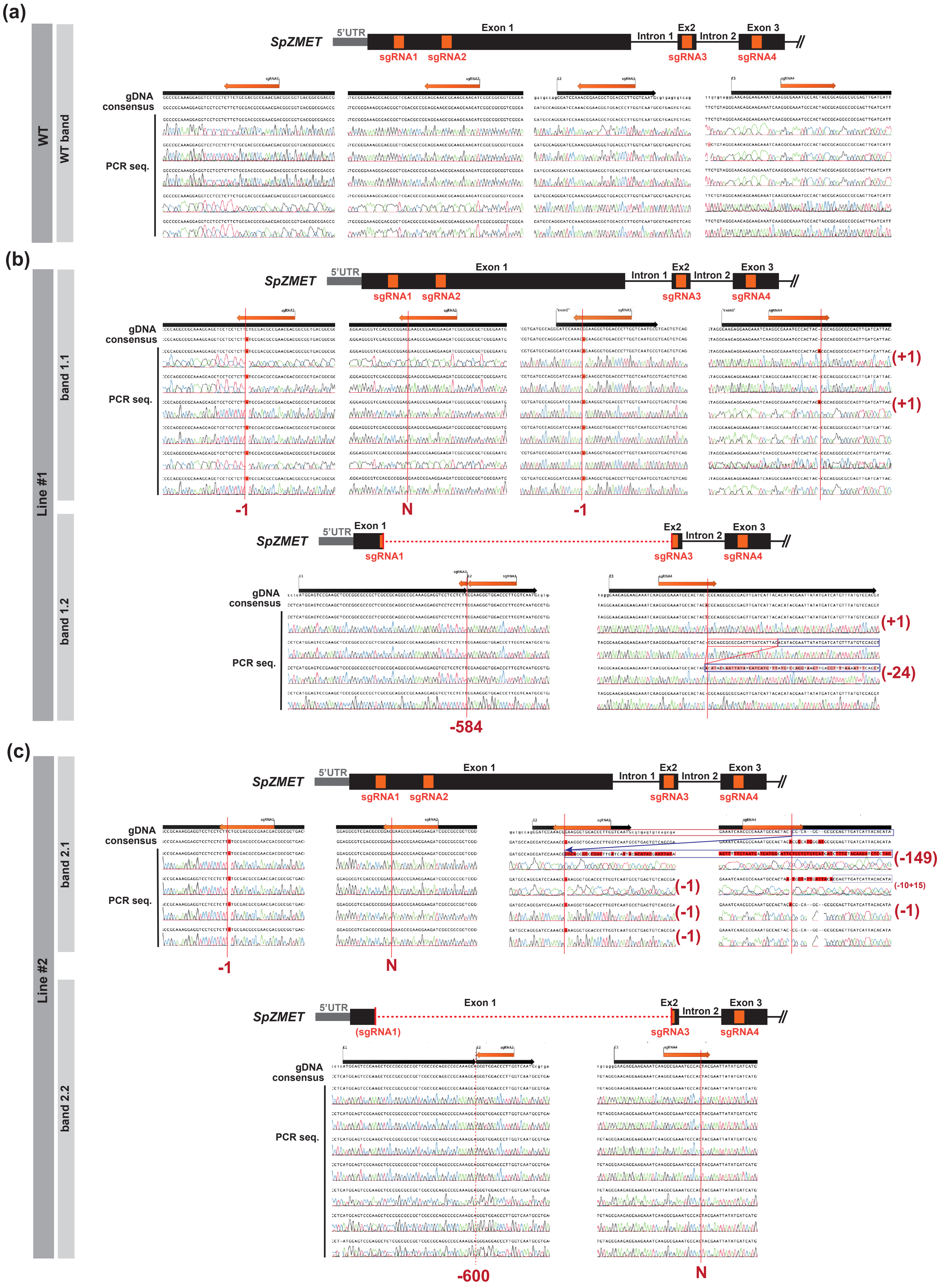


Continues on next page

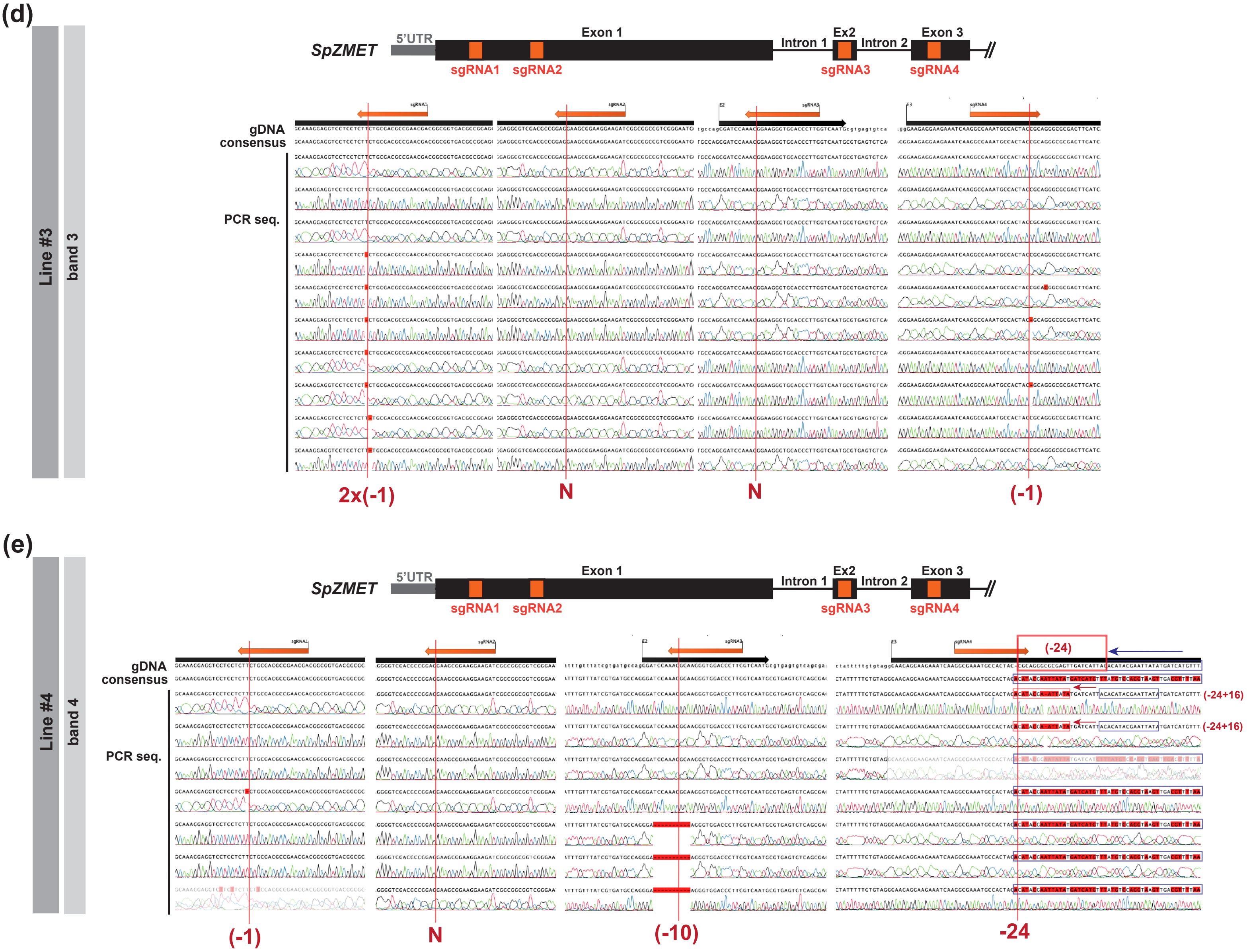


**Figure S8: Analysis of CRISPR/Cas9 induced mutations in *SP162*.**
**(a-e)** Sanger sequencing trace data chromatograms from individual PCR molecules cloned and sequenced from independent regenerated *SP162* lines expressing *zCas9i* from Figure 6. The position of target sequences for each of the four sgRNAs designed is indicated with an orange arrow. Conserved mutations identified across all molecules from the same PCR and the resulting sequence modification are indicated with red numbers. Mutations occurring in some, but not all, molecules are indicated in brackets. N indicates no editing.

### SUPPLEMENTAL FIGURE S9

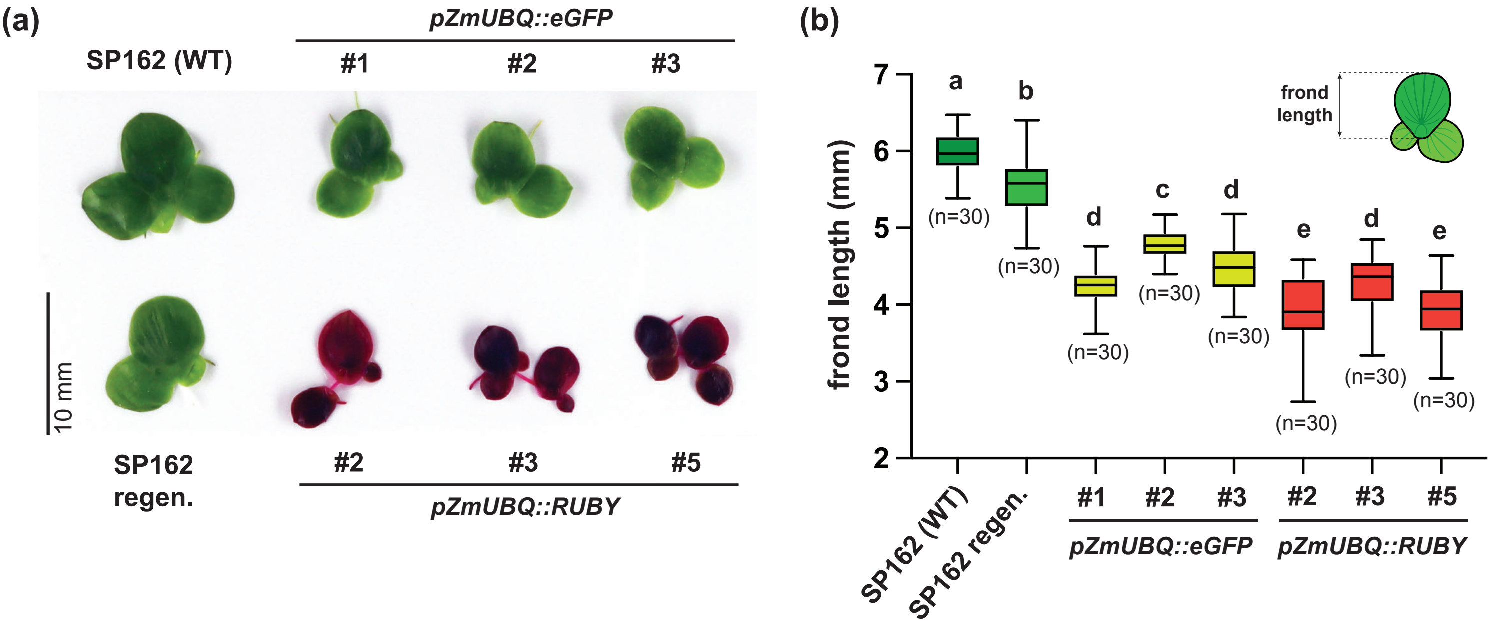


**Figure S9: Frond size variation among regenerated *SP162* lines.**
**(a)** Images of representative fronds of WT, regenerated non-transformed, and three independently regenerated *eGFP* and *RUBY* transgenic *SP162* cultures. **(b)** Mean frond length in mm for each genotype shown in (a). Mean is indicated by solid bar; the boxes extend from the first to the third quartile and whiskers stretch to the furthest value within 1.5 times the interquartile range. Different letters indicate statistically difference at $P<0.05$ in a one-way ANOVA (n=30) statistical analysis.

### SUPPLEMENTAL FIGURE S10

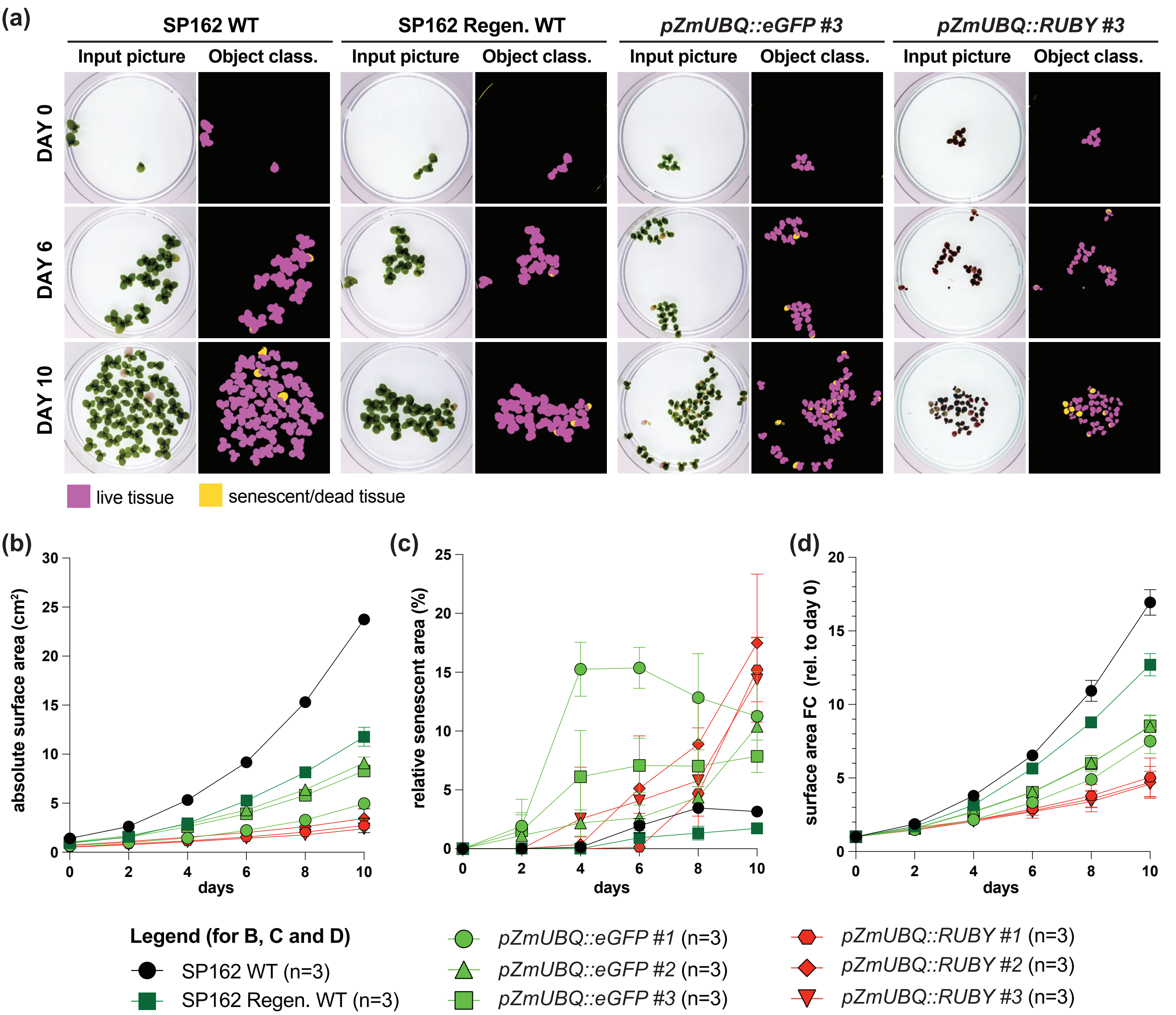


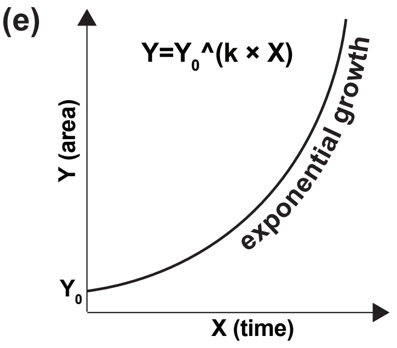


(f)

| | SP162 WT | SP162 Regen WT | *pZmUBQ::eGFP* #1 | *pZmUBQ::eGFP* #2 | *pZmUBQ::eGFP* #3 | *pZmUBQ::RUBY* #2 | *pZmUBQ::RUBY* #3 | *pZmUBQ::RUBY* #5 |
|---|---|---|---|---|---|---|---|---|
| **$R^2$** | 0.9804 | 0.9742 | 0.9808 | 0.9724 | 0.9777 | 0.8481 | 0.8805 | 0.8813 |
| **$Y_0$ (constrained)** | 1 | 1 | 1 | 1 | 1 | 1 | 1 | |
| **k** | 0.2869 | 0.2590 | 0.2008 | 0.2174 | 0.2176 | 0.1645 | 0.1576 | 0.1540 |
| **95% CI (k)** | 0.2818 to 0.2918 | 0.2534 to 0.2644 | 0.1962 to 0.2052 | 0.2119 to 0.2226 | 0.2127 to 0.2224 | 0.1521 to 0.1757 | 0.1474 to 0.1670 | 0.1438 to 0.1632 |
| **Doubling Time (DT)** | 2.416 | 2.676 | 3.452 | 3.188 | 3.185 | 4.212 | 4.397 | 4.502 |
| **95% CI (DT)** | 2.375 to 2.459 | 2.621 to 2.735 | 3.378 to 3.533 | 3.114 to 3.270 | 3.117 to 3.259 | 3.944 to 4.558 | 4.151 to 4.702 | 4.246 to 4.820 |

**Figure S10: Growth analysis of regenerated *SP162* lines.**
**(a)** Input pictures and Ilastik object classification output images of life (magenta) and senescent (yellow) areas of representative points during a 10-days growth time-course of WT, regenerated non-transformed, and three independently regenerated *eGFP* and *RUBY* transgenic *SP162* cultures. **(b)** Absolute area ($cm^2$) 10-day growth curves of WT *SP162* and regenerated lines. **(c)** Relative senescent area (%) to total culture area (live and senescent) in WT *SP162* and regenerated lines during the 10-day time-course. **(d)** Relative area fold change (FC) growth curves, normalized to area of the initial inoculum (day 0) of WT *SP162* and regenerated lines. In (b-d), each point indicates the mean value of three replicates. Error bars represent standard error of the mean. **(e)** Exponential growth curve model. **(f)** Results of non-linear fit of relative growth (d) to an exponential growth model. Doubling time (DT) is indicated in days.

#### SUPPLEMENTAL FIGURE S11

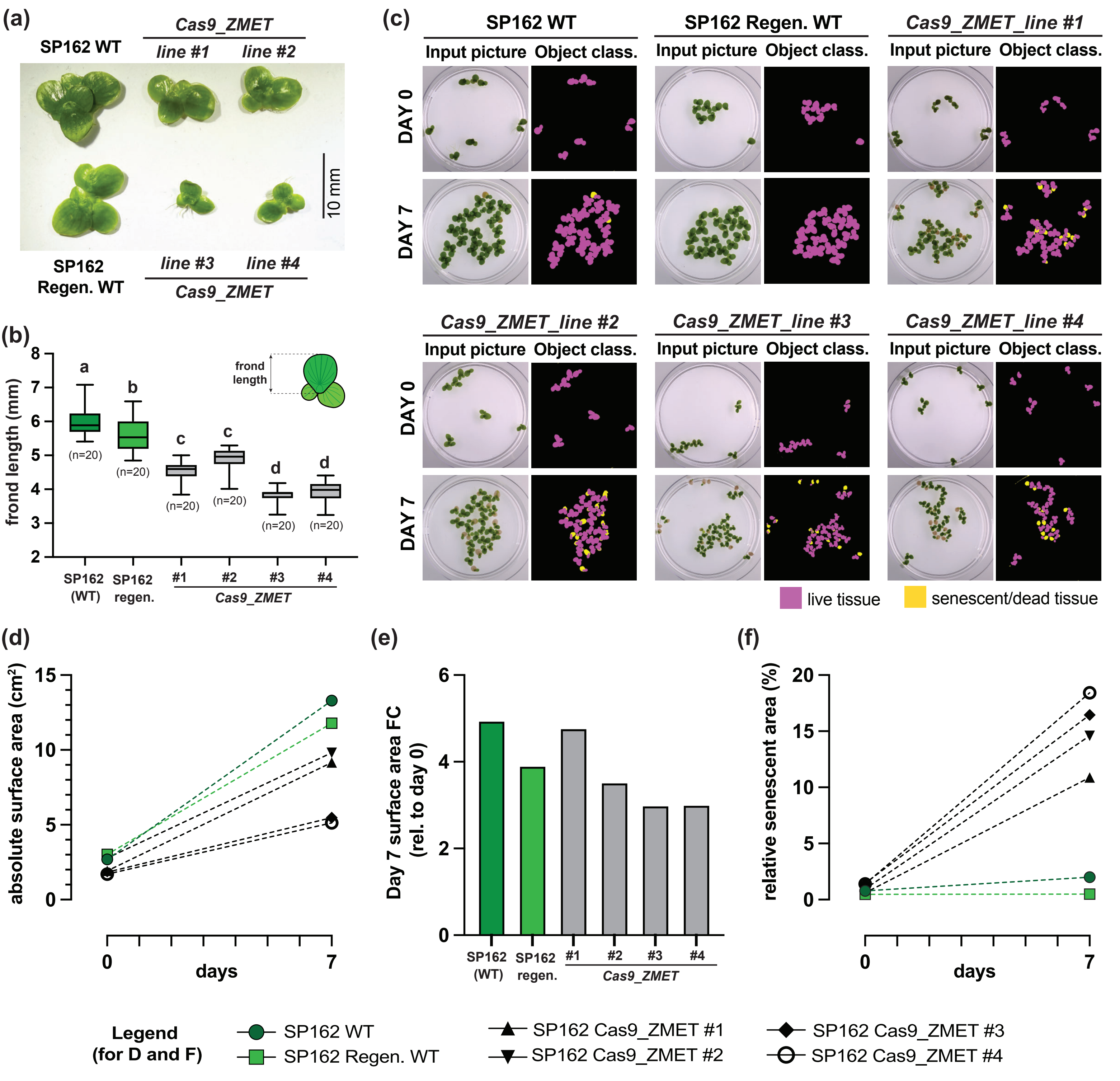


**Figure S11: Phenotypic analysis of regenerated *SP162* lines expressing *zCas9i*.**
**(a)** Images of representative fronds of WT, regenerated non-transformed, and four independently regenerated *zCas9i-2A-eGFP 4x@ZMET* lines analyzed in Figure 6. **(b)** Mean frond length in mm for each genotype shown in (a). Mean is indicated by solid bar; the boxes extend from the first to the third quartile and whiskers stretch to the furthest value within 1.5 times the interquartile range. Different letters indicate statistically difference at $P<0.05$ in a one-way ANOVA (n=20) statistical analysis. **(c)** Input pictures and Ilastik object classification output images of life (magenta) and senescent (yellow) areas at 0 (initial inoculum) and after 7 days of growth of genotypes in (a-b). **(d)** Absolute area ($cm^2$) at 0 and 10 days of the indicated lines. **(e)** Relative area fold change (FC) after 7 days of growth, normalized to area of the initial inoculum (day 0), of WT SP162 and indicated regenerated lines. **(f)** Relative senescent area (%) to total culture area (live and senescent) in WT *SP162* and regenerated lines indicated at 0 and 10 days.

#### SUPPLEMENTAL FIGURE S12

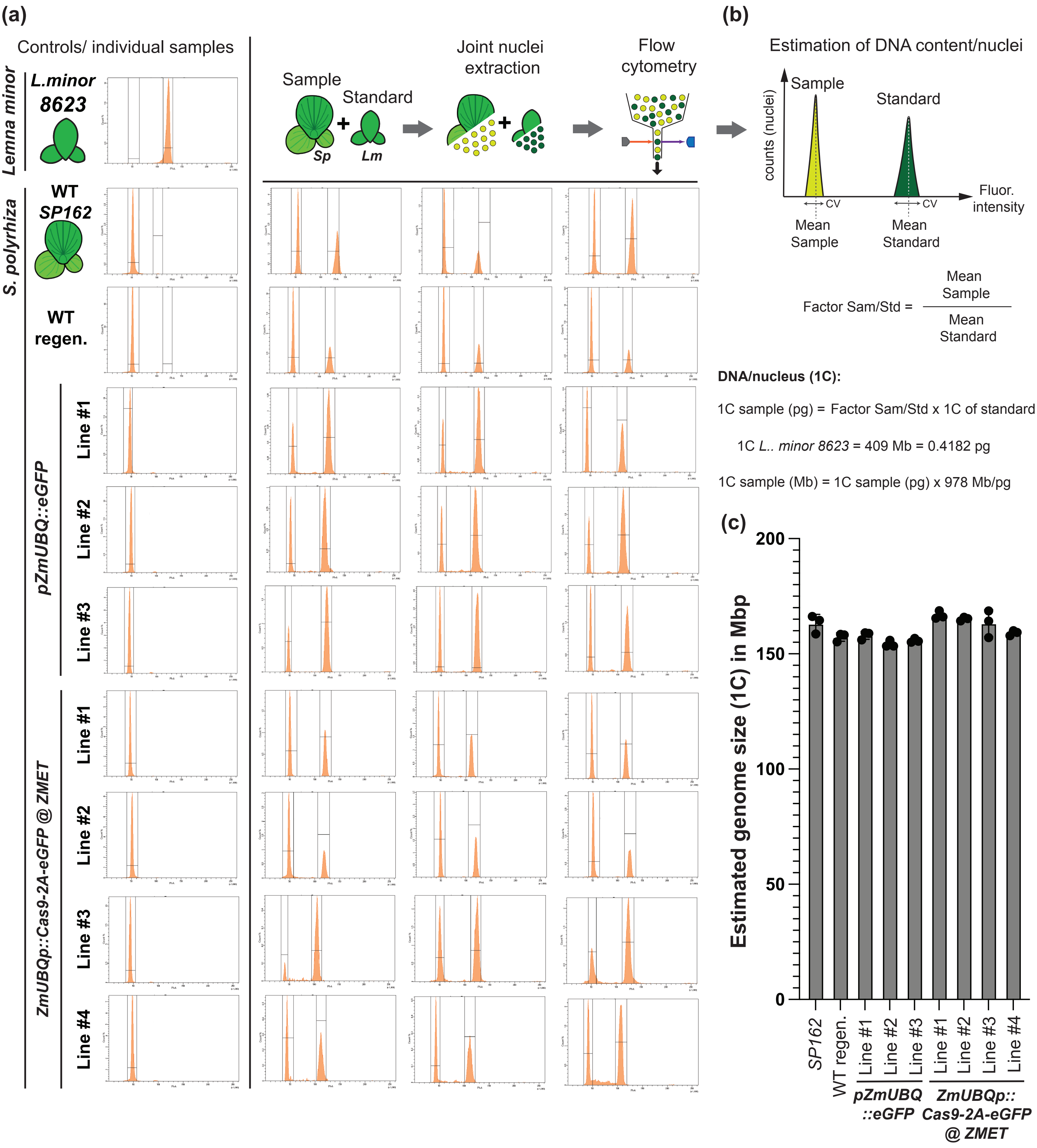


**Figure S12: Flow cytometry measurement of DNA content in regenerated *SP162* lines.**
**(a)** FACS plots of dapi stained nuclei etxracted from individual samples or mixed extraction with the standard (*L. minor #8623*) in triplicates. **(b)** Simple description of the procedure for the estimation of DNA content by FACS. **(c)** Estimated DNA content in each of the lines investigated. Barrs indicate mean values of three independent replicates (dots). Error bars represent the standard deviation of the mean.

#### SUPPLEMENTAL FIGURE S13

##### Transient expression procedure at Mari-Ordonez's Lab

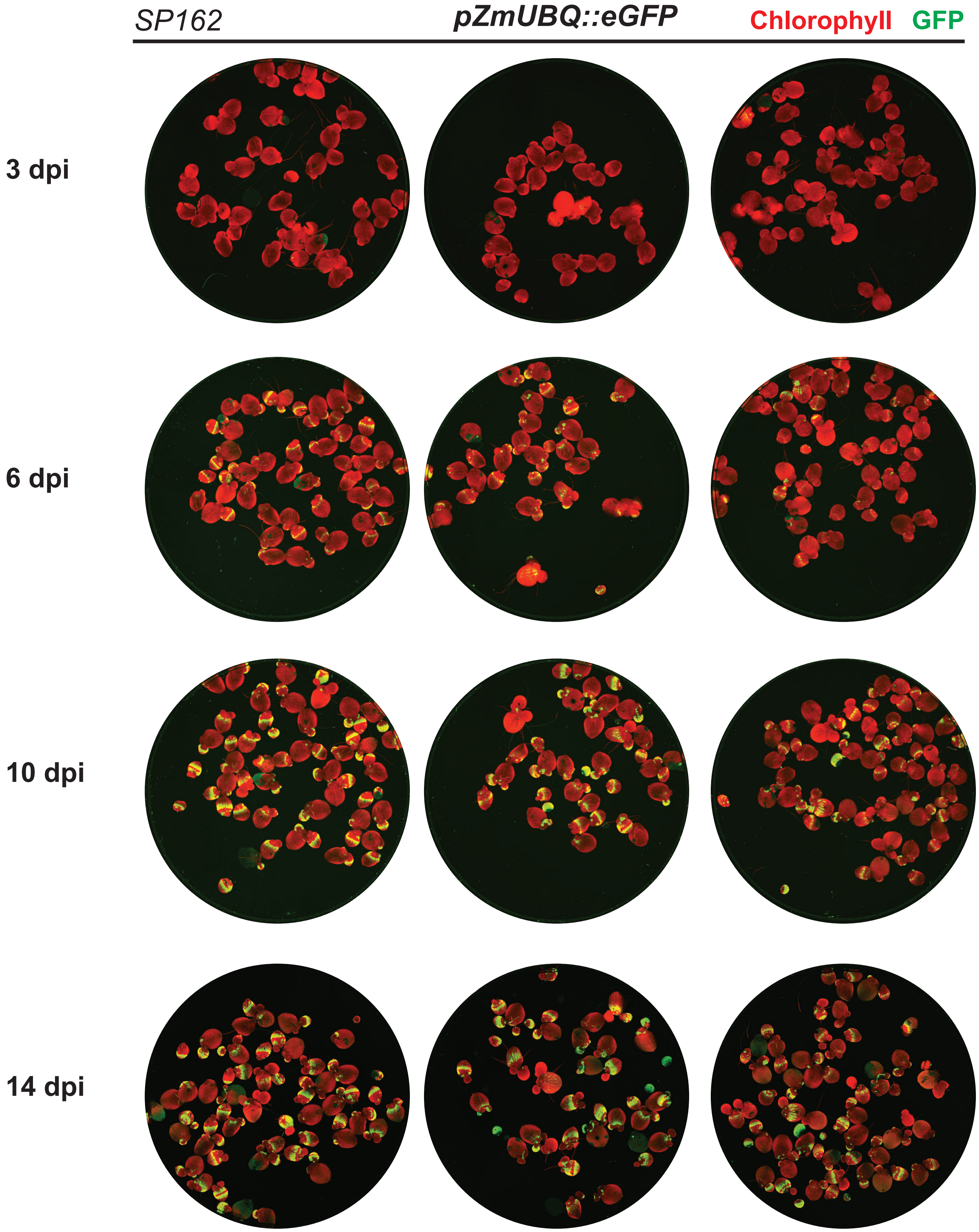


**Figure S13: GFP fluorescent development during transient expression.**
Biomolecular scanning of GFP fluorescence (green) and chlorophyll auto-fluorescence (red) of *SP162* cultures infiltrated with *ZmUBQ::eGFP* at different days post infiltration (dpi) in triplicates.

#### SUPPLEMENTAL FIGURE S14

##### Transient expression procedure at Hubert's Lab

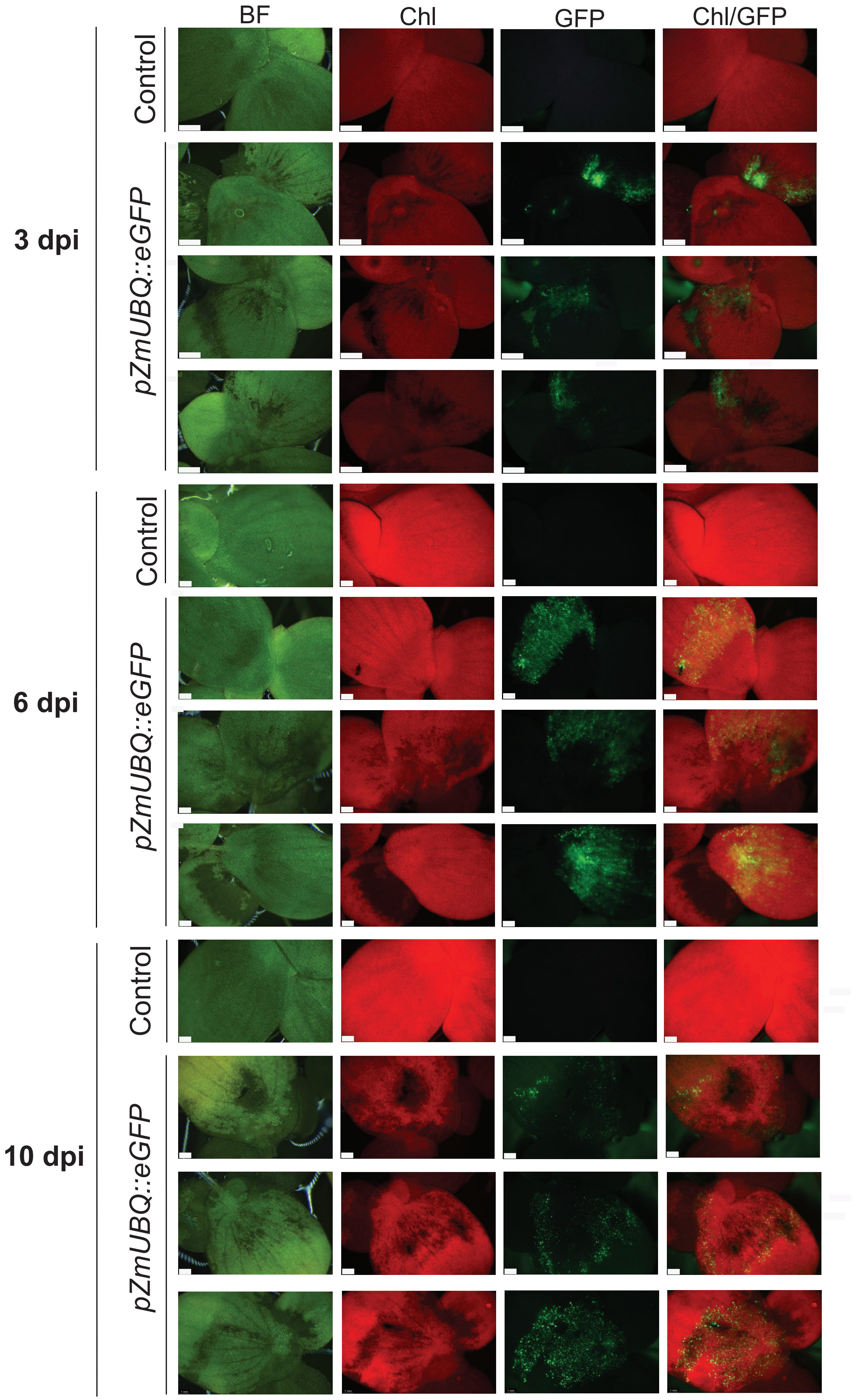


**Figure S14: GFP fluorescent development during transient expression carried in the Hubert's lab.** *Agrobacterium* infiltrated *SP162* fronds displaying transient expression of GFP. The pictures were taken 2, 6 and 10 days post infiltration (dpi). Control, plants without infiltration; BF: Bright field, Chl: chlorophyll autofluorescence.

### SUPPLEMENTAL FIGURE S15

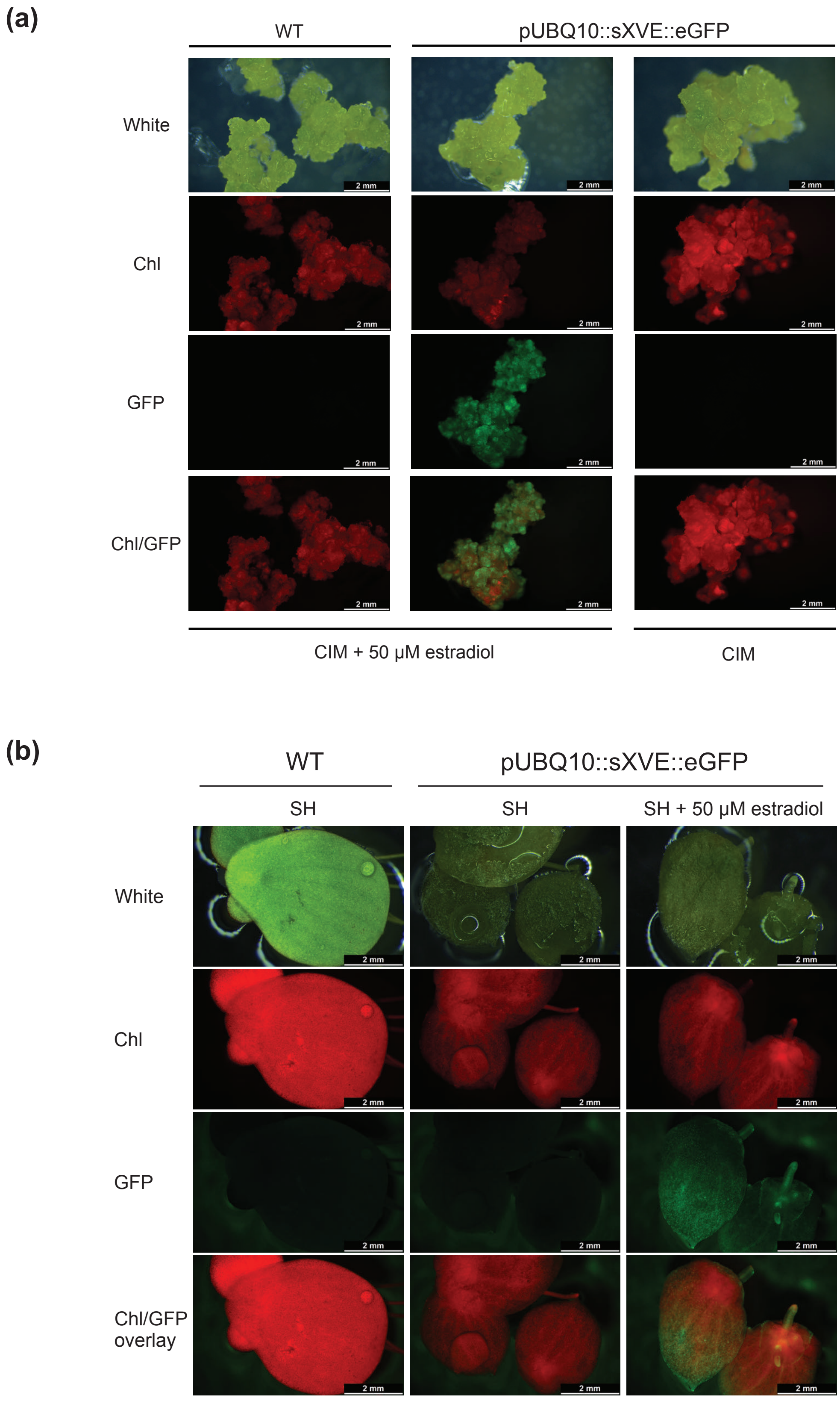


**Figure S15: Stable estradiol-inducible GFP expression in *SP162* calli and regenerated fronds.** Whole transformed *SP162* callus **(a)** and regenerated fronds **(b)** bearing the expression cassette *pUBQ10::sXVE::eGFP.* The pictures were taken after 2days of incubation on CIM or CIM suplemented with 50 µM estradiol under standard callus culture conditions for calli or in SH-mediu, or SH-medium supplemented with 50 µM estradiol. GFP: GFP fluorescence, Chl: chlorophyll autofluorescence, Chl/GFP: Overlay

### SUPPLEMENTAL FIGURE S16

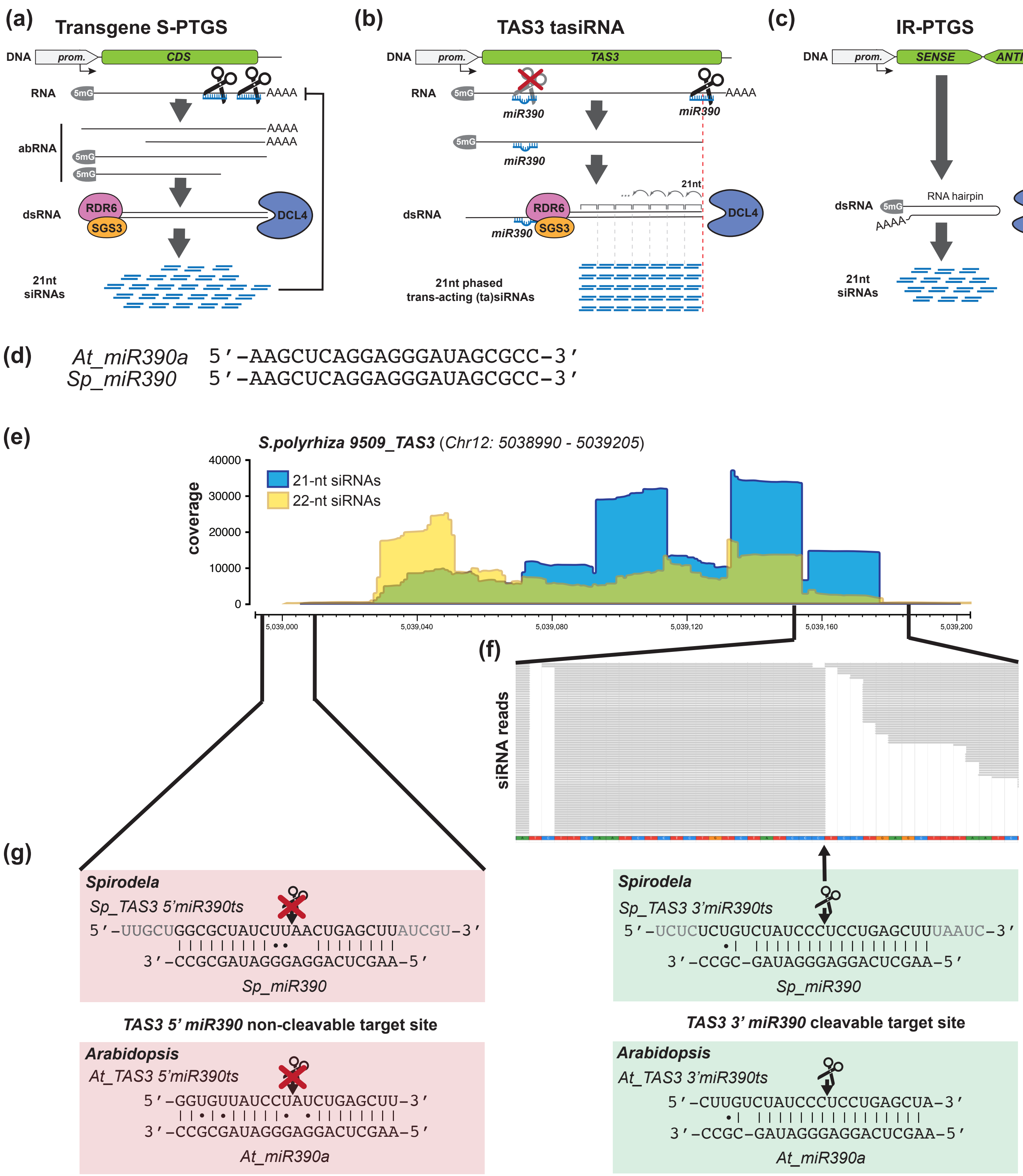


**Figure S16: siRNAs biogenesis in plants and tasiRNAs identification in *S. polyrhiza*.**
**(a-c)** Simplified schematic representation of major DICER-LIKE 4 (DCL4) siRNA biogenesis pathways in flowering plants. **(a)** Transgene induced sense-post-transcriptional gene silencing (S-PTGS) is believed to be initiated by the formation of aberrant (ab)RNA lacking cap or polyA tails due to strong or high expression. Such abRNA serves as substrate to the RNA-DEPENDENT RNA POLYMERASE 6 (RDR6) aided by SUPRESSOR OF RNA SILENCING 3 (SGS3) to produce double-stranded (ds)RNA, processed into 21-nt siRNAs by DCL4. siRNAs then direct silencing complex to cleave and degrade complementary mRNAs. **(b)** Trans-acting (ta)siRNAs originate from dedicated loci that are targeted by micro (mi)RNAs defining the region and initial position to be converted into dsRNA by RDR6, which is processed in a characteristic and precise 21-nt phased fashion by DCL4. In the conserved *TAS3*, the tasiRNA are defined by two miR390 target sites (ts), a cleavable and a non-cleavable due to miss-matches over the catalytic site. **(c)** In the case of inverted-repeat (IR)-PTGS, RDR6 and SGS3 are dispensable for the biogenesis of siRNAs as dsRNA is formed as the result of intramolecular folding of the precursor transcript. **(d)** Sequence conservation between *Arabidopsis* and *Spirodela miR390*. **(e)** 21-22-nt siRNAs coverage at *Spirodela TAS3-tasiRNA*. **(f)** mapped 21-nt siRNAs reads around the predicted miR390 cleavage site. **(g)** alignment of *miR390* at *TAS3* cleavage site in *Spirodela* and *Arabidopsis* for comparison. Lines indicate perfect complementarity; dots, G:U wobbles.
