## Supplemental Methods for "Strain, procedures, and tools for reproducible genetic transformation and genome editing of the emerging plant model *Spirodela polyrhiza* (L.) Schleid"

#### Table of contents:


| Page | |
|---|---|
| 2 | **Supplemental Method S1:** Media composition and preparation. |
| 3 | **Supplemental Method S2:** *Spirodela* frond sterilization. |
| 4 | **Supplemental Method S3:** Turion induction and germination. |
| 5 | **Supplemental Method S4:** Callus induction and culture. |
| 6 | **Supplemental Method S5:** Frond regeneration from callus. |
| 7 | **Supplemental Method S6:** Callus transformation and selection. |
| 9 | **Supplemental Method S7:** Transient expression. |
| 11 | **Supplemental Method S8:** Extended experimental procedures. |
| 20 | **Supplemental Method S9:** List of oligonucleotides. |

### SUPPLEMENTAL METHOD S1: MEDIA COMPOSITION AND PREPARATION

#### *S. polyrhiza* frond culture media

##### Schenk and Hildebrant based

| Schenck and Hildebrandt (SH) medium (Duchefa-Biochemie #S0225) | | |
|---|---|---|
| **Macro-elements** | **[mg/L]** | **[mM]** |
| $CaCl_2$ | 151 | 1.36 |
| $KNO_3$ | 2500 | 24.73 |
| $MgSO_4$ | 195.05 | 1.62 |
| $(NH_4)H_2PO_4$ | 300 | 2.61 |
| **Micro-elements** | **[mg/L]** | **[µM]** |
| $CoCl_2{\cdot}6H_2O$ | 0.1 | 0.42 |
| $CuSO_4{\cdot}5H_2O$ | 0.2 | 0.80 |
| Fe(II)NaEDTA | 19.8 | 53.94 |
| $H_3BO_3$ | 5 | 80.87 |
| KI | 1 | 6.02 |
| $MnSO_4{\cdot}H_2O$ | 10 | 59.17 |
| $Na_2MoO_4{\cdot}2H_2O$ | 0.1 | 0.41 |
| $ZnSO_4{\cdot}7H_2O$ | 1 | 3.84 |
| **Vitamins** | **[mg/L]** | **[µM]** |
| Myo-Inositol | 100 | 554.94 |
| Nicotinic acid | 5 | 1.624 |
| Pyridoxine HCl | 0.5 | 2.43 |
| Thiamine HCl | 5 | 14.825 |

| SH MEDIUM (SOLID) | |
|---|---|
| **SH basalt salt mixture** | 3.2 g/L |
| **Autoclavable additives** | (e.g. sucrose 10 g/L) |
| **Gelrite (recommended) OR** | 4 g/L |
| **Phyto-agar** | 8 g/L |
| **$ddH_2O$ (*)** | Bring to final volume |
| **Adjust pH to** | 5.8 |
| **Autoclave** | 15 min @ 125ºC |
| **Let cool down** | |
| **Non-autoclavable additives** | (e.g. antibiotics)(**) |
| **Pour plates and let solidify** | (**)(***) |

* Depending on tap water hardness precipitates form after autoclave, distilled water is thus recommended.

** Autoclaved plain solid SH media can be stored in bottles at room temperature protected from light and melted by microwaving to be poured or add non-autoclavable additives.

*** Store at 4ºC depending on additives.

| SH MEDIUM (LIQUID) | |
|---|---|
| **SH basalt salt mixture** | 3.2 g/L |
| **Autoclavable additives** | (e.g. sucrose 10 g/L) |
| **$ddH_2O$ (*)(**)** | Bring to final volume |
| **Adjust pH to** | 5.8 |
| **Autoclave** | 15 min @ 125ºC |
| **Let cool down** | |
| **Non-autoclavable additives** | (e.g. antibiotics, acetosyr) |
| **Store** | (**) (***) |

* Depending on tap water hardness precipitates form after autoclave, distilled water is thus recommended.

** Precipitates tend to appear during storge or Spirodela culture.

*** Plain SH medium can be stored at room temperature protected from light. Store at 4ºC depending of additives.

#### N-media based

| Modified N-medium | | | | | |
|---|---|---|---|---|---|
| **STOCK SOLUTIONS (*) (**)** | | | | | **FINAL** |
| **STOCK SOL.** | **Compound** | **[g/L]** | **[mM]** | | |
| **#1** | $KH_2PO_4$ | 27.22 | 200 | | 0.1 mM |
| **#2** | $Ca(NO_3)_2{\cdot}4H_2O$ | 47.23 | 200 | | 1 mM |
| **#3** | $KNO_3$ | 161.8 | 1600 | | 8 mM |
| | $H_3BO_3$ | 0.0618(***) | 1 | | 5 µM |
| | $MnCl_2{\cdot}4H_2O$ | 0.5145 | 2.6 | | 13 µM |
| | $Na_2MoO_4{\cdot}2H_2O$ | 0.018(****) | 0.080 | | 0.4 µM |
| | $MgSO_4{\cdot}7H_2O$ | 49.30 | 200 | | 1 mM |
| **#4** | Fe(II)NaEDTA | 1.835 | 5 | | 25 µM |

* Filter sterilise each of the four stock solutions. DO NOT AUTOCLAVE!

** Keep all stock sol. at 4ºC. If Sol. #3 might forms crystal precipitates, warm up in water bath at 37-40ºC before use.

*** start from a 10 mM stock, add 100 mL/L to Stock sol.#3

**** start from a 4 mM stock, add 20 mL/L to Stock sol.#3

| N MEDIUM (LIQUID) | |
|---|---|
| **$ddH_2O$ (*)** | Final volume - stock solutions vol. (**) |
| **Stock solution #1** | 0.5 mL/L |
| **Stock solution #2** | 5 mL/L |
| **Stock solution #3** | 5 mL/L |
| **Stock solution #4** | 5 mL/L |
| **Adjust pH to** | 5.8 |
| **Autoclave** | 15 min @ 125ºC |
| **Let cool down** | |
| **Store** | (***) |

* Depending on tap water hardness precipitates form after autoclave, distilled water is thus recommended.

** Mix stock solutions into water, mixing concentrated stocks precipitate

*** Plain N medium can be stored at room temperature protected from light. Store at 4ºC depending of additives.

#### *S. polyrhiza* tissue culture media

##### Murashige and Skoog based

| Murashige and Skoog (MS) medium (Duchefa-Biochemie #M0245) | | |
|---|---|---|
| **Macro-elements** | **[mg/L]** | **[mM]** |
| $CaCl_2$ | 332.02 | 2.99 |
| $KNO_3$ | 1900 | 18.79 |
| $MgSO_4$ | 180.54 | 1.5 |
| $(NH_4)NO_3$ | 1650 | 20.61 |
| $KH_2PO_4$ | 170 | 1.25 |
| **Micro-elements** | **[mg/L]** | **[µM]** |
| $CoCl_2{\cdot}6H_2O$ | 0.025 | 0.11 |
| $CuSO_4{\cdot}5H_2O$ | 0.025 | 0.10 |
| Fe(II)NaEDTA | 36.70 | 100 |
| $H_3BO_3$ | 6.20 | 100.27 |
| KI | 0.83 | 5 |
| $MnSO_4{\cdot}H_2O$ | 16.90 | 100 |
| $Na_2MoO_4{\cdot}2H_2O$ | 0.25 | 1.03 |
| $ZnSO_4{\cdot}7H_2O$ | 8.60 | 29.91 |
| **Vitamins** | **[mg/L]** | **[µM]** |
| Myo-Inositol | 100 | 554.94 |
| Nicotinic acid | 0.5 | 4.06 |
| Pyridoxine HCl | 0.5 | 2.43 |
| Thiamine HCl | 1 | 2.965 |

| Hormone Stock solutions (2,4-D, TDZ, Zeatin) | |
|---|---|
| **Dissolve in 1M KOH:** | 1 mL (1/10th final vol) |
| **Thidiazuron (TDZ) OR** | 10 mg (1 mg/mL final) |
| **2,4-Dichlorophenoxyacetic acid (2,4-D) OR** | 22.1 mg (2.21 mg/mL final) |
| **Zeatin (Zea)** | 10 mg (1 mg/mL final) |
| **Dissolve** | (*) |
| **$ddH_2O$** | 9 mL (bring to final volume) |
| **Filter sterilise** | |
| **Aliquote** | 1 mL/aliquote (**) |

* Dissolve with vigorous shaking. Don't add water until is fully dissolved.

** Store TDZ and Zeatin at -20ºC, 2,4-D at 4ºC.

#### CALLUS INDUCT. & REGEN.

| CIM: CALLUS INDUCTION MEDIUM (SOLID) | |
|---|---|
| **MS basalt salt mixture** | 4.4 g/L |
| **2-(*N*-Morpholino)ethanosulfonic acid monohydrate (MES)** | 0.5 g/L |
| **Sucrose** | 10 g/L |
| **Polyvinylpyrrolidone (PVP)-40** | 0.1 g/L |
| **Citric acid** | 50 mg/L |
| **2,4-Dichlorophenoxyacetic acid (2,4-D)** | 0.221 mg/L (1 µM) (*) |
| **Gelrite** | 4 g/L |
| **Adjust pH to** | 5.8 |
| **Autoclave** | 15 min @ 125ºC |
| **Let cool down** | |
| **Thidiazuron (TDZ)** | 0.4 mg/L (1.82 µM) (**) |
| **Pour plates and let solidify** | (***) |

* From a filter sterilised 2.21 g/L 2,4-D stock stored at 4ºC

** From a filter sterilised 1 g/L TDZ stock stored at -20ºC

*** Store plates @ 4ºC for up to 1 month

| FRM: FROND REGENERATION MEDIUM (SOLID) | |
|---|---|
| **MS basalt salt mixture** | 4.4 g/L |
| **2-(*N*-Morpholino)ethanosulfonic acid monohydrate (MES)** | 0.5 g/L |
| **Sucrose** | 10 g/L |
| **Gelrite** | 4 g/L |
| **Adjust pH to** | 5.8 |
| **Autoclave** | 15 min @ 125ºC |
| **Let cool down** | |
| **Thidiazuron (TDZ) OR** | 1 mg/L (4.54 µM) (*) |
| **Zeatin (Zea)** | 2,2 mg/L (10 µM) (**) |
| **Pour plates and let solidify** | (***) |

* From a filter sterilised 1 mg/mL TDZ stock stored at -20ºC

** From a filter sterilised 1 mg/mL Zea. stock in water stored at -20ºC

*** Store plates @ 4ºC for up to 1 month

#### CALLUS TRANSF. & SELECT.

| CIM-AGROBACT. CO-CULTURE MEDIUM (SOLID) | |
|---|---|
| **SOLID CIM MEDIA** | |
| **After autoclave** | **and TDZ addition** |
| **Acetosyringone** | 100 µM (*) |
| **Pour plates and let solidify** | (**) |

* From a 100 mM Acetosyringone stock in DMSO stored at -20ºC

** Prepare fresh. Store plates @ 4ºC for no longer than 1 week.

| CIM-AGROELIMINATION MEDIUM (SOLID) | |
|---|---|
| **SOLID CIM MEDIA** | |
| **After autoclave** | **and TDZ addition** |
| **Cefotaxime (Cef)** | 0.30 mg/mL (*) |
| **Ticaricillin (Tic)** | 0.25 mg/mL (**) |
| **Pour plates and let solidify** | (***) |

* From a filter sterilised 300 mg/mL Cef. stock in water stored at -20ºC

** From a filter sterilised 250 mg/mL Tic. stock in water stored at -20ºC

*** Store plates @ 4ºC for up to 1 month.

| CIM-SELECTION MEDIUM (SOLID) | |
|---|---|
| **SOLID CIM MEDIA** | |
| **After autoclave** | **and TDZ addition** |
| **Cefotaxime (Cef)** | 0.30 mg/mL (*) |
| **Ticaricillin (Tic)** | 0.25 mg/mL (**) |
| **Hygromycin (Hyg) (*)** | 5 (low) or 10 (high) mg/L (**) |
| **Pour plates and let solidify** | (***) |

* adjust weak (low) and strong (high) selection for each antibiotic or selection marker

** From a filter sterilised 40 mg/mL Hyg. stock in water stored at -20ºC

*** Store plates @ 4ºC for up to 1 month.

#### *S. polyrhiza Agrobacterium* infection media

| AGROINFECTION PRETREATMENT MEDIUM (LIQUID) (*) | |
|---|---|
| **L-glutamate** | 20 mg/L |
| **Filter sterilise** | **(**)** |

* Only required for vacuum infiltration of agrobacterium

** Sterile media can be stored at 4ºC

| AGROINFECTION MEDIUM (LIQUID) | |
|---|---|
| **$MgCl_2$** | 10 mM |
| **Sucrose** | 50 g/L |
| **Filter sterilise** | **(*)** |
| **Acetosyringone** | 150-200 µM (**) (***) |
| **Silwet L-77** | 0.02% (****) |

* sterile media can be aliquoted and frozen at -20ºC without acetosyr.

** From a 100 mM Acetosyringone stock in DMSO stored at -20ºC. Use media immediately do not store with acetosyr.

*** Use 200 µM for calli or 150 µM for frond transformation

**** Only for vacuum frond infiltration of agrobacterium. Add only right before vacuum.

### SUPPLEMENTAL METHOD S2: SPIRODELA FROND STERILIZATION

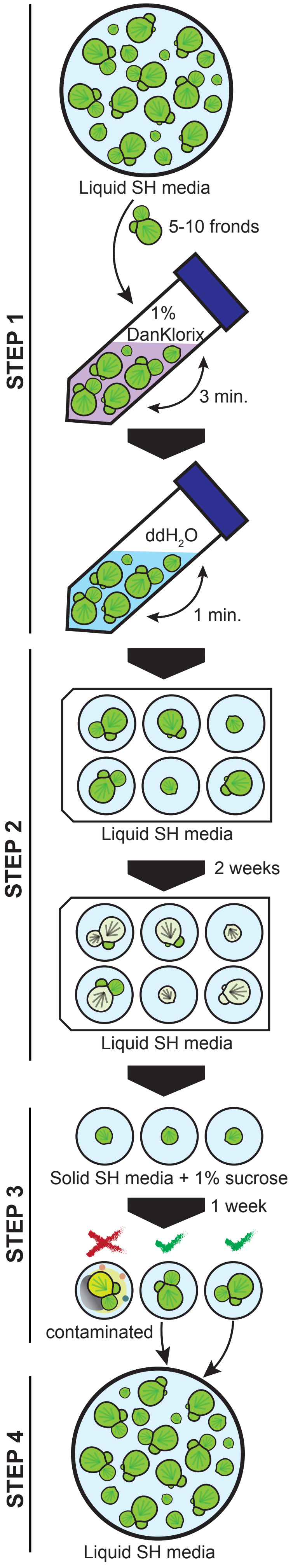


#### STEP 1: FROND STERILIZATION.

**NOTE 1:** Work always under sterile conditions in sterile hood.

**1.1)** Starting from a Spirodela liquid culture, transfer 5-10 fronds (including attached daughters) to a 50 mL sterile test tube.

**1.2)** Add 10 mL of a 1% NaClO solution from a DanKlorix dilution (comercial 2.8% w/v NaClO-based desinfectant) in sterile ddH$_2$O.

**1.3)** Agitate/invert gently for 3 minutes.

**1.4)** Aspirate/remove desinfectant solution and add 40 mL of sterile ddH$_2$O.

**1.5)** Agitate/invert gently for 1 minute.

**1.6)** Aspirate/remove water and repeat wash twice with 30 mL sterile ddH$_2$O.

**1.7)** After last wash, pour plants in water into a sterile 10 cm petri dish. Repeat with as many batches of Spirodela fronds as desired.

#### STEP 2: FROND RECOVERY.

**2.1)** Transfer individual fronds to wells in a 6-well plate containing 5 ml of SH-media. Seal with Leucopore and transfer to growth chamber.

**NOTE 2:** Change or re-sterilize handling equipment (forceps, loops...) between fronds to prevent contamination from non fully sterilized fronds.

**2.2)** Grow fronds for 2 weeks. Fronds exposed to desinfectant will bleach and some daughter fronds will emerge green.

#### STEP 3: TEST FOR MICROBIAL GROWTH.

**3.1)** Transfer the content of each well to individual 3.5 or 10 cm petri dishes with solid (0.8% w/v agar) SH-media with 1% (w/v) sucrose. Seal with Leucopore and transfer to growth chamber.

**3.2)** After a week, check plates for microbial contamination growth.

**NOTE 3:** If contamination appears in all samples:
- At step 1.3) agitate/invert for 6 min (less surviving fronds), or
- At step 1.2) use 2% DanKlorix dilution and apply vacuum for 5 minues and/or,
- At step 2.1) add 200mg/L cefotaxime to liquid SH media.

#### STEP 4: AMPLIFICATION.

**4.1)** Transfer fronds from plates showing no signs of microbial contamination to containers/flasks with SH media, keep in sterility and refresh media every 1-2 weeks.

### SUPPLEMENTAL METHOD S3: TURION INDUCTION AND GERMINATION

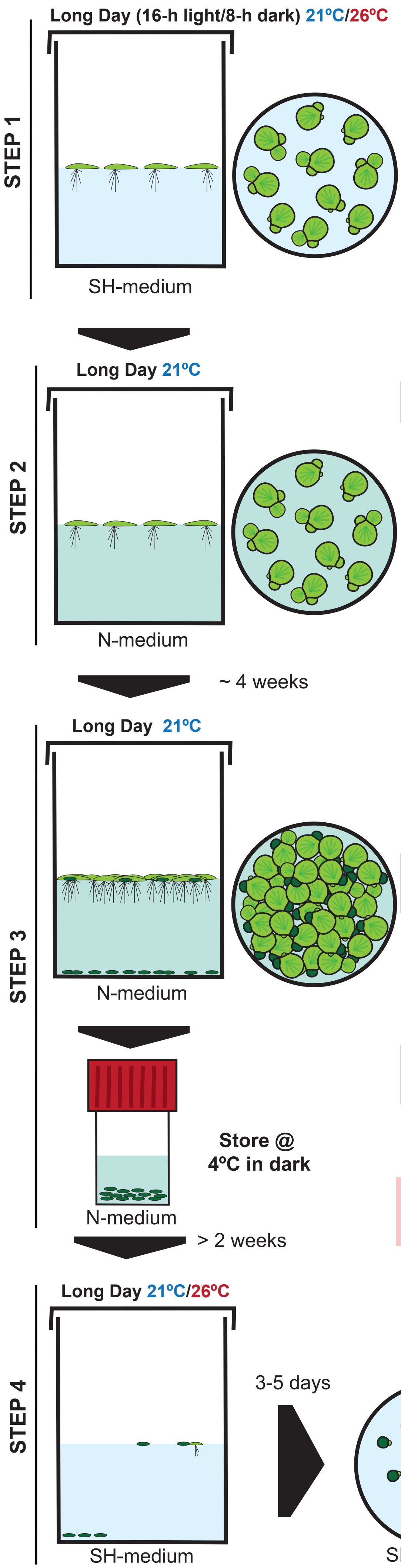


##### STEP 1: CULTURE EXPANSION.

**1.1)** Grow a Spirodela culture in liquid SH-medium to 80%, or more, confluence in long day conditions at 21ºC.

**NOTE 1:** Turion induction can be performed in various containers (beakers, Erlenmeyers, Magenta vessels...). We found induction in containers allowing for large volumes (0.5-1L) to be most effective and requiere less culture maintenance.

##### STEP 2: TURION INDUCTION.

**2.1)** Transfer the whole culture to liquid N-medium at 21ºC. and let proliferate to saturation without splitting the culture.

**NOTE 2:** Turion induction is faster and more homogeneous at 15ºC.

**2.2)** Change media every 3-4 weeks. Transfer the whole culture to fresh liquid N-media.

**2.3)** After about 4 weeks in N-media turions should start to appear at the budding pockets of Spirodela fronds.

**2.4)** Keep the cultures until turions sink to the bottom of the container.

##### STEP 3: TURION HARVEST AND STORAGE.

**3.1)** At each media replacement, after fronds have been transfer to fresh media, discard most of the media without disturbing the turions. Pass the remaining media with turions through a sterile metalic sieve to harvest the turions.

**NOTE 3:** The Spirodela culture can be maintained in N-media to keep harvesting turions.

**3.2)** Transfer ccollecccted turions to a Gosselin screw-cap container (50 mL Falcon tubes are also suitable) containing liquid N-medium and store at 4ºC in darkness.

**NOTE 4:** Turions can be kept for several months, even a year, in cold without any great loss of germination viability.

##### STEP 4: TURION GERMINATION.

**NOTE 5:** Turion dormancy needs about 2-3 weeks of stratification during cold storage to be released and allow high rates of turion germination.

**4.1)** Transfer a group of turions from storage to liquid SH-medium in the desired container and culture them under long day conditions at 21 or 26ºC. Turions should emerge to the surface in the first 1-2 days and initiate germination with first frond appearing within 3-5 days.

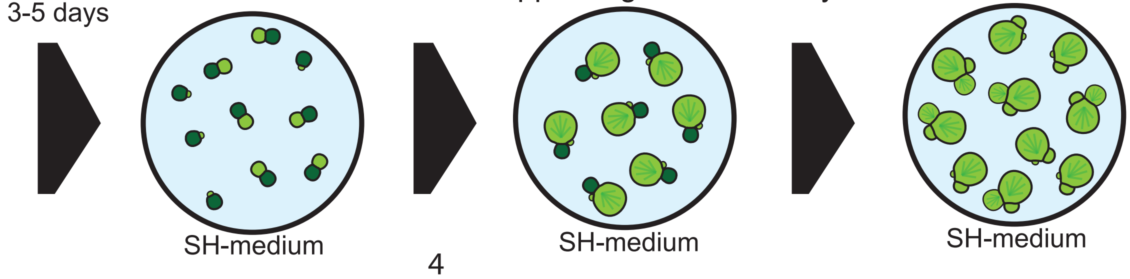

### SUPPLEMENTAL METHOD S4: CALLUS INDUCTION AND CULTURE

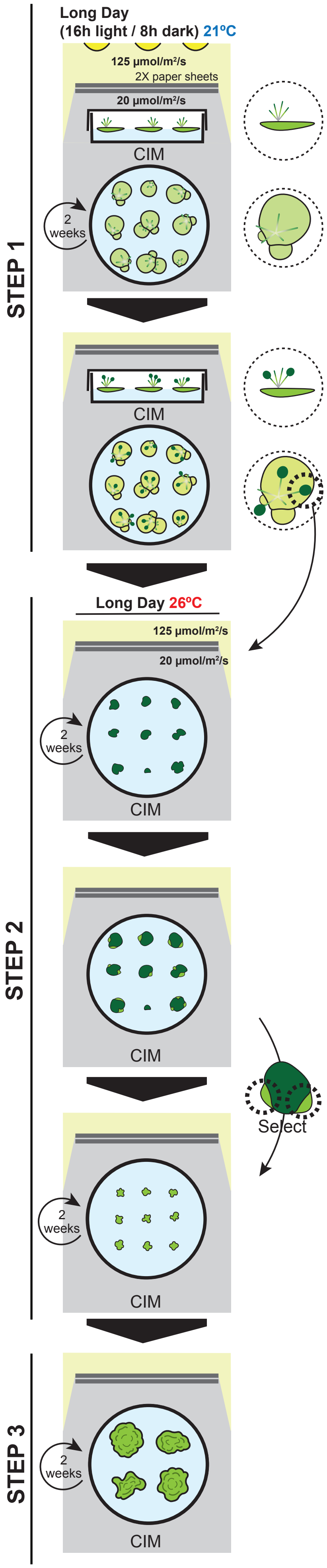


#### STEP 1: CALLUS INDUCTION.

**NOTE 1:** Work always under sterile conditions in sterile hood.

**1.1)** Starting from a Spirodela liquid culture, transfer 10-20 fronds on their dorsal side (roots up) to 10 cm plates containing 40-50 mL of solid CIM.

**NOTE 2:** Genlty press the fronds against the media to ensure as much contact surface between the fronds and the media.
**NOTE 3:** Follow several plates in case of contamination.

**1.2)** Culture plates in a growth chamber at 21ºC under long day conditions (18h light / 6 h darkness) under shade (light intensity ~20 µmol/m²/s) by covering the plates with two layers of white paper.

**NOTE 4:** Keep fronds and calli under shade all all time from now on. Low light prevents browning, wich negatively impacts calli growth and survability.

**1.3)** Transfer fronds to plates with fresh solid CIM every 2 weeks.

**1.4)** Keep subculturing fronds in solid CIM until the formation of ~2-3 mm long compact dark green calli from root tips (or meristematic pockets in the case of callus induction from previously regenerated plants).

#### STEP 2: FRIABLE CALLUS CULTURE.

**2.1)** Detach/separate calli from fronds and transfer to solid CIM (20-40 calli per plate).

**2.2)** Transfer calli to plates with fresh solid CIM every 2 weeks. Culture calli at 26ºC under long day conditions (18h light / 6 h darkness) under shade (light intensity ~20 µmol/m²/s).

**NOTE 5:** During callus subculture, avoid passing browning or necrosing tissue/sectors at every passage as they can negatively affect callus growth.

**2.3)** During calli subculture, select sectors that appear pale green (not white) to transfer into fresh plates as they appear when they are 2-3 mm in size.

**2.4)** Transfer calli to plates with fresh solid CIM every 2 weeks. Untill they form pale green, fast growing and highly friable callus that easily disgregate when applying gentle pressure with forceps.

#### STEP 3: CALLUS MAINTENANCE AND AMPLIFICATION.

**3.1)** Subculture pale green friable calli every two-three weeks.

**NOTE 6:** culture periods longer than 3 weeks leads to browning.

**3.2)** For callus amplification for downstream uses (e.g. transformation), transfer small calli fragments (2-3 mm) to several plates one to two weeks before their use. Do not overgrow.

### SUPPLEMENTAL METHOD S5: FROND REGENERATION FROM CALLUS

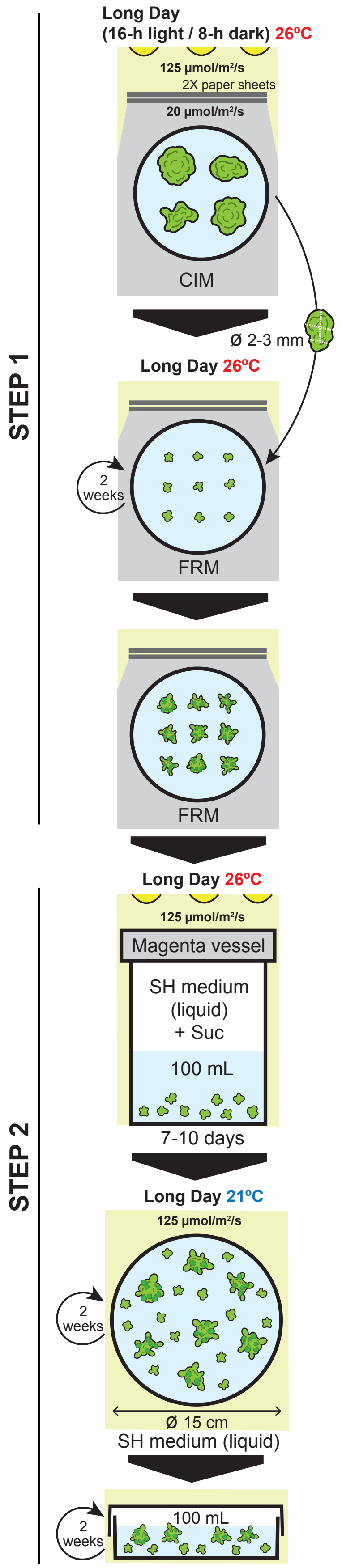


#### STEP 1: INITIATION OF REGENERATION.

**1.1)** Dissect calli into small pices (2-3 mm ø) and transfer into solid frond regeneration media (FRM). Culture at 26ºC under long day light regime shadowed with 2 layers of paper.

**NOTE 1:** Use fast growing calli that had been transfered to fresh CIM 1-2 weeks in advance.
**NOTE 2:** Prepare several FRM plates with ~16 calli/plate.
**NOTE 3:** Regeneration can be tested using TDZ or Zeatin. In our hands Zeatin has provided more consisten initiation, but it might depend on local conditions.

**1.2)** Transfer fronds to plates with fresh solid FRM every 2 weeks.

**NOTE 4:** Friable calli tend to fragment easily. Transfer always as much calli fragments as possible avoiding browning parts if they appear.

**1.3)** Keep subculturing calli into FRM until first signs of regeneration appear.

**NOTE 5:** Initiation of regeneration usually starts after 4 weeks and is generally characterized by the apparition of small (<1 mm) darker green, triangular frond primordia appearing individually or in clusters that further develop into "tongue-like" leafy structures.
**NOTE 6:** Transgenic calli tend to take longer to initiate regeneration than non-transformed calli.

#### STEP 2: ACCLIMATATION TO AUTOTROPHIC GROWTH IN LIQUID.

**2.1)** Transfer calli to 100 mL of liquid SH medium containing 10 g/L of sucrose in Magenta vessels.

**2.2)** Culture regenerating calli for 7-10 days at 26ºC under long day conditions (18h light / 6 h darkness) without shade (light intensity ~125 µmol/m²/s).

**NOTE 7:** As regeneration takes long time in sucrose containing media, antibiotics can be added to media (0.25 mg/mL ticarcillin + 0.30 mg/mL cefatoxime)

**2.3)** Transfer calli to 100 mL of liquid SH medium without sucrose in 15 cm ø petri dishes or culture plates.

**2.4)** Culture regenerating calli at 21ºC under long day conditions (18-h light / 6-h darkness) without shade (light intensity ~125 µmol/m²/s).

**2.5)** Refresh medium every 2 weeks.

**NOTE 8:** Icreased buoyancy in calli with regenerating fronds will make them float.
**NOTE 9:** Once sucrose is removed, regeneration can also be continued at 26ºC. If possible grow in both and monitor growth and development to choose the best condition.

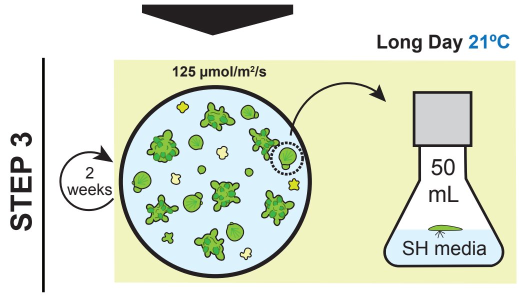


#### STEP 3: END OF REGENERATION.

**3.1)** As independent fronds emerge, trasfer them to infividual flasks containing SH medium and culture them at 21ºC under long day conditions.

**NOTE 10:** We noticed better development of regenerated fronds at 21ºC, once a stable population has been stablished it can be kept at 21ºC or transfered to 26ºC for faster growth.

### SUPPLEMENTAL METHOD S6: CALLUS TRANSFORMATION AND SELECTION

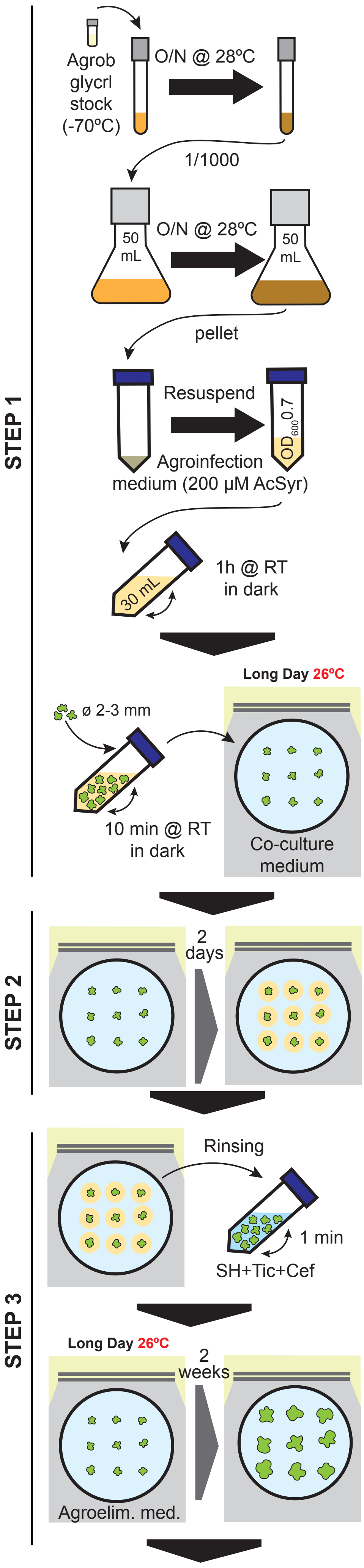


Continues next page

**STEP 1: CALLI INFECTION WITH AGROBACTERIUM.**

**1.1)** From agrobacterium glycerol stock, grow an O/N 2 mL Agrobacterium pre-culture in LB with appropiate antibiotics at 28ºC.

**1.2)** Inoculate a 20-50 mL LB with appropiate antibiotics with 1 µL/mL agrobacterium preculture. Grow O/N at 28ºC.

**1.3)** Measure the $OD_{600}$ and pellet the necessary culture volume for 20-15 min at 3000-5000x *g* at room temp. (RT) and to obtain a final 30 mL at $OD_{600}$ 0.7 bacterial suspension in agroinfection medium (10 mM $MgCl_2$, 5% sucrose, 200 µM acetosyringone).

**1.4)** Incubate the bacterial solution protected from light with gentle agitation for 1h at room temperature.

**1.5)** During bacteria incubation time, dissect calli into at least 20-40 small pices (2-3 mm ø) and leave them in liquid SH medium in a wide opening container for easy access (e.g. 50 mL test tube).

**NOTE 1:** Use small calli that had been transfered to fresh CIM 1-2 weeks in advance.

**1.6)** Remove SH medium and add the 30 mL of bacterial suspension to the calli. Incubate for 10 min with mild agitation at room temperature in dark.

**STEP 2: CALLI-AGROBACTERIUM CO-CULTURE.**

**2.1)** Remove bacterial suspension and/or transfer the calli to a clean empty petri dish.

**2.2)** Transfer calli to 10 cm ø dishes containing 50 mL of coculture media (solid CIM + 100 µM acetosyringone).

**NOTE 2:** Prepare at least 3 plates with ~16 calli/plate.

**2.3)** Co-culture for 2 days at 26ºC under long day light regime shadowed with 2 layers of paper.

**NOTE 3:** Agrobacterium will grow on calli and on the media around them.

**STEP 3: AGROBACTERIUM ELIMINATION AND RECOVERY.**

**3.1)** Transfer calli to 50 mL test tubes containing 30 mL liquid SH media + 0.3 mg/mL Cefotaxime + 0.25 mg/mL Ticarcillin. Rinse calli by inverting the tube for 1 minute.

**3.2)** Pour the content of the tube into an empty petri dish and transfer calli to plates with solid agroelimination medium (solid CIM + 0.3 mg/mL Cefotaxime + 0.25 mg/mL Ticarcillin).

**NOTE 4:** Transgene expression might already be detected after co-culture.

**3.3)** Let calli recover in CIM-agroelimination medium for 2 weeks at 26ºC under long day light regime shadowed with 2 layers of paper.

**NOTE 5:** If using a fluorescent or visible reporter, expression can be followed to assess transformation efficiency before selection.
**NOTE 6:** A significant fraction of transgene expression might be transient and lost during calli subculture.

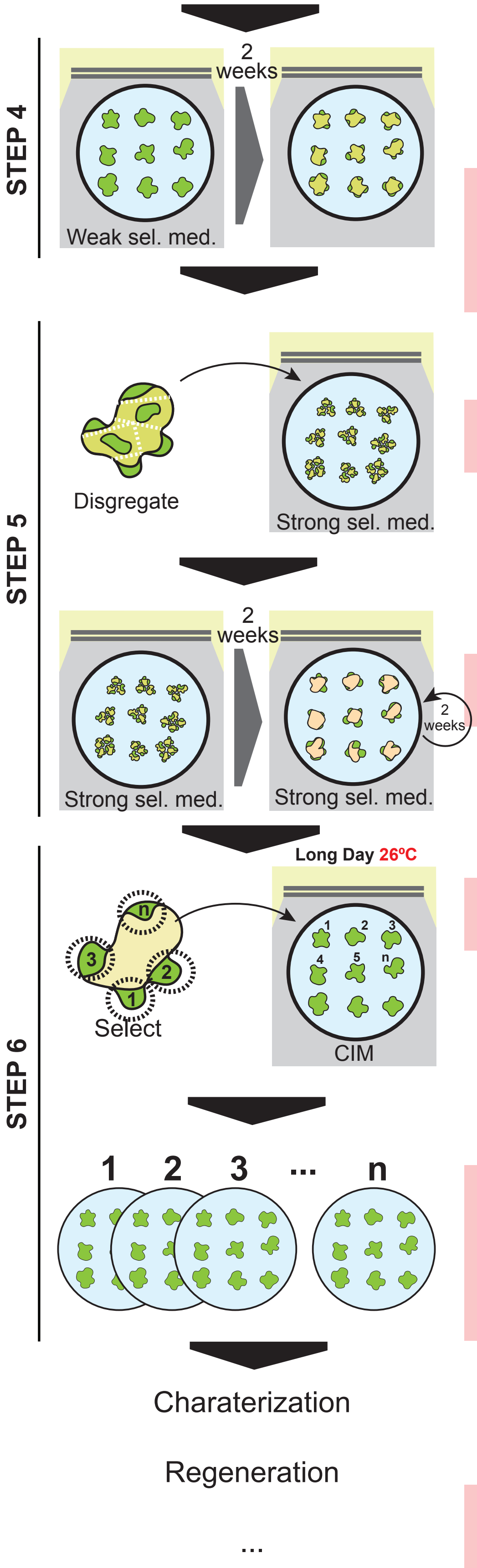


**STEP 4: WEAK SELECTION.** ⚠ **(see table below for antibiotic guidelines)**

**4.1)** Transfer calli to weak selection medium (solid CIM + 0.3 mg/mL Cefotaxime + 0.25 mg/mL Ticarcillin + low dose selection/antibiotic).

**NOTE 7:** Weak selection should allow transgenic cells to stablish a growing cell polpulation and slow the growth of non-transgenic cells without causing widespread death and browning, which can negatively affect (and cause the loss) of trasngenic cells.

**4.2)** Maintain in weak selection media for two weeks.

**NOTE 8:** Depending on selection type, non-transgenic cells/calli will become paler/yellow.

**STEP 5: STRONG SELECTION.** ⚠ **(see table below for antibiotic guidelines)**

**5.1)** Transfer calli to strong selection medium (solid CIM + 0.3 mg/mL Cefotaxime + 0.25 mg/mL Ticarcillin + high dose selection/antibiotic).

**NOTE 9:** Using forceps/loop, disgregate/spread calli on medium when transfering to increase contact of tissue with selection media.

**5.2)** Keep on strong selection media, transfering to new strong selection plates every 2 weeks, until clear healthy pale green sectors are visible to naked eye over yellow/pale brown calli.

**NOTE 10:** At every media change,avoid transfering calli sectors displaying excessive/dark browning.

**STEP 6: TRANSGENIC CALLI AMPLIFICATION.**

**6.1)** Select and trasfer ONLY resistant, pale green (if visible/fluorecent marker used, check expression) calli sectors to CIM media at 26ºC under long day light regime shadowed with 2 layers of paper.

**NOTE 11:** Mark/label every resistant calli sector independently as they might originate from independent transformation events. Although chimeric calli are also quite likely to ocurr, characterization and identification of independent transgenic events are facilitated downstream if already separated at this stage.

**6.2)** Transfer calli to fresh CIM plates every 2 weeks untill fast growing, heathy pale green friable calli has been obtained for further processes (e.g. regeneration).

**NOTE 12:** If non-trasngenic calli (e.g. non fluorecent) is suspected to reapear or not have been properly selected against, amplify calli maintaining strong selection.
**NOTE 13:** If agrobacterium regrows in CIM following removal of Tic and Cef, add both antibiotics back to CIM (agroelimination medium).

#### ANTIBIOTIC SELECTION GUIDELINES

| ANTIBIOTIC | WEAK SELECT. | STRONG SELECT. | NOTES |
|---|---|---|---|
| HYGROMYCIN | 5 mg/L | 10 mgL | Calli are very sensitive to hygromycin. Easy to see resistant sectors. |
| SPECTINOMYCIN | 5 mg/L | 10 mgL | Growth not completely arrested at low concentrations, however they bleach easily. To facilitate identification of resistant sectors is recommended to break down and spread calli when transferring to strong selection. |
| G418 | 25 mg/L | 50 mgL | |
| KANAMYCIN | 50 mg/L | 100 mgL | |

### SUPPLEMENTAL METHOD S7: TRANSIENT EXPRESSION.

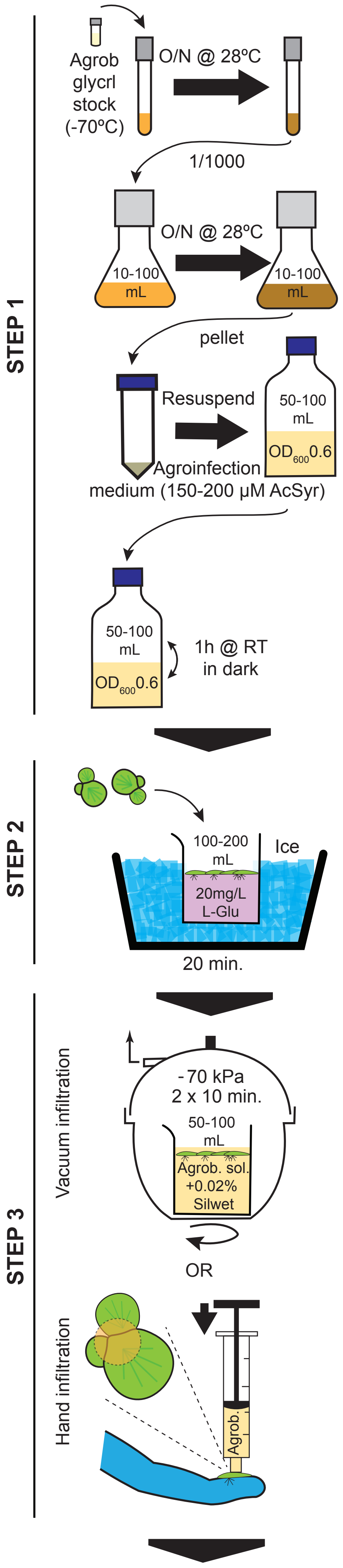


Continues next page

**STEP 1: PREPARATION OF AGROBACTERIUM CELLS.**

**1.1)** From agrobacterium glycerol stock, grow an O/N 2-5 mL Agrobacterium pre-culture in LB with appropiate antibiotics at 28ºC.

**1.2)** Inoculate a 10-100 mL LB with appropiate antibiotics with 1 µL/mL agrobacterium preculture. Grow O/N at 28ºC.

**1.3)** Measure the $OD_{600}$ (measure the OD of a 1:10 dilution) and pellet the necessary culture volume for 10-15 min at 3000 - 5000x *g* at room temp. (RT) and to obtain a final 50-100 mL at $OD_{600}$ 0.6 bacterial suspension in agroinfection medium (10 mM $MgCl_2$, 5% sucrose, 150-200 µM acetosyringone).

**NOTE 1:** Do not let agrobacterium culture to grow over $OD_{600}$ 3, efficiency of transformation is negatively affected if the original culture is too dense before diluting it.

**1.4)** Incubate the bacterial solution protected from light with gentle agitation for 1h at room temperature.

**1.5)** During bacteria incubation time, pre-treat plants (Step 2).

**STEP 2: PLANT PRE-TREATMENT.**

**2.1)** Transfer Spirodela fronds (20+) to a 0.5-1L beaker on ice containing 100-200 mL of a prechilled (on ice or at 4ºC), filter sterilized, 20mg/L L-glutamine solution.

**NOTE 2:** Best results are obtained when using a rapidly growing 50-80% confluent Spirodela culture grown in large media volumes (0.5-1L) 2-5 days after refreshing SH-medium.
**NOTE 3:** Do not store glutamine for more than 1 month as it degrades into toxic ammonia at a rate of 0.1-0.2% / day at 4ºC.

**2.2)** incubate the plants on the L-glu solution on ice for 20 min.

**STEP 3: AGROBACTERIUM INFILTRATION.**

**3.1)** To the bacteria solution add Silwet L-77 to a final 0.02% (v/v) concentration.

**NOTE 4:** For co-infiltration of several agrobacterium/plasmids mix bacteria solutions in equal proportions before infiltration.

<u>OPTION 1</u>: VACUUM INFILTRATION.
**3.2a)** Transfer pretreated plants to a 0.5L beaker containing the agrobac. solution. Apply vacuum (-70 kPa) twice for 10 min. gently releasing vacuum in between. Gently swirl the vacuum chamber at regular intervals.

**NOTE 5:** Duckweed stomata are only located at the dorsal (upper) side of the fronds. Swirling facilitates infiltration of the bacteria solution ensuring the upper surface of the fronds is covered in it.

<u>OPTION 2</u>: HAND INFILTRATION.
**3.2b)** Infiltrate individual fronds manually with a needless 1mL syringe. Aim at the dorsal side of the frond, where budding pockets are located. Position the syringe perpendicular to the frond and ensure there are no air bubbles between the bacteria solution and the frond. Apply pressure firmly to infiltrate the frond.

**NOTE 6:** Apply the right amount of pressure to infiltrate the duckweed (infiltrated area becomes darker and translucent). Too much presure results in tissue damage which negatively affects transformation efficiency.

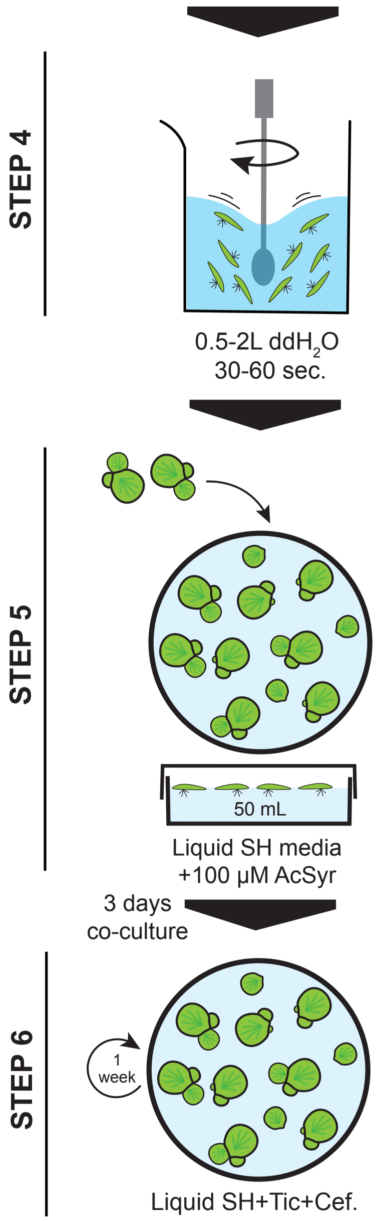


**STEP 4: WASHING OFF AGROBACTERIUM SOLUTION.**

**4.1)** Transfer infitrated fronds to a beaker containing 0.5-2L of $ddH_2O$ at room temperature. Stir for 30-60 seconds to wash off any remaining agrobacterium infitration solution from the surface of the plants.

**4.2)** Repeat wash.

**STEP 5: PLANT-AGROBACTEIUM CO-CULTURE.**

**5.1)** Transfer Spirodela fronds to 10 cm ø petri dish plant culture plates containing 30 mL of liquid SH-media + 100 µM acetosyringone. Seal with leucopore or para-film.

**NOTE 7:** Add fronds to no more than 1/3 or 1/2 of the plate surface to allow the culture to grow. Spirodela slows down growth in dense or very confluent cultures, which significantly decreases transformation efficiency. Split in several plates if needed.

**5.2)** Culture the plants for 3 days at 21ºC under long day conditions.

**STEP 6: AGROBACTERIUM ELIMINATION AND PLANT CULTURE.**

**6.1)** Transfer plants to plates cointaing 30 mL SH-medium with 0.3 mg/mL Cefo-taxime + 0.25 mg/mL Ticarcillin.

**6.2)** Refresh media (SH-medium with 0.3 mg/mL Cefotaxime + 0.25 mg/mL Ticarcillin) every week.

**NOTE8:** Depending on promoter used, expression of transgenes can be observed as early as after co-culture and increases in the following days.

#### SUPPLEMENTAL METHODS S8: Extended experimental procedures.

***SP162* turion induction, storage and germination.**

*SP162* was cultured in liquid SH-medium until 50-80% confluence. The culture was then transferred to liquid N-medium and let grow under long day conditions at 21ºC till turions started appearing at the bottom of the container. Turions were harvested weekly passing the media through a sieve after transferring the culture to fresh medium. Turions were stored in darkness at 4ºC in N-medium in Gosselin screw cap containers (Corning, TPC30C-002). Turions were germinated in liquid SH-medium at 21ºC under long-day conditions. For turion germination assays germination, as those with a frond emerging, were counted in 2-3 biological replicates from turions stored at 4ºC for different periods. Media and step-by-step procedure can be found in **Method S1, S3**.

**Molecular cloning.**

All pre-existing plasmids directly used or source of sequences for cloning in this study are publicly available from Addgene (https://www.addgene.org): *pZmUBQ:RUBY* (#160909) (He et al. 2020) was used for transformation and as source for *pZmUBQ*, *P2A* and *HygR* (*NptII*) cloning; *eGFP* sequence was amplified from *pTKan-p35S::Csy4-pNOS::cogGFP* (#110145) (Liang et al. 2017); the *KanR* cassette was amplified from *pNK3071* (#219755) (Shakhova et al. 2024); zCas9i was cloned from pDGE666 (#153231) (Stuttmann et al. 2021); *pYLCRISPR/Cas9Pubi-H* (#66187), which served as template for the amplification of the 2x *BsaI* cloning cassette, and the vectors template for direct Golden Gate sgRNA cloning under *pOsU3m, pOsU6a, pOsU6a/LacZ, pOsU6b or pOsU6c* correspond to #66187, #66193, #66194, #66195, #66196 and #66197 respectively (Ma et al. 2015). In addition, the *PDK* intron was cloned from *pKANNIBAL* (NovoPro #V010838) (Wesley et al. 2001).

Destination plasmids for Golden Gate Assembly: Plasmids for constitutive expression compatible with Golden Gate cloning with two different plant selectable markers (hygromycin or kanamycin) were generated by introducing 2 *BsaI* sites positioned between a *ZmUBQ* promoter and a *35S* terminator: *pZmUBQ_2x_BsaI_35St_HygR* and *pZmUBQ_2x_BsaI_35St_KanR*. Briefly, a *BsaI* restriction site was removed from the *pZmUBQ* by PCR from a plasmid harboring an expression cassette for *RUBY* and *HygR* (Addgene, #160909). Subsequently, the *RUBY* coding sequence and terminator were excised by digestioin with *BcuI* (Thermo Scientific, #FD1253) and *HindIIII* (Thermo Scientific, # FD0505). A cassette containing two *BsaI* sites upstream of a 35S terminator, obtained by digesting a PCR product amplified with primers containing *BcuI* and *HindIII* sites, was ligated into the digested vector using T4 DNA ligase (Thermo Scientific, #EL0021) to generate *pZmUBQ_2x_BsaI_35St_KanR*. The resulting plasmid was digested with *HindIII* and *MluI* (Thermo Scientific, # FD0564) to remove the Hygromycin resistance cassette. A PCR fragment containing an expression cassette for Kanamycin resistance in plants, amplified from (Addgene #219755) using oligos including *HindIII* and *Mlu*I restriction sites, was digested and ligated into the vector using T4 DNA ligase generating *pZmUBQ_2x_BsaI_35St_HygR*.

Constitutive and inducible expression plasmids: The destination vectors pZmUBQ_*2x_BsaI_35St_HygR* and *pZmUBQ_2x_BsaI_35St_KanR* were used for Golden Gate Assembly of various CDSs, amplified by PCR using oligos including *BsaI* restriction sites. For *pZmUBQ_eGFP* the *eGFP* coding sequence was amplified from (Addgene #110145). For *pZmUBQ_FP:PFhp* the last 322 bp of *eGFP* were amplified from (Addgene #110145) to generate a forward and a reverse complementary sequence. The *PDK* intron was amplified from pKANNIBAL (V010838, NovoPro) using flanquing primers containing *BsaI* sites and cloned between the two *GFP* fragments.

To assembly the plasmid *pZmUBQ::eGFP_SpecR*, first the *HptII* gene was removed from *pZmUBQ::eGFP_HygR* by PCR and replaced by *SpcN* gene, which confers resistance to spectinomycin. The *SpcN* gene was codon-optimized for nuclear expression in monocotyledons and fused to an improved chloroplast transit peptide (Shen et al. 2017) and acquired via gene synthesis (GeneArt Strings DNA Fragments, Invitrogen). The insert was ligated into the backbone plasmid by using NEBuilder® HiFi DNA Assembly (NEB, #E2621) protocol.

To assembly the plasmid *pUBQ10::sXVE::eGFP*, the estradiol-inducible system (including the *sXVE* expression cassette and *LexA* promoter) was PCR amplified from the plasmid *pGPTVII.hyg*. Then, the *ZmUBQ* promoter was removed from *pZmUBQ::eGFP_HygR* via PCR and replaced by the estradiol-inducible promoter using NEBuilder® HiFi DNA Assembly (NEB, #E2621) protocol.

To assembly the plasmid *pZmUBQ::DsRed2*, the *eGFP* was removed from *pZmUBQ::eGFP* by using *SpeI* and *BsrGI* (New Englad Biolabs) digest and ligated to *DsRed2* sequence previously PCR amplified from *pGreen-DsRed2* vector (gently provided by Prof. Andreas Wachter, JGU Mainz) using NEBuilder® HiFi DNA Assembly (NEB, #E2621) protocol.

Constitutive zCas9i_P2A_GFP, sgRNA multiplex system cloning: For *pZmUBQ_zCas9i_P2A_GFP,* the *zCas9i* was amplified from (Addgene #153231) (pDGE666) and the *eGFP* from (Addgene #110145). After cloning these two fragments by Golden Gate into *pZmUBQ_2x_BsaI_35St_HygR*, the resulting plasmid was digested with *MluI*. A PCR fragment containing 2 *BsaI* sites, amplified by PCR from *pZmUBQ_2x_BsaI_35St*, using oligos including *MluI* sites, was digested and inserted by T4 DNA ligation. In addition, A set of Oligonucleotide- based direct sgRNA cloning plasmids was constructed for use as donors in Golden Gate Assembly containing diverse RNApol III promoters and a sgRNA scaffold (*sgRNAscf*): *pOsU3m_sgRNAscf*, *pOsU6a_sgRNAscf*, *pOsU6aLacZ_sgRNAscf*, *pOsU6b_sgRNAscf*, *pOsU6c_sgRNAscf*. Initially, the Addgene 66193 plasmid was digested with *BsaI* (New England Biolabs, # R3733S) to eliminate the RNApol III promoter, the sgRNA scaffold and an internal *BsaI* site in the ampicillin resistance gene. A modified ampicillin resistance gene lacking the *BsaI* site was amplified by PCR and cloned into the digested vector using T4 DNA ligase. The resulting plasmid was then digested *HindIII* and *XhoI*, and a fragment containing sgRNA scaffold with an upstream *BsaI* site was cloned by T4 DNA ligase as a digested PCR fragment amplified with oligonucleotides with *HindIII* and *XhoI*. The intermediate plasmid with the *BsaI_sgRNA* scaffold was subsequently used to clone, by T4 DNA ligase, various RNA pol III promoters, each with a downstream *BsaI* site introduced by PCR. The promoters were PCR-amplified from the following Addgene plasmids described in Ma et al., *Molecular Plant*, 2015: *pOsU3m* from Addgene 66193, *pOsU6aLacZ* from Addgene 66195, *pOsU6b* from Addgene 66196 and *pOsU6c* from Addgene 66197.

sgRNA cloning: The oligonucleotide- based direct sgRNA cloning plasmids described above were used for the insertion of custom-designed sgRNA. Each plasmid was digested by *BsaI* Oligonucleotides (19-20 nt) corresponding to specific sequences immediately upstream of a protospacer adjacent motif (PAM) in the genomic region of *SpZMET* were designed. Oligo design was performed according to (Ma et al. 2015). Forward oligonucleotides were synthesized with four additional nucleotides at the 5′ end, corresponding to the overhang generated after *BsaI* digestion downstream of the RNA polymerase III promoter. The reverse complementary oligonucleotides were synthesized with a 3′ terminal “caaa” sequence, matching the overhang produced upstream of the sgRNA scaffold following *BsaI* digestion. Oligonucleotide pairs were annealed and phosphorylated using T4 PNK (Thermo Scientific, #EK0031), then cloned into the *BsaI* digested plasmids using T4 DNA ligase. The resulting plasmids were: pOsU3m_sgRNA4_sgRNAscf, pOsU6a_sgRNA1_sgRNAscf, pOsU6aLacZ_ sgRNA3_sgRNAscf, pOsU6c_ sgRNA2_sgRNAscf.

For the final *pZmUBQ_zCas9i_P2A_GFP_4x_sg_ZMET*, 4 amplicons containing different sgRNAs specific for *SpZMET* were amplified by PCR with primers containing *BsaI* sites in both ends following the primer combinations described in Ma et al., *Molecular Plant*, 2015 for 4x sgRNA cloning from *pOsU3m_sgRNA4_sgRNAscf*, *pOsU6a_sgRNA1_sgRNAscf*, *pOsU6aLacZ_sgRNA3_sgRNAscf*, *pOsU6c_sgRNA2_sgRNAscf* as described below, were cloned by Golden Gate Assembly.

All plasmids were transformed into chemically competent *E.coli* DH5a and confirmed by SANGER sequencing by the Molecular Biology Service facility of the Vienna BioCenter or at LCG Genomics before transformation into *Agrobacterium*.

##### ***S. polyrhiza*** **162 whole genome sequencing and assembly**

High-molecular-weight (HMW) genomic DNA was prepared by first isolating nuclei from fresh plant tissue, followed by extraction with the Promega Wizard HMW DNA Extraction Kit (protocol followed from step 3 onward). Briefly, ~100 mg tissue was combined with 1.5 mL nuclei extraction buffer in a gentleMACS M Tube and mechanically dissociated at 4 °C using a gentleMACS Dissociator. The homogenate was passed through a 40-µm cell strainer and centrifuged at 1,000 × g for 6 min to pellet nuclei. The supernatant was discarded and the pellet was gently resuspended in 500 µL Promega HMW Lysis Buffer using wide-bore tips. DNA extraction then followed the kit protocol from step 3 onward, except that nuclei derived from 100 mg tissue were used. DNA from eight samples was pooled and further purified using an AMPure XP bead cleanup (1:1 ratio). DNA concentrations were measured with a DeNovix spectrophotometer and a Qubit fluorometer. The Oxford Nanopore (ONT) sequencing library was prepared using the Nanopore SQK-LSK114 Kit following the ONT protocol, and sequencing was performed in a PromethION flow cell (v10.4.1 FLO-PRO114M). Base calling of the raw data was performed using Dorado v0.9.5 with model dna_r10.4.1_e8.2_400bps_sup@v5.0.0. The Nanopore reads were then filtered using NanoFilt v2.8.0 to retain the reads with average base quality ≥Q90 and length ≥5 Kb (Coster et al. 2018) .

The filtered ONT reads were used for error correction and *de novo* genome assembly, followed by Clair3 variant calling and Racon polishing inside the PECAT v0.0.3 program, resulting in 30 linear contigs for haplotype 1 (Nie et al. 2024) . This assembly was then further polished with Pilon v1.23 using the previously available Illumina short-read data for the *SP162* strain (filtered using Trimmomatic v0.39 with parameters “LEADING:10 TRAILING:10 SLIDINGWINDOW:4:15 MINLEN:36”) (Walker et al. 2014; Wang et al. 2024) . The polished assembly was scaffolded into 20 pseudochromosomes via genome-wide synteny with the European *SP9509* reference genome using Chromosemble in Satsuma v2 (default parameters) (Grabherr et al. 2010; Ernst et al. 2025). The genome assembly completeness was assessed with BUSCO v5.4.3 using the viridiplantae_odb10 dataset (Simão et al. 2015). Liftoff v1.6.3, a ‘lift over’ approach (Shumate and Salzberg 2021), was used to predict the coding genes in the *SP162* genome using the previously annotated *SP9509* genome (Ernst et al. 2025) . The parameters were “-copies -sc 0.90 -exclude_partial -polish” and other default settings.

**Design, cloning and analysis of guide sequences for CRISPR/Cas9 and their targets.**

Guide sequences for cloning sgRNAs targeting *SpZMET* were designed using CRISPOR V5.2 (https://crispor.gi.ucsc.edu; https://github.com/maximilianh/crisporWebsite) (Concordet and Haeussler 2018) run locally. The coding sequence of the four first exons of *SpZMET* were used as target genomic sequence and the NGG PAM sequence (20bp-NGG) was chosen for guide design. Resulting candidate guide target sequences were further filtered by: Efficiency/On-target score >0.6, first targeted codon starts with A/G, 50-70% CG content, no homeopolymeric regions of more than 3 constitutive bases were allowed, no termination in TC dinucleotide. The filter list of targets was further check for off-targets within the genome using a locally installed version of Cas-OFFinder (http://www.rgenome.net/cas-offinder/) (Bae et al. 2014). Selected sequences were cloned into sgRNA vectors as described in above.

**Analysis of CRISPR/Cas9 editing.**

Genomic DNA was extracted using the DNeasy Plant mini kit (Qiagen #69104) from 100 mg of flash-frozen tissues in liquid nitrogen mixed with 100 µL of 1mm ø glass beads (Carl-ROTH #A554.1) in a 1.5 mL microcentrifuge tube and ground to a fine powder on a Silamat S6 (Ivoclar) twice for 3 s. The genomic region containing sgRNA target sites was PCR amplified using primers listed in **Method S9**. PCR products were separated by agarose gel electrophoresis, gel-purified and cloned using CloneJET PCR Cloning Kit (Thermo Fisher, #K1231). Single colonies were selected, and plasmid DNA was extracted and sequenced. The sequences were analyzed using CLC Main Workbench 25 (Qiagen Digitals).

**Image analysis for phenotyping.**

3-4 fronds (including attached daughter fronds) were inoculated in 94 mm ø petri dishes with liquid SH-medium and image immediately. Cultures were imaged at regular intervals (every two days). Images were size and pixel normalized (72 pixels/inch) in Adobe Photoshop using the diameter of the plate at the surface of the media as reference. The normalized images were use for frond length measurements using Fiji (V2.16.0; https://imagej.net/software/fiji/)(Schindelin et al. 2012). For quantification of growth parameters, the same images were used to identify total, live or senescent frond areas using Ilastik (v1.4.1; https://www.ilastik.org), a supervised machine-learning software (Sommer et al. 2011), suitable for duckweed image analysis (Romano et al. 2022). First, pixel classification probabilities for green/red, senescent and background/medium were trained on images containing both types of tissues and then applied in bulk to all images using standard parameters following software user guidelines. The resulting pixel probabilities were used for bulk object classification with default parameters to obtain object areas for life and senescent tissue used for further analysis. In addition, object classification results were export as image

(.tiff) files for validation and further manual curation of the predictions when needed. Output images were auto-contrasted in Photoshop to increase the visibility of predicted areas. They were further pseudo-colored and merged using Fiji.

**Southern blot analysis**

DNA blots were performed following standard procedures (Brown 1993) . In brief, genomic DNA was extracted from 100 mg of flash-frozen, ground *Spirodela* fronds or calli using Dneasy Plant mini kit (Qiagen, #69104) following manufacturer's instructions. A total of 3-5 µg of genomic DNA was digested overnight with *XhoI* (Thermo Scientific, #FD0694), separated by gel electrophoresis on a 1% agarose gel, transferred by capilarity to Neutral Nylon membrane (GVS, #1213403) and cross-linked by UV irradiation with Stratalinker UV crosslinker 1800 (Stratagene) twice with 1,2 x $10^5$ µJoules. Probes were produced from PCR templates, radiolabelled with [α-32P]-dCTP (Hartmann Analytic, #SRP-305) using Prime-it II Random Primer Labelling kit (Agilent, #300385). Hybridization was carried out using PerfectHyb™ Plus Hybridization Buffer (Sigma Aldrich, #H7033). The signal was detected on a Phosphor screen (BAS-IP SR 2025; Sigma-Aldrich #GE-28-2564-78) and read with an Amersham Typhoon imaging system (Cytivia) after over-night exposure.

**Small RNA blot analysis.**

Total RNA was purified as described above. 5 µg of RNA were mixed with an equal volume of 2X NovexTM TBE-Urea sample buffer (ThermoFisher, #LLC6876), separated on a 17,5% polyacrylamide-urea gel (National Diagnostics #EC-833), transferred by electroblotting onto Neutral Nylon membrane (GVS, #1213403) and crosslinked chemically (12.5M 1-methylimidazol (Merk, #M50834), 31.25mg/mL N-(3-dimethylaminopropyl)-N'-ethylcarbodiimide hydrochloride (Merk, #E7750) as described in (Pall and Hamilton 2008). PCR fragment probes were radiolabelled with [α-32P]-dCTP (Hartmann Analytic, #SRP-305) using Prime-it II Random Primer Labelling kit (Agilent, #300385) and oligonucleotides used as a probe were radiolabelled with [$\gamma$-32P]-ATP (Hartmann analytic #SRP-501), using T4 Polynucleotide kinase (Thermofisher, #EK0031). Hybridization was carried out using PerfectHyb™ Plus Hybridization Buffer (Sigma Aldrich, #H7033) and membranes were washed with 2XSSC, 0.1%SDS. The signal was detected on a Phosphor screen (BAS-IP SR 2025; Sigma-Aldrich #GE-28-2564-78) and read with an Amersham Typhoon imaging system (Cytivia) after over-night exposure. Probes and oligonucleotides used are listed **Method S9**.

**Protein extraction and western blot.**

Protein extraction: 100mg of material of Spirodela fronds or calli was flash-frozen upon harvesting and ground with a Silamat S6.The ground tissue was mixed with 500µl of extraction buffer (0.7 M sucrose, 0.5 M Tris-HCl, pH 8, 5 mM EDTA, pH 8, 0.1 M NaCl, 2% b-mercaptoethanol) supplemented with cOmpleteTM mini EDTA-free protease inhibitor cocktail (Roche, #04693159001). Then, 1 volume of TRIS-buffered phenol (pH 8) was added and the mix was homogenyzed for 5 minutes. After centrifugation (12000g for 10 min at 4ºC), the upper phenolic phase was recovered, and proteins were precipitated methanol at -20C overnight using 5 volumes of 0.1 ammonium acetate dissolved in methanol. After pelleting by centrifugation (5000g for 15 min at 4ºC) and washing twice with 0,1 M ammonium acetate dissolved in methanol, the protein pellet was resuspended in 50-100 µL of resuspension buffer (3% SDS, 62,3 mM Tris-HCl, pH 8, 10% glycerol).

Protein blots: Western blot was performed following standard denaturing SDS-PAGE and blotting procedures (Sambrook and Russell 2001). 150 to 300 µg of protein were resolved on SDS-polyacrilamide gels and transferred onto Inmobilon-P PVDF membrane (Millipore, # IPVH00010). Membranes were blocked and incubated with the appropriate primary antibodies in PBS with 0.1% Tween-20 and 5% non-fat dried milk. Primary antibody was incubated overnight at 4ºC, followed by incubation with HRP-conjugated secondary antibody for 1 hour at room temperature. Detection was performed using clarity Max™ Western ECL Substrate (Biorad, #1705062), and signal was imaged using the ChemiDoc Imaging system (Bio-Rad). Following detection, protein loading was verified by coomassie staining of the membranes.

**Genome size estimation by flow cytometry.**

DNA content of *S. polyrhiza* WT and regenerated lines was estimated by flow cytometry according to the stablished methodology for plants (Doležel et al. 2007). *L. minor* 8623 was used as standard with a known DNA content of 1C = 0.4182 pg (Hoang et al. 2019). 25 mg of *S. polyrhiza* or *L. minor* fronds, alone or mixed, were disrupted by chopping with a double-edged razor blade (Wilkinson sword) in a 94 mm petri dish containing 100 µL of nuclei isolation buffer (NIB) [0.1 mM citric acid, 0.5% (v/v) Triton-X 100, pH1.5]. After chopping, 450 µL of NIB were slowly added and the resulting suspension was filtered through a 35 µm pore nylon cell strainer cap tube (Corning, #352235). RNase A (Thermo-Scientific, #EN0531) was added to a final 0.15 mg/mL concentration and incubated in a water bath at 37ºC for 30 min. Nuclei were stained by adding 2 mL of 6 mg/mL pH 9,5 Propidium iodide solution (Merck, #P4170-25MG) and incubated for 16h at 4ºC in darkness. DNA content measurements were performed using a FACS LSR II Fortessa (BD Biosicences). Absolute DNA contents were calculated based on the average intensity values of G1 peak of the sample relative to the standard. Each line was measured in three independent biological replicates.

##### Bacteria titers

Fronds infiltrated with *Agrobacterium* were surface sterilized in 40 mL 70% ethanol and rinsed twice in an equal volume of $ddH_2O$. To extract bacteria, individual fronds were transferred to 1.5 mL microcentrifuge tubes containing 200 µL 1X PBS buffer and manually disrupted using a pellet-pestle (Merck, #BAF199230001) and incubated at room temperature with gentle agitation for 10 min. Tissue debris was spun down at 5000*g* for 1 min and 50 µL were transferred to 96-well plates where 1:10 serial dilutions were performed. 5 µL droplets from each dilution were deposited on LB-agar plates containing the appropriate antibiotics for either non-transformed *Agrobacterium* EHA105 (50 µg/mL rifampicin, 50 µg/mL streptomycin) or cells transformed with the T-DNA vector (50 µg/mL rifampicin, 50 µg/mL streptomycin, 50 µg/mL spectinomycin). Plates were let to dry and incubated at 28ºC for two days. Colonies in in the last dilution with more than one colony (if possible) were counted to calculate colony forming units (cfu) per frond.

##### *S. polyrhiza* 9509 TAS3 siRNAs analysis.

*S. polyrhiza* 9509 genome and mapped small RNA libraries were obtained from (doi:10.5281/zenodo.10911532.) (Dombey et al. 2025) and loaded on locally installed JBrowse 2 (V3.0.3, https://jbrowse.org/jb2/) (Diesh et al. 2023). *S. polyrhiza* 9509 miR390 was identified in through sequence homology search in publicly available *S. polyrhiza* 9509 small RNA libraries (PRJNA1164696) using the *Arabidopsis* miR390 as bait. The small RNA coverage and sequence of the region corresponding to the location of previously identified putative *TAS3* locus were extracted for visualization and identification of the miR390 target sites by sequence alignment using the CLC software.

### SUPPLEMENTAL METHOD S9: List of Oligonucelotides.

#### Genotyping CRISPR/Cas9 @ SpZMET

| Name | Description | sequence (5' -> 3') | Use |
|---|---|---|---|
| CDo0075 | CDO0075_PfSpZ1 | CAGAAGCAAAAGGACTGCCG | used as Forward primer for Sanger sequencing of Zmet crispr mutant lines |
| CD00076 | CDO0076_PrSpZ1 | AAACGTCAACTTACCTGGAC | used as Reverse primer for Sanger sequencing of Zmet crispr mutant lines |

#### Cloning of SpZMET gides

| Name | Description | sequence (5' -> 3') | Use |
|---|---|---|---|
| CDo0025 | CD25SpZMET G1PF | GTTGCAAGGGTCCACCCTTCCGTT | oligo forward used to clone sgRNA3 into pOsU6b_gRNAscf |
| CDo0026 | CD26SpZMET G1PR | AAACAACGGAAGGGTGGACCCTTG | oligo reverse used to clone sgRNA3 into pOsU6b_gRNAscf |
| CDo0027 | CD27SpZMET G2PF | GGCAGGCGAAATGCCACTACCGC | oligo forward used to clone sgRNA4 into pOsU3m_gRNAscf |
| CDo0028 | CD28SpZMET G2PR | AAACGCGGTAGTGGCATTTCGCC | oligo reverse used to clone sgRNA4 into pOsU3m_gRNAscf |
| CDo0029 | CD29SpZMET G3PF | GCCGTCGTTCGGCGTCGCAGAAG | oligo forward used to clone sgRNA1 into pOsU6a/lacz_sgRNAscf |
| CDo0030 | CD30SpZMET G3PR | AAACCTTCTGCGACGCCGAACGA | oligo reverse used to clone sgRNA1 into pOsU6a/lacz_sgRNAscf |
| CDo0031 | CD31SpZMET G4PF | TCAGATCTTCCTTCCGCTTCCTC | oligo forward used to clone sgRNA2 into pOsU6c_sgRNAscf |
| CDo0032 | CD32SpZMET G4PR | AAACGAGGAAGCGGAAGGAAGAT | oligo reverse used to clone sgRNA2 into pOsU6c_sgRNAscf |

#### Amplification U3/6promoter-sgRNA-SGRNA scafold  for Golden Gate in destination plasmid pZmUBQ_zCas9i_P2A_GFP

| Name | Description | sequence (5' -> 3') | Use |
|---|---|---|---|
| VBo0149 | V149Pps-GGL(NoSpeI) | TTCAGAggtctcTctcgtAcTGGAATCGGCAGCAAAGG | oligo forward used to amplify the cassette pOsU3m_sgRNA4_sgRNAscf for cloning by GoldenGate into pZmUBQ_zCas9i_P2A_GFP |
| Co0114 | Pgs-2 | AGCGTGGGTCTCGTCAGGGTCCATCCACTCCAAGCTC | Xingliang Ma et al, Molecular Plant 2015. oligo reverese used to amplify the cassette pOsU3m_sgRNA4_sgRNAscf for cloning by GoldenGate into pZmUBQ_zCas9i_P2A_GFP |
| Co0115 | Pps-2 | TTCAGAGGTCTCTCTGACACTGGAATCGGCAGCAAAGG | Xingliang Ma et al, Molecular Plant 2015. oligo forward used to amplify the cassette pOsU6alacz_sgRNA1_sgRNAscf for cloning by GoldenGate into pZmUBQ_zCas9i_P2A_GFP |
| Co0116 | Pgs-3 | AGCGTG GGTCTCGTCTTCACTCCATCCACTCCAAGCTC | Xingliang Ma et al, Molecular Plant 2015. oligo reverese used to amplify the cassettepOsU6alacz_sgRNA1_sgRNAscf for cloning by GoldenGate into pZmUBQ_zCas9i_P2A_GFP |
| Co0117 | Pps-3 | TTCAGAGGTCTCTAAGACTTTGGAATCGGCAGCAAAGG | Xingliang Ma et al, Molecular Plant 2015. oligo forward used to amplify the cassette pOsU6b_sgRNA3_sgRNAscf for cloning by GoldenGate into pZmUBQ_zCas9i_P2A_GFP |
| Co0118 | Pgs-4 | AGCGTGGGTCTCGAGTCCTTTCCATCCACTCCAAGCTC | Xingliang Ma et al, Molecular Plant 2015. oligo reverese used to amplify the cassette pOsU6b_sgRNA3_sgRNAscf for cloning by GoldenGate into pZmUBQ_zCas9i_P2A_GFP |
| Co0119 | Pps-4 | TTCAGAGGTCTCTGACTACATGGAATCGGCAGCAAAGG | Xingliang Ma et al, Molecular Plant 2015. oligo forward used to amplify the cassette pOsU6c_sgRNA2_sgRNAscf for cloning by GoldenGate into pZmUBQ_zCas9i_P2A_GFP |
| VBo0150 | V150Pgs-GGR(NoMluI) | AttaTtggtctcGaccgtAcTCCATCCACTCCAAGCTC | oligo reverese used to amplify the cassette pOsU6c_sgRNA2_sgRNAscf for cloning by GoldenGate into pZmUBQ_zCas9i_P2A_GFP |

#### oligo-probes and primers used to amplify DNA templates used as probes

| Name | Description | sequence (5' -> 3') | Use |
|---|---|---|---|
| Co0132 | ZmUBQ probe PF | GGTCGTGCCCCTCTCTAG | oligo forward used to amplify part of the ZMUBQpromoter used as probe for Southern blot |
| Co0133 | ZmUBQ probe PR | GTCCAGAGGCAGCGACAG | oligo reverse used to amplify part of the ZMUBQpromoter used as probe for Southern blot |
| VBo0358 | VB358 eGFPhpF1F | GGAGGACGGCAACATCCTGGGGCAC | oligo forward used to amplify the last 322bp of eGFP, used as probe for Low Molecular Weight Northern Blot |
| VBo0359 | VB359 eGFPhpF1R | CTTGTACAGCTCGTCCATGCCGTG | oligo reverse used to amplify the last 322bp of eGFP, used as probe for Low Molecular Weight Northern Blot |
| Co0225 | TAS3 F | AGAGATTATCCATCGGTGTTACT | oligo forward used to amplify part of the TAS3 locus in Spirodela polyrhiza, used as probe for Low Molecular Weight Northern Blot |
| Co0226 | TAS3 R | CGACAATTCGATCAACCTTACA | oligo reverse used to amplify part of the TAS3 locus in Spirodela polyrhiza, used as probe for Low Molecular Weight Northern Blot |
| Co0138 | mir159 | TAGAGCTCCCTTCAATCCAAA | oligo used as probe for detection of mir159 in Low Molecular Weight Northern Blot |
| Co0062 | snoRNA_U6 | AGGGGCCATGCTAATCTTCTC | oligo used as probe for detection of U6 in Low Molecular Weight Northern Blot |

#### qPCR

| Name | Description | sequence (5' -> 3') | Use |
|---|---|---|---|
| CDo0077 | eGFP PF1 | ATCATGGCCGACAAGCAGAAG | oligo forward used in qPCR to amplify part of the eGFP sequence |
| CDo0078 | eGFP PR1 | TCTCGTTGGGGTCTTTGCTC | oligo reverse used in qPCR to amplify part of the eGFP sequence |
| qo0039 | SpActin PF2 | CTCTCTCTATGCCAGTGGTCG | oligo forward used in qPCR to amplify part of the actin cDNA sequence |
| qo0044 | SpActin PR3 | TCGTAGATCGGAACCGTGTG | oligo reverse used in qPCR to amplify part of the actin cDNA sequence |
